# High-throughput competitive binding assay for targeting RNA with small molecules: discovery of new PreQ_1_ riboswitch ligands

**DOI:** 10.1101/2024.05.22.595132

**Authors:** Sophie E.L. Wintermans, Jana S. Hoffmann, Mariska D. Tacoma, Indy Broekhuizen, Bjorn R. van Doodewaerd, Paul P. Geurink, Antonius P.A. Janssen, Marta Artola, René R.C.L. Olsthoorn

## Abstract

In the evolving landscape of RNA targeting, there is an indisputable need for new screening methodologies to find small molecules targeting relevant tertiary RNA structures, like viral pseudoknots or bacterial riboswitches. Here, we developed a competitive binding high-throughput screening assay to identify ligands for the bacterial PreQ_1_-I riboswitch. In this assay, ligands compete with a rationally designed quencher-labeled antisense for binding to the riboswitch. The method is validated for the *Fusobacterium nucleatum* (*Fnu*), *Thermoanaerobacter tengcongensis* (*Tte*), *Bacillus subtilis* (*Bsu*) and *Enterococcus faecalis* (*Efa*) PreQ_1_ riboswitches, using the natural riboswitch ligand PreQ1 and various analogues. A commercial RNA-focused library consisting of ∼15,000 compounds was then screened against the *Fnu* riboswitch, leading to the identification of 4 hits exhibiting competitive binding activity. These hits were evaluated in in vitro translation assays against several PreQ_1_ riboswitches. The most promising hit **4494** showed competitive binding activity to the *Fnu*, *Tte*, *Bsu* and *Efa* riboswitches, and was able to inhibit translation of a *Tte* riboswitch-regulated reporter gene, making it an interesting starting point for the development of new antibiotics. In essence, this HTS assay has the potential to discover highly sought-after RNA targeting small molecules for complex and clinically relevant RNA structures.

**Graphical abstract:** 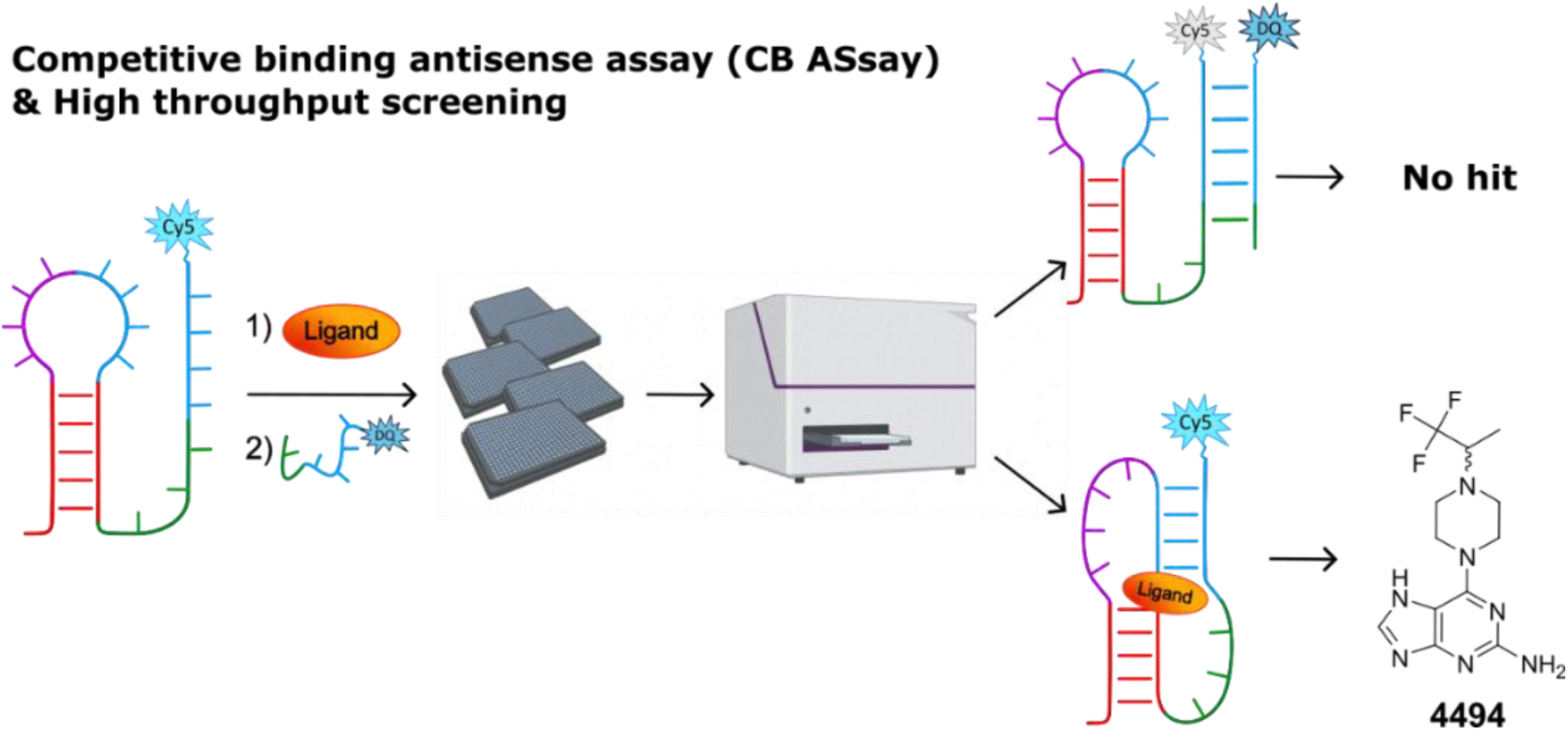

## Introduction

Targeting RNA with small-molecule drugs is an emerging research field in medicinal chemistry. However, some of the key challenges in targeting tertiary RNA structures include identifying relevant RNA targets and having access to effective high throughput screening assays. Promising and recently discovered RNA targets are bacterial riboswitches^1^. These RNA structures are often found in the 5’untranslated region (5’-UTR) of bacterial messenger RNA (mRNA) and change conformation upon binding of a ligand (*e.g.* metabolites, co-enzymes or ions) leading to the alteration of gene expression^2–5^. Riboswitches consist of a ligand-binding aptamer domain and an expression platform that exerts genetic control, often over genes that are involved in metabolic pathways^6^. In nature, metabolites are able to bind to RNA with an affinity and selectivity that is comparable to ligand-protein interactions^2^, and riboswitches can have distinct small-molecule binding pockets^5^, making them interesting targets for medicinal chemistry research. However, current drug development for such tertiary structures suffers from a lack of effective high throughput screening assays.

In this research, a high throughput screening is developed for the PreQ_1_-I riboswitch, which is the smallest riboswitch known to date and is present in several clinically relevant human pathogenic bacteria, such as *Bacillus anthracis*, *Clostridium difficile*, *Enterococcus faecalis*, *Haemophilus influenzae* and *Neisseria meningitidis*^7–11^. The PreQ_1_ riboswitch is involved in the biosynthesis of the hypermodified queuosine (Q) nucleoside (Figure 1A). Queuosine occupies the anticodon wobble position of several tRNAs (tRNA^Tyr^, tRNA^Asp^, tRNA^His^ and tRNA^Asn^)^12,13^ and plays an important role in the correct decoding of the corresponding codons^14,15^. Q has shown to be important for viability and virulence of various bacteria^14,16,17^. In particular, the PreQ_1_ riboswitch is known to directly regulate the expression of four enzymes (QueCDEF) involved in the biosynthesis of PreQ_1_ (a key precursor of Q) and a PreQ_1_ transporter protein^8,18–20^. By binding to PreQ_1_, the riboswitch either prematurely stops transcription by forming a terminator hairpin (Figure 1B), or prevents translation by sequestering the Shine-Dalgarno (SD) sequence (Figure 1C)^9,21^.

**Figure 1.**
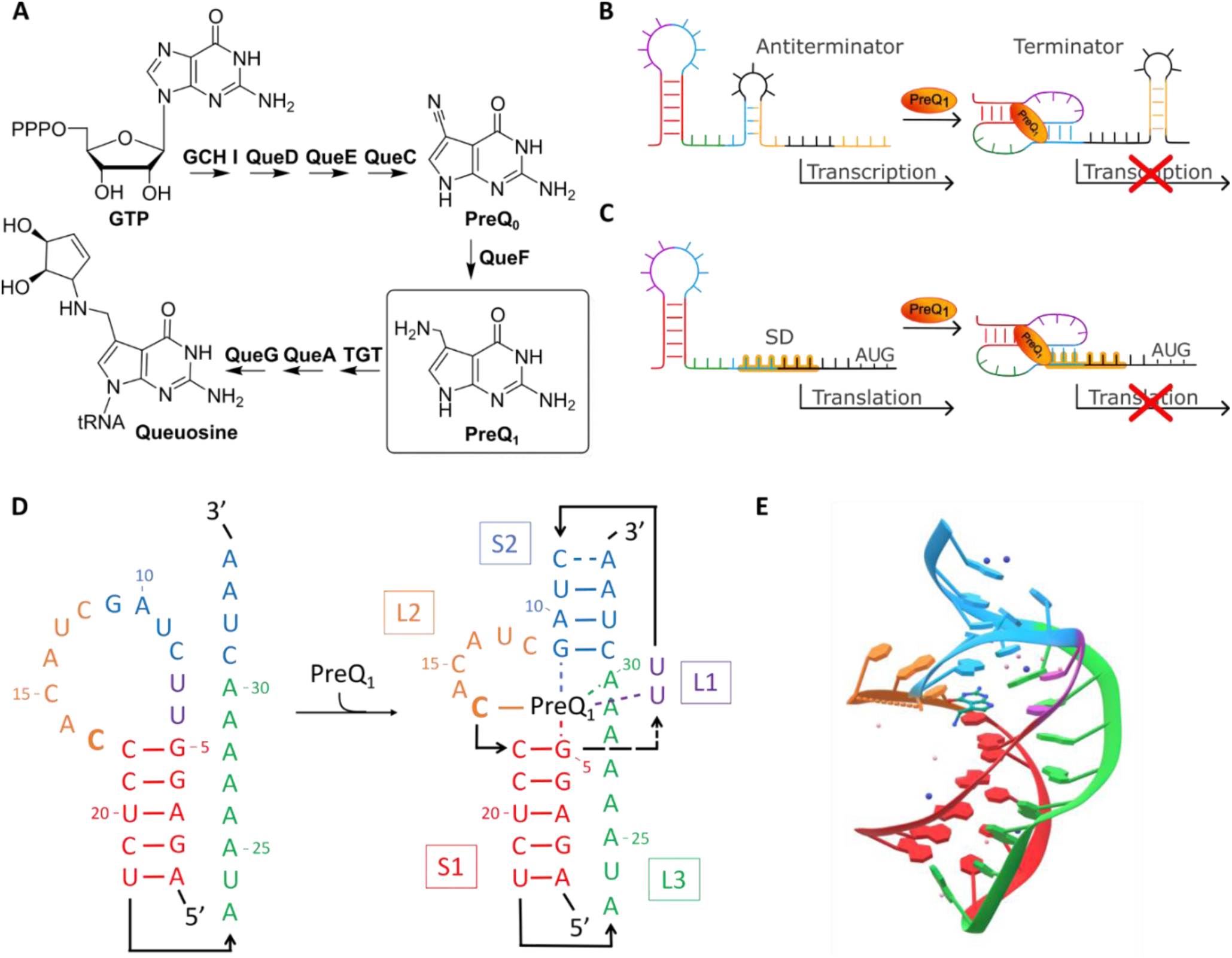
**A**) Biosynthetic pathway of queuosine from GTP: enzymes involved and chemical structures of key intermediates PreQ_0_ and PreQ_1_ are shown. **B**) PreQ_1_ riboswitch gene regulation: termination of transcription. Binding of PreQ_1_ to the riboswitch leads to the formation of a terminator hairpin, thereby prematurely terminating transcription. **C**) PreQ_1_ riboswitch gene regulation: prevention of translation. Binding of PreQ_1_ to the riboswitch leads to the sequestration of the Shine-Dalgarno (SD) sequence, thereby preventing ribosomes from binding the mRNA and blocking translation. **D**) 2D structure of the *Bacillus subtilis* (*Bsu*) PreQ_1_ riboswitch. Interactions of PreQ_1_ with several bases of the riboswitch are shown as dotted lines. **E**) 3D structure of the *Bsu* PreQ_1_ riboswitch bound to PreQ_1_ (PBD: 3FU2).

When bound to PreQ_1_, the PreQ_1_-I riboswitch forms a hairpin- or H-type pseudoknot (PK) (Figure 1D and 1E)^22–24^. Stems S1 and S2 are connected by single-stranded loops (L1, L2 and L3) that can form tertiary interactions such as base triples with base pairs in the stem or interact with ligands to further stabilize the PK. PreQ_1_ aptamers are able to bind PreQ_1_ with nanomolar affinities and are able to distinguish between closely related purines, as Roth *et al.* showed for several PreQ_1_ analogues binding to the *Bacillus subtilis* (*Bsu*) aptamer^8,25,26^. Tight PreQ_1_ binding is mainly driven by hydrogen bonding and stacking interactions^23^. Specifically, out of the ten potential hydrogen-bonding groups of PreQ_1_, nine are recognized by the riboswitch^8^. Notably, L2 contains a highly conserved cytidine (C17) that is essential for PreQ_1_ binding as they form a Watson-Crick-Franklin (WCF) base pair. Close analogues of PreQ_1_, such as PreQ_0_ and guanine, have several fold lower affinities for the riboswitch, demonstrating that other functional groups of PreQ_1_, including the aminomethyl group, are also very relevant for ligand recognition.

Development of synthetic ligands for the PreQ_1_ riboswitch able to suppress the production of proteins essential for Q synthesis and transport is of great interest for developing new antibiotics. Several PreQ_1_ analogues and synthetic ligands for the PreQ_1_ riboswitch have been discovered and investigated^27–30^, some of which are able to mimic PreQ_1_ function in *in vitro* and *in vivo* riboswitch assays. However, the discovery of new structurally diverse ligands of the PreQ_1_ riboswitch remains an interesting field of research.

Previous research has shown that competitive displacement of specific (fluorescent) binders, such as fluorescent intercalators^31^, the Tat peptide from the HIV-1-TAR hairpin^32–35^ and labeled antisense RNA^36^, is an effective strategy for identifying small-molecule ligands for relatively simple RNA structures. In this work, a high throughput competitive binding (CB) antisense assay (ASsay) was developed to identify small-molecule ligands that bind to the PreQ_1_ riboswitch (Figure 2), thereby demonstrating that competitive binding assays are also applicable to more complex (and clinically relevant) RNA structures. The method employs a 3’-Cy5-labeled pseudoknot (Cy5-PK) and a 5’-dark quencher-labeled AS (DQ-AS) that is complementary to the bases at the 3’-end of the PK (comprising S2 and a part of L3 of the PK), as the 3’-end is the link between the aptamer and expression platform. When the Cy5-PK and DQ-AS are hybridized, the FRET pair is brought into proximity and Cy5 is quenched, resulting in a drop in Cy5 signal. Addition of a PK-binding ligand can stabilize the Cy5-PK, thereby preventing DQ-AS binding, which results in an unchanged fluorescent signal.

**Figure 2.**
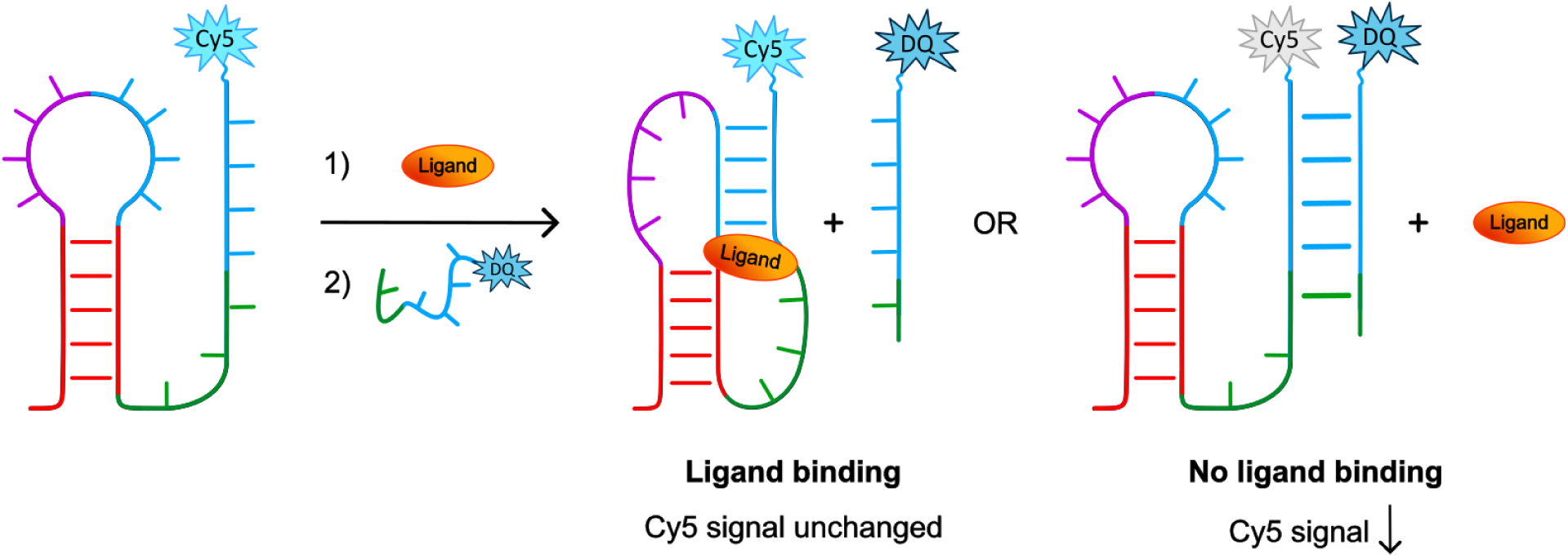
Schematic overview of the here developed competitive binding ASsay. A Cy5-labeled pseudoknot (Cy5-PK) is incubated with a potential ligand, after which a dark quencher labeled antisense (DQ-AS) is added. If a small molecule ligand binds to the PK, the PK is stabilized and hybridization with the DQ-AS is prevented. In contrast, if the ligand does not bind to the Cy5-PK, the DQ-AS hybridizes with the Cy5-PK and quenches Cy5 fluorescence signal.

The CB ASsay was validated by testing PreQ_1_ and several analogues on four different Cy5-labeled PreQ_1_-I riboswitches from *Fusobacterium nucleatum* (*Fnu*), *Thermoanaerobacter tengcongensis* (*Tte), Bacillus subtilis* (*Bsu*) and *Enterococcus faecalis* (*Efa*). Subsequently, a HTS was performed and four *Fnu* riboswitch-binding molecules were identified with the CB ASsay, which were also tested with the *Tte*, *Bsu* and *Efa* riboswitches. Hits were then evaluated in functional assays with PreQ_1_-I riboswitches from *Fnu*, *Bsu*, *Efa*, *Tte* and *Escherichia coli* (*Eco)*, and finally docked using the known crystal structures of the *Bsu* and *Tte* riboswitches to identify possible binding modes.

## Results

### Competitive binding assay development for PreQ_1_ riboswitch ligands

The competitive binding ASsay was developed using the *Fusobacterium nucleatum* (*Fnu*) PreQ_1_ riboswitch^24,37^, which is a well characterized and clinically relevant target^38,39^. The *Fnu* PK was first extended at the 3’-end with a d[CCTT] spacer and a terminal Cy5 label (Figure 3A). Next, various antisense lengths were evaluated for their ability to bind to the PK using native polyacrylamide gel electrophoresis (PAGE) (Figure S1). We postulated that the ideal antisense should exhibit approximately 80% binding affinity for the PK at a 1:1 ratio of PK to AS, thereby allowing sufficient competition with screened ligands while simultaneously ensuring a favorable signal-to-background ratio. Of all antisenses that fulfilled the 80% binding criterium, the shortest antisense (ANTI-*Fnu*-GT) was selected as it was the weakest competitor and thus suitable for screening weaker ligands. The antisense was subsequently extended at the 5’-end with a d[TT] spacer and an IowaBlack® RQ (IBRQ) dark quencher (DQ) label. As a control, a mutant *Fnu* PK was tested alongside the wildtype (WT) PK variant. These two PKs only differ at nucleotide position 17, having either a cytosine (WT) or uracil (mutant) (Figure 3B). In the CB ASsay, the PK and ligand were pre-incubated for 1 hour, followed by the addition of labeled AS and further incubation of additional 2 hours, after which the Cy5 fluorescence was measured. In all experiments, final PK and AS concentration were 50 nM. The CB ASsay was conducted at room temperature in a near-physiological buffer^26,40,41^ (100 mM Tris (pH =7.5), 100 mM KCl, 10 mM NaCl, 1 mM MgCl_2_, 0.1% (v/v) DMSO and 0.01% (v/v) Tween20). It is worth mentioning that the assay showed to be robust towards variations in concentration of individual buffer components (Figure S2). To determine the maximal Cy5 signal, a positive control was established by adding a 10-fold excess of unlabeled AS.

**Figure 3.**
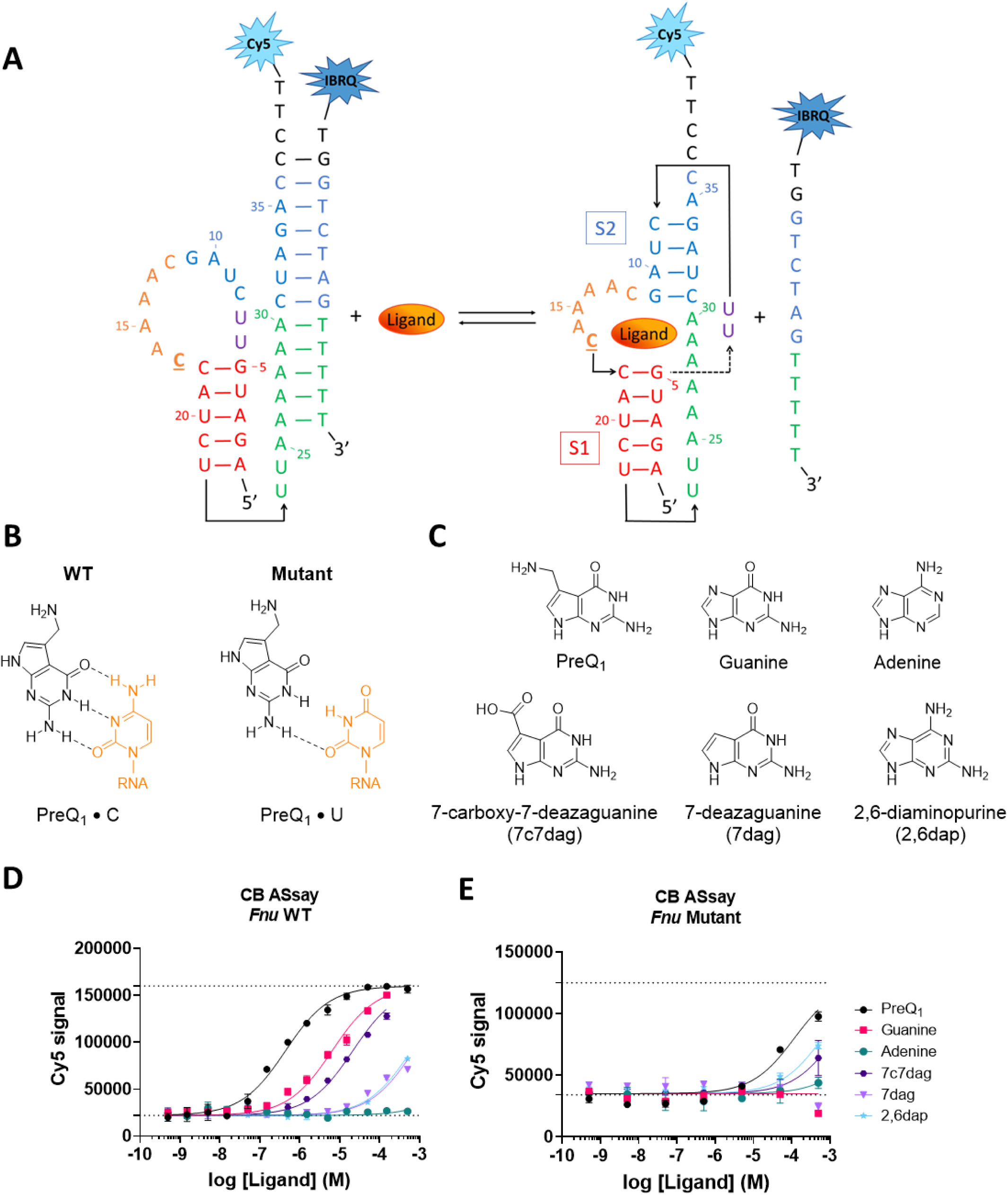
Competitive binding assay for the PreQ_1_ riboswitch. **A)** Schematic representation of the equilibrium between the labeled PreQ_1_ riboswitch of *Fusobacterium nucleatum* (*Fnu*) and labeled antisense (AS) with and without ligand binding. The essential nucleotide C17 is <u>underlined</u>. **B)** Base pairing of PreQ_1_ to C17 (WT) or C17U (mutant). **C)** Molecular structures of PreQ_1,_ guanine, adenine, 7-carboxy-7-deazaguanine (7c7dag), 7-deazaguanine (7dag) and 2,6-diaminopurine (2,6dap). **D-E)** Competitive binding assay of PreQ_1_ and its analogues for the *Fnu* WT PK (**D**), *Fnu* mutant PK (**E**). Dotted lines denote the minimum and maximum Cy5 signal as determined by the negative (DMSO) and positive (10x excess unlabeled AS) controls.

To validate the assay, PreQ_1_ and various analogues (guanine, adenine, 7-deazaguanine (7dag), 7-carboxy-7-deazaguanine (7c7dag) and 2,6-diaminopurine (2,6dap), Figure 3C) were tested in the competitive binding assay for their ability to prevent AS binding to the WT riboswitch and retain Cy5 signal (Figure 3D). Upon addition of PK-stabilizing ligands, the Cy5 signal was indeed preserved, indicating that the ligands are able to compete with the AS.

For every PreQ_1_ analogue, the concentration at which the ligand competed with 50% of AS-PK binding (apparent half-maximal effective concentration, EC_50_) was determined (Table 1). In line with literature^8^, PreQ_1_ is the most potent ligand, with an EC_50_ of 0.44 ± 0.07 µM. Close analogues that lack or have a modified aminomethyl group, such as guanine (EC_50_ = 6.9 ± 0.7 µM), 7c7dag (EC_50_ = 24 ± 3 µM) and 7dag (EC_50_ = >500 µM), showed a lower binding affinity for the *Fnu* riboswitch, as also observed for *Bsu*^8^. Analogues that lack both the aminomethyl group and have modifications in the WCF face, including adenine and a key exocyclic amine in the WCF face crucial for ligand recognition^8^, retained some affinity for the *Fnu* PreQ_1_-PK (EC_50_ = 609 ± 109 µM).

**Table 1.** Overview of EC_50_ values from the CB ASsay and apparent *K*_D_ values of PreQ_1_ and its analogues, as determined for the *Fusobacterium nucleatum* (*Fnu*), *Thermoanaerobacter tengcongensis* (*Tte*), *Bacillus subtilis* (*Bsu*) and *Enterococcus faecalis* (*Efa*) WT and *Fnu* C17U mutant PreQ_1_ riboswitches. ndb = no detectable binding

|  | *Fnu* |  |  | *Tte* | *Bsu* | *Efa* |
| --- | --- | --- | --- | --- | --- | --- |
|  | WT |  | Mutant | WT | WT | WT |
| | EC<sub>50</sub> (μM)<br>CB ASsay | $K_D$ (μM)<br>MST | EC<sub>50</sub> (μM)<br>CB ASsay | EC<sub>50</sub> (μM)<br>CB ASsay | EC<sub>50</sub> (μM)<br>CB ASsay | EC<sub>50</sub> (μM)<br>CB ASsay |
| **PreQ<sub>1</sub>** | 0.44 ± 0.07 | 0.36 ± 0.02 | 122 ± 77 | 0.009 ± 0.002 | 2.1 ± 0.3 | 6.1 ± 0.8 |
| **Guanine** | 6.9 ± 0.7 | 12 ± 2 | ndb | 0.33 ± 0.1 | 220 ± 69 | 568 ± 53 |
| **Adenine** | ndb | >500 | >500 | ndb | ndb | ndb |
| **7c7dag** | 24 ± 3 | 11 ± 1 | >500 | 0.38 ± 0.1 | >500 | 332 ± 31 |
| **7dag** | >500 | ndb | ndb | 2.9 ± 0.9 | ndb | ndb |
| **2,6dap** | 609 ± 109 | 610 ± 127 | >500 | 30 ± 8 | >500 | ndb |

The observed trend in ligand activity with this new CB ASsay was further validated through the assessment of their apparent binding affinities to the *Fnu* WT PreQ_1_ riboswitch using microscale thermophoresis (MST) (Table 1 and Figure S3). Remarkably, the determined apparent *K*_D_ and EC_50_ values exhibit excellent agreement, differing by only a factor of two. Noteworthy is the higher sensitivity of the CB ASsay to weak-binding ligands, particularly evident with 7dag, surpassing the observed sensitivity of MST. For PreQ_1_, both the apparent *K*_D_ and EC_50_ values closely align with the reported apparent *K*_D_ of 0.28 ± 0.05 µM in the literature^37^.

To demonstrate that the retention of Cy5 signal in the competitive binding assay is driven by ligand occupancy, PreQ_1_ and its analogues were additionally tested using the C17U mutant PK (Figure 3B, 3E and Table 1). The substitution of the cytosine to an uracil reduces PreQ_1_ affinity, due to the disruption of the WCF base pairing^8^. As expected, PreQ_1_ and other guanine analogues showed a reduced activity with the mutant PK, whereas adenine analogues (adenine and 2,6dap) exhibited similar or increased activity. Additionally, the competitive binding between the AS and PreQ_1_, guanine and 7c7dag was further validated with native PAGE (Figure S4), where upon increasing ligand concentration, a clear decrease in AS-PK complex and increase of PK conformation was observed.

### Expanding the CB ASsay to other PreQ_1_ riboswitches

Encouraged by these results, the CB ASsay was extended to the well-studied *Thermoanaerobacter tengconensis* (*Tte*), *Bacillus subtilis* (*Bsu*) and clinically relevant *Enterococcus faecalis* (*Efa*) PreQ_1_ riboswitches. In contrast to the *Fnu* pseudoknot, the second stems of *Tte*, *Bsu* and *Efa* PKs are partially stabilized by non-WCF bps, introducing greater complexity to the design of AS sequences. Fortunately, AS selection for the *Tte* riboswitch (Figure S5) was similar to that of *Fnu*, with the PK and PK-AS complexes separating clearly on gel and showing length- and linker-dependent binding behavior. Antisense ANTI-*Tte*-GT3 was selected as it fulfilled the previously postulated 80% binding criterium that worked well for *Fnu*. In contrast, AS selection for *Bsu* and *Efa* proved more difficult, as the PK and PK-AS complexes could not be easily separated on native gel. To overcome this issue, unlabeled antisenses of varying lengths were screened by co-incubating them with Cy5-labeled PKs and increasing concentrations of PreQ_1_, and their competitive binding was visualized using native PAGE (Figure S6 and S7). In the absence of PreQ_1_, all PK was bound to the AS, presumably due to the lower stability of stem S2 of *Bsu* and *Efa* compared to *Fnu*. Since none of the tested antisenses complied with the 80% binding criterium, the antisenses that showed a clear binding to the PK in the absence of PreQ_1_ and clear competitive binding upon addition of PreQ_1_ (ANTI-*Bsu*-GT1 and ANTI *Efa*-GT1) were selected.

All selected ASs were subsequently labeled with a 5’-terminal DQ, and PreQ_1_ and its analogues were tested in the corresponding CB ASsay (Figure 4) from which EC_50_ values were determined (Table 1). The *Tte* riboswitch, which has an apparent *K*_D_ of 2 nM for PreQ_1_^25^, indeed showed high affinity for PreQ_1_ and its analogues, reaching nanomolar EC_50_ values for PreQ_1_ and a similar trend in analogue activity as observed for *Fnu*. Notably, for both *Bsu* and *Efa*, the EC_50_ values of PreQ_1_ were an order of magnitude higher than those observed for *Fnu,* despite PreQ_1_ having a higher reported binding affinity to the *Bsu* riboswitch (*K*_D_ = 50 nM)^8^. This clearly underscores the influence of both the intrinsic stability of the RNA structure and the binding potency of the AS to the target on the obtained EC_50_ values. Therefore, it highlights the importance of appropriately tuning the signal window by adjusting factors such as antisense length, antisense complementarity and/or RNA nucleotide- or backbone modifications for each specific RNA target^36,42^.

**Figure 4.**
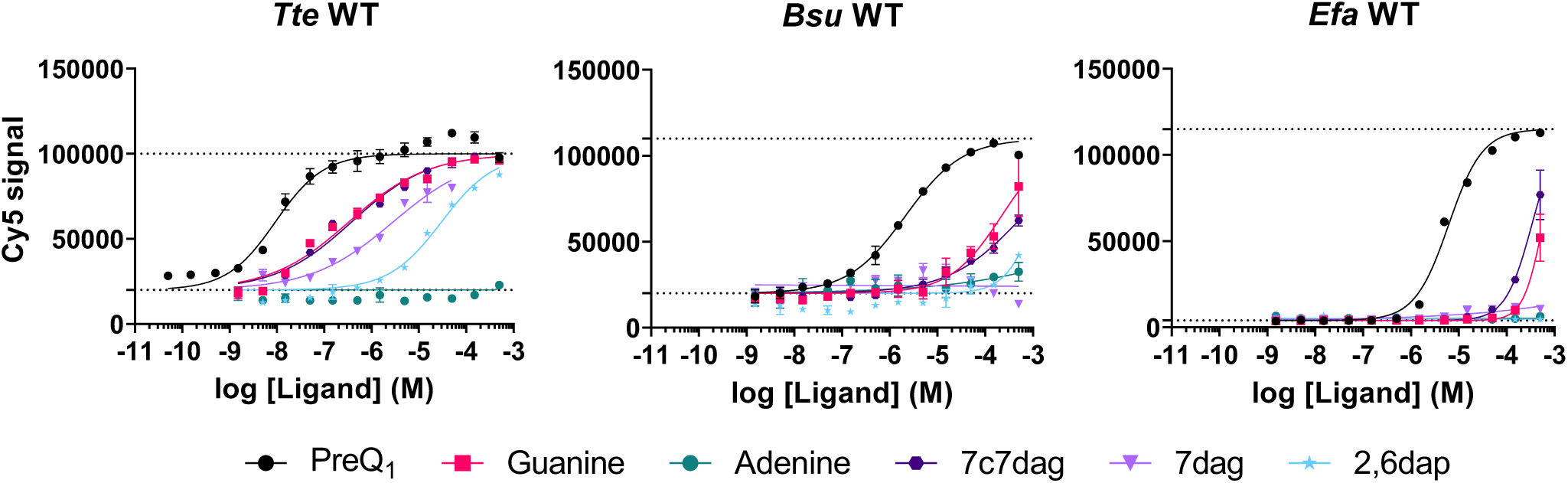
Competitive binding assay of PreQ_1_ and its analogues for the *Thermoanaerobacter tengcongensis* (*Tte*), *Bacillus subtilis* (*Bsu*), and *Enterococcus faecalis* (*Efa*) WT PK. Dotted lines denote the minimum and maximum Cy5 signal as determined by the positive and negative controls.

### High throughput screening (HTS) and hit confirmation

The CB ASsay was subsequently adapted into a HTS format for the discovery of new ligands targeting the *Fnu* PreQ_1_ riboswitch. A commercially available RNA-focused library from Enamine, comprising 15,520 RNA-binding compounds, was screened in duplicate at a final ligand concentration of 25 µM. Positive and negative controls were established using an excess of unlabeled AS (without quencher) and DMSO, respectively (Figure S8A-B). For additional quality control, 25 µM PreQ_1_ and guanine were screened alongside the compounds. The HTS data (see supporting Excel file) were analyzed with KNIME Analytics Platform (see Supporting KNIME files). Z’-scores were calculated for every plate and ranged from 0.764 to 0.969, with an average Z’-score of 0.815, indicating that the assay was of high quality with a good distinction between positive and negative signals.

Raw fluorescence data (Figure S8C-D) were normalized to account for plate, row and column, minimizing potential plate effects^43^. The resulting ‘B-scores’ of both screenings are shown in Figure 5A. Unfortunately, the ‘strongest’ hits from either the first or second screening did not overlap. Therefore, 107 initial hits were identified based on the following criteria: compounds exhibiting high activity (B-score >10) in one of the single screenings and compounds displaying moderate activity (B-score 5 - 10) in both screenings. The 107 initial hits, along with 136 of their most active analogues, underwent a duplicate screening at 100 µM (Figure 5B). To minimize the chance of false negatives, both the confirmed hits and initial hits were re-tested at 5, 25, 50 and 100 µM, resulting in 21 promising hits (Table S1, Figure S9). Among these, based on displayed activity and distinctiveness of the molecular structure, the 9 most promising compounds were selected for re-purchase and further evaluation. Preliminary CB ASsays revealed five false positives, with one showing no activity and four displaying nonspecific enhancement of Cy5 signal due to visible insolubility^44^.

**Figure 5.**
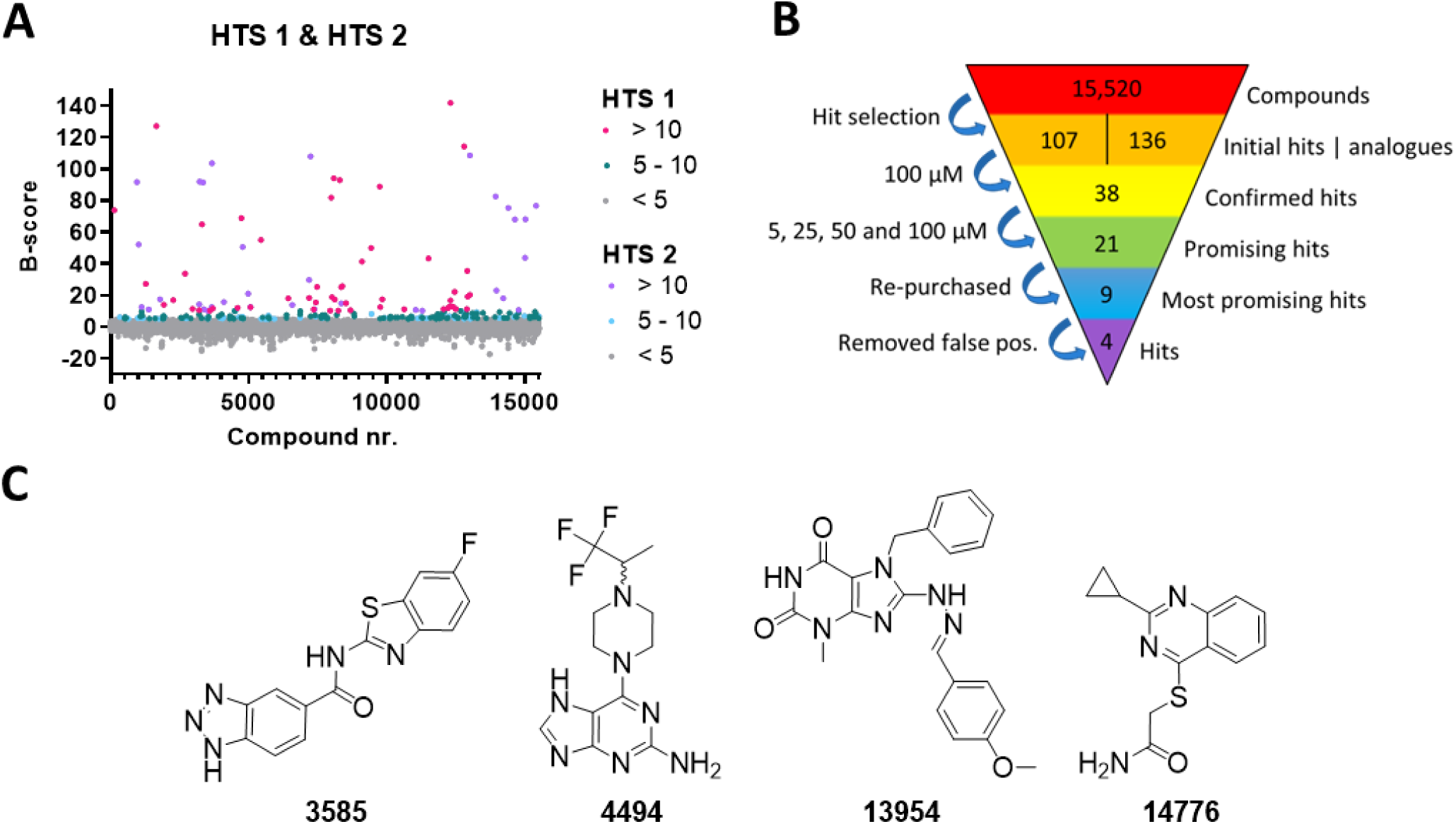
Results of the HTS competitive binding assay for the *Fnu* PreQ_1_ riboswitch. **A)** Calculated B-scores for all screened compounds in both high throughput screenings. High activity hits (B-score >10) are indicated in pink (HTS 1) or purple (HTS 2), moderate activity hits (B-score = 5–10) in green (HTS 1) or blue (HTS 2), inactive compounds (B-score <5) in grey. **B)** Workflow of hit selection and confirmation. **C)** Chemical structures of the four identified hits: **3585**, **4494**, **13954** and **14776**.

The four selected hits (Figure 5C) present typical drug-like properties^45^, with an average molecular weight of 323 Da, 1.8 H-bond donors, 4.8 H-bond acceptors, 3.5 rotatable bonds and a calculated octanol-water partition coefficient (cLogP) of 2.39. The total hit rate was 0.03%, which is in line with typical hit rates found for RNA targeting high throughput screenings^36,46^. For these hits, EC_50_ values were determined using both the WT and C17U mutant *Fnu* PreQ_1_ riboswitch (Figure 6A-B and Table 2). All hits exhibited competitive binding only at high ligand concentration (EC_50_ >100 µM) for both the WT and mutant PK. Unfortunately, higher ligand concentrations were limited by the solubility of the compounds in the aqueous buffer. Compound **4494**, the only compound soluble at millimolar concentrations, showed a decrease in competitive binding activity for the mutant *Fnu* PK, suggesting a similar binding mode as PreQ_1_.

**Figure 6.**
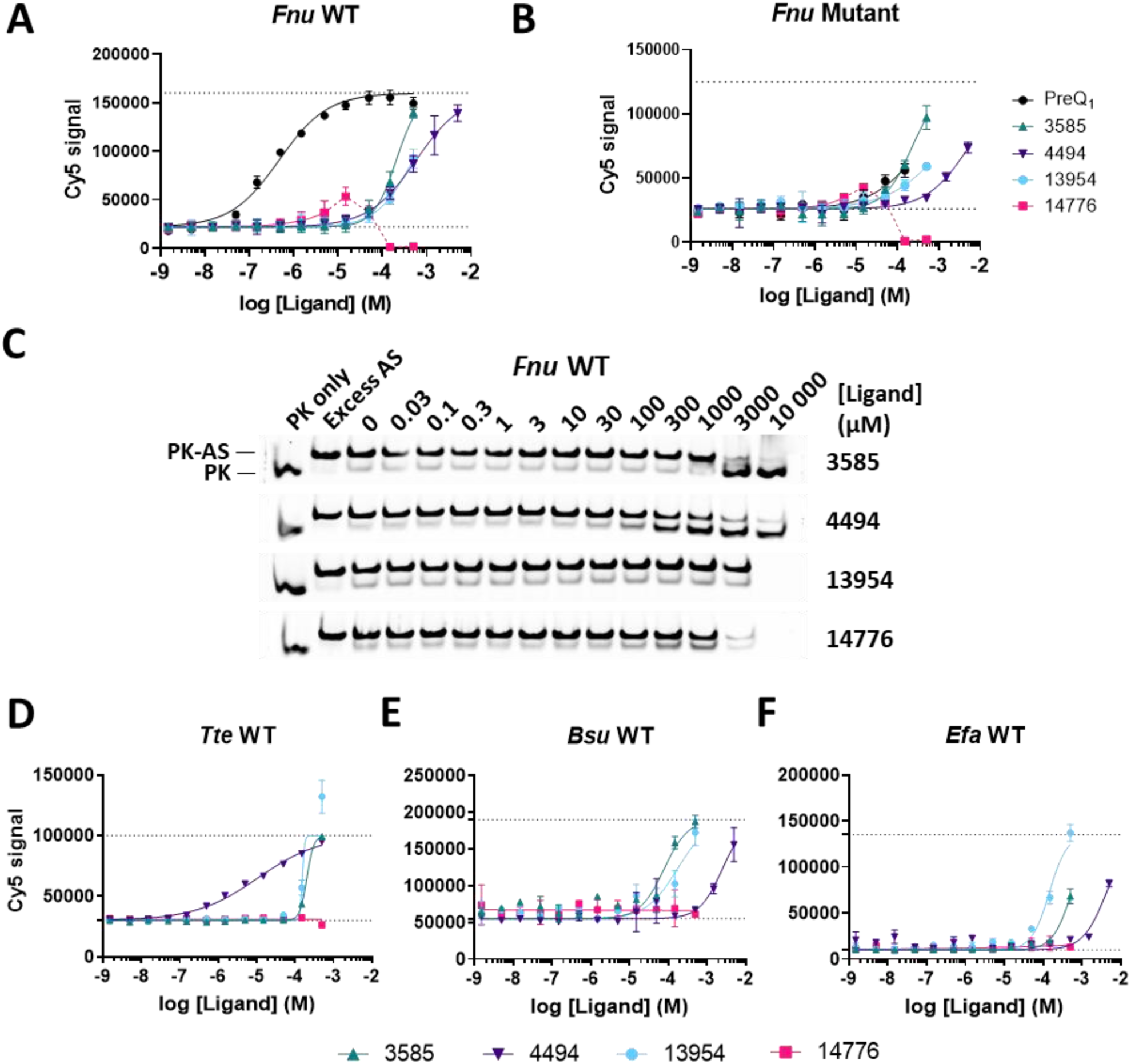
Hit validation of identified hits. **A-B)** Competitive binding assay of identified hits for the *Fnu* WT PK (**A**) and *Fnu* mutant PK (**B**). Dotted lines denote the minimum and maximum Cy5 signal as determined by the positive and negative controls. **C)** Competitive binding assay with the *Fnu* WT PK visualized on native PAGE. Cy5-PK is pre-incubated with ligand, after which unlabeled AS is added and the two conformations (PK-ligand and PK-AS) are separated on gel. **D-E)** Competitive binding assay of identified hits and the *Tte* WT PK (**D**), *Bsu* WT PK (**E**) and *Efa* WT PK (**F**).

**Table 2.**
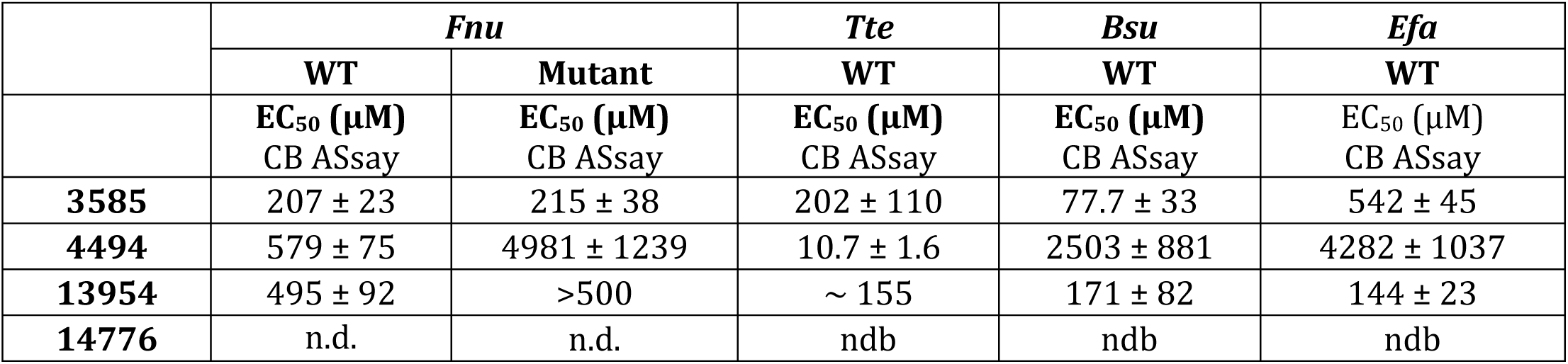
Overview of calculated EC_50_ values of the identified hits, as determined for the *Fusobacterium nucleatum* (*Fnu*), *Thermoanaerobacter tengcongensis* (*Tte*), *Bacillus subtilis* (*Bsu*) and *Enterococcus faecalis* (*Efa*) WT and *Fnu* C17U mutant PreQ_1_ riboswitches. n.d. = not determined due to compound interference. ndb = no detectable binding

To assess if ligand binding indeed induces PK formation, the CB ASsay was visualized with native PAGE (Figure 6C). Again, only **4494** demonstrated a clear dose-dependent competition with the AS (EC_50_ = 446 µM). While both **3585** and **14776** displayed a more prominent PK band at higher concentrations, it remains unclear whether they promote ligand-induced PK formation as **3585** shows an unlikely, rapid turning point for PK formation and **14776** is a highly colored compound that quenches fluorescent signal at high concentrations and thus interferes with the read-out. Further assessments with assays that do not rely on fluorescence may provide more insight into their binding behavior. Unfortunately, compound **13954** did not significantly promote PK formation on native gel.

To determine the selectivity of the hit compounds, the CB ASsay was also conducted in the presence of tRNA and with the *Tte*, *Bsu* and *Efa* PreQ_1_ riboswitches. Notably, the addition of tRNA did not decrease the competitive binding of the hits, indicating that they do not bind nonspecifically to structured RNA (Figure S10). Intriguingly, the quenching effect of **14776** at high concentrations was also reduced by the presence of tRNA. Addition of tRNA in the HTS could potentially reduce the number of false positives caused by nonspecific binding and possibly reduce the number of false negatives due to quenching. Therefore, the competitive binding of the hits to *Tte*, *Bsu* and *Efa* PreQ_1_ riboswitches was also measured in the presence of tRNA (Figure 6D-F). Observed ligand activity varies a lot between different riboswitch sequences. Compound **14776** only exhibited some activity for *Fnu*, whereas **3585** and **13954** displayed a high micromolar EC_50_ for all riboswitches with slight variations between sequences. Notably, compound **4494** showed significantly higher activity for the *Tte* riboswitch (EC_50_ = 10.7± 1.6 µM) compared to *Fnu*, *Bsu* and *Efa*. Because of this, competitive binding of the hits to the *Tte* PK was also evaluated on native gel (Figure S11), which confirmed that **4494** showed significant competitive binding.

To further evaluate the riboswitch ligands, PreQ_1_ and the four hits were tested in functional assays, including frameshift and translation prevention assays. Frameshift assays measure the increase in −1 ribosomal frameshifting (−1 FS) that is caused by, for example, ligand-induced formation and/or stabilization of a tertiary RNA structure. A stable tertiary RNA structure impedes the ribosome from moving along the mRNA, which allows the ribosome to slip into a different reading frame and continue translation to produce an alternative protein product. Frameshift assays have previously been used to measure pseudoknot formation of the PreQ_1_ riboswitch upon addition of PreQ_1_ and analogues^38^. To evaluate hit binding, FS assays were carried out with the *Fnu*, *Bsu*, *Tte* and *Escherichia coli* (*Eco*) riboswitches (Figure S12). Frameshifting was induced for all riboswitches upon addition of PreQ_1_ (of note, high concentrations of PreQ_1_ inhibited this process). Compound **4494** was the sole hit that clearly enhanced frameshifting with the *Tte* riboswitch and, to lesser degree, the *Eco* riboswitch, thus demonstrating its affinity to these riboswitches and its capacity to stabilize the pseudoknot structure. Compared to the *Bsu* and *Eco* riboswitches, the *Tte* riboswitch has a relatively accessible PreQ_1_ binding site (Figure S13), which could explain why the large (2,2,2-trifluoro-1-methylethyl)-piperazine group of **4494** is more easily accommodated in the *Tte* structure. These results underscore the importance of testing RNA target sequences from different strains to fully evaluate hit potential.

In the translation prevention assay, the translation efficiency of a small reporter peptide under the regulatory control of the PreQ_1_ riboswitch is assessed. Translational control by the riboswitch occurs through the sequestration of the Shine Dalgarno (SD) sequence within the second stem of the pseudoknot, thereby preventing ribosomes from binding the bacterial mRNA. For this assay, the *Tte* and *Eco* PreQ_1_ riboswitches were selected (Figure S14), given their roles as translational regulators in their biological context. Upon the addition of PreQ_1_, translation was markedly reduced by 90% for both the WT and C17U mutant *Tte* and *Eco* riboswitches, albeit the mutants required a higher concentration of PreQ_1_ to achieve this effect. Compounds **3585** and **14776** also exhibited translation reduction at millimolar concentrations, with minimal difference in effective concentrations for the WT and mutant riboswitches. Consequently, discerning whether the translation decrease is caused by specific ligand-RNA interactions or due to off-target effects becomes challenging. Remarkably, **4494** is the only compound able to reduce translation of the WT riboswitches while losing part of its activity against mutant riboswitches. This observation aligns with observations from the CB ASsay, indicating that **4494** reduces translation through specific riboswitch interactions. Furthermore, these results underscore the potential of **4494** to disrupt the biological function of PreQ_1_ riboswitches, rendering it a promising candidate for antibiotic drug development.

Thus, compound **4494** emerged as the most promising potential PreQ_1_ riboswitch ligand, displaying clear competitive binding on native gel, and losing activity in both CB and functional assays upon mutation of the cytosine crucial for binding with PreQ_1_. This suggested that **4494** may adopt comparable binding pose to the PreQ_1_ riboswitch. Upon examining the molecular structure of **4494**, a striking similarity to the chemical scaffold of PreQ_1_ becomes apparent, prompting further docking studies. Unfortunately, no high-resolution crystal structure of the *Fnu* riboswitch was available. Therefore, possible binding modes of the identified hits were docked using the *Bsu* (PDB ID: 3FU2) and *Tte* (PDB ID: 6VUI) PreQ_1_ riboswitches.

The co-crystallized PreQ_1_ (Figure 7A and 7C) was removed from the structures and replaced by the hit molecule, after which the suggested binding poses were manually inspected and selected (Figure 7B and 7D, Figure S15-S18). The most probable docking poses of **4494** suggest that it forms hydrogen bonds with the essential cytosine, albeit at a different angle than PreQ_1_. The PreQ_1_ binding pocket does not allow for modifications at the carbonyl position, forcing the ligand to turn to accommodate the (2,2,2-trifluoro-1-methylethyl)-piperazine group. This is especially evident for *Bsu*, which has a narrow binding pocket that forces the ligand to turn almost 90 degrees. Alternatively, *Tte* has a larger and more accessible binding pocket, which is why it can probably accommodate docking poses that more closely resemble the PreQ_1_ orientation. This could also explain the higher affinity of **4494** to the *Tte* riboswitch. Of note, docking indicated no preference for a specific stereoisomer.

**Figure 7.**
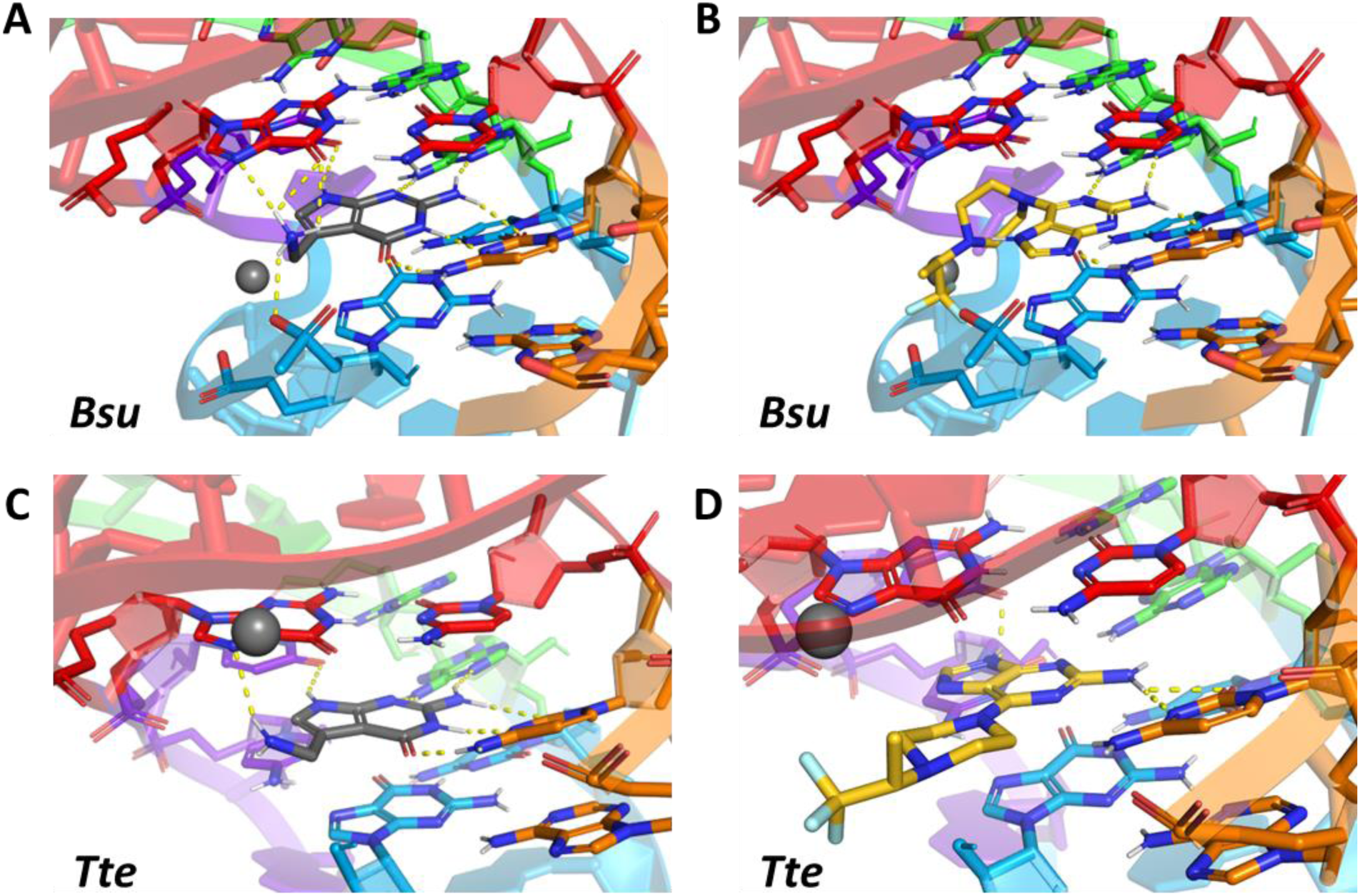
Docking of PreQ_1_ and **4494** to the *Bacillus subtilis* (*Bsu*) (PDB ID: 3FU2) and *Thermoanaerobacter tengcongensis* (*Tte*) PDB ID: 6VUI) PreQ_1_ riboswitch. Most probable binding pose of **4494** was manually selected. Hydrogen bonds are indicated with yellow dashed lines. **A)** Crystal structure of the *Bsu* PreQ_1_ riboswitch co-crystallized with PreQ_1_. **B)** Close up docked structure of **4494** bound to the *Bsu* PreQ_1_ riboswitch. **A)** Crystal structure of the *Tte* PreQ_1_ riboswitch co-crystallized with PreQ_1_. **D)** Close up docked structure of **4494** bound to the *Tte* PreQ_1_ riboswitch.

In both cases, the flipped confirmation results in a mismatched hydrogen bonding scheme, potentially explaining the observed lower affinity when compared to PreQ_1_. Interestingly, the proposed docking modes offer clear opportunities for hit optimization. It must be noted that the riboswitches were modelled as rigid structures, so possible rearrangement of RNA is not taken into account in this docking analysis. Nevertheless, the finding that PreQ_1_-like molecules can be modified at the WCF-face and still retain some affinity to the PreQ_1_ riboswitch is rather surprising. Given that most published modifications to PreQ_1_ have focused on the aminomethyl group^9,28,47,48^, exploring new modification at the carbonyl position may provide new insights for developing the next generation of PreQ_1_ riboswitch ligands.

## Discussion

In the emerging field of small-molecule RNA targeting, there is a growing demand for new HTS methods. Currently, RNA-targeted small-molecule screenings are conducted using *in silico*, *in vitro* or *in cellulo* methods. *In silico* approaches, while relatively fast and cost-effective, require high-resolution structural knowledge of the target RNA and are susceptible to false positives and negatives if they cannot accurately account for RNA flexibility^35^. Cellular assays identify biologically active hits, but false positives and negatives can arise from off-target interactions or poor cellular uptake. *In vitro* screening methods make use of purified RNA structures, offering the advantage of circumventing issues with off-target effects, cellular uptake, or lack of structural information. However, many current *in vitro* HTS methods rely on immobilizing the RNA target or small molecules^32,49,50^, which can influence the RNA conformation and/or the rotational freedom of the ligand, potentially altering ligand-RNA interactions. Other methods, including mass spectrometry (MS) or nuclear magnetic resonance (NMR), can detect low affinity compounds but are susceptible to artifacts or are low throughput, respectively^46^. One successful method that integrates experimental data with *in silico* screening is the 2D combinatorial screening (2DCS), which led to the creation of Inforna database^51^. This database is used to select RNA ligands for relatively simple RNA motifs, while ligands for neighboring motifs within the RNA target can be assembled into high-affinity hits. Another assay that is similar to our HTS method is the recently described strand-invasion assay that allowed the selection of high affinity ligands for the theophylline aptamer^36^. However, targeting more complex 3D RNA structures has not been reported thus far.

Here we present the development of an *in vitro* and in-solution high throughput competitive binding ASsay to identify new small-molecule ligands for more complex tertiary structured RNA targets, complemented by a custom screening workflow for HTS data analysis (see Supporting KNIME files). The potential of the assay was demonstrated by conducting a HTS with the clinically relevant *Fnu* PreQ_1_ riboswitch, leading to the discovery of four hits. These hits underwent further evaluation using both CB ASsays and functional frameshifting and translation prevention assays against the *Fnu*, *Bsu*, *Efa*, *Tte* and *Eco* PreQ_1_ riboswitches. The most promising hit, compound **4494**, exhibited competitive binding activity for the WT *Fnu*, *Tte*, *Bsu* and *Efa* riboswitches, clearly promoted *Fnu* and *Tte* pseudoknot formation when visualized on native PAGE and both stimulated frameshifting and prevented translation with the *Tte* riboswitch. In addition, activities in the competitive binding- and functional assays were diminished upon a C17U mutation of the essential cytosine in the riboswitch binding pocket, indicating a binding mode similar to that of the natural ligand PreQ_1_. This was supported by molecular docking, which suggested that **4494** can be recognized in a comparable fashion by the riboswitch, albeit at a different angle to accommodate the (2,2,2-trifluoro-1-methylethyl)-piperazine group. The docking studies also revealed clear opportunities for ligand optimization that could enhance the binding potency of **4494**. Importantly, this research shows that the CB ASsay is able to detect weak but biologically relevant hits for clinically relevant, tertiary RNA structures.

While this research presents the development of a CB ASsay for the PreQ_1_ riboswitch, it is likely applicable to other diverse RNA targets. Previous competitive binding assays rely on competition with specific (fluorescent) binders, such as the Tat peptide from the HIV-1-TAR hairpin^32–35^, which limit the number of targetable RNA structures. In contrast, this assay employs an AS that is in theory customizable for any RNA or DNA target. Moreover, the assay stringency is tunable by changing the length or number of modified nucleotides of the AS^36^, allowing for identification of hits with varying binding potencies. Another advantage of the CB ASsay is that all the components of the screening are in solution, allowing for the native folding of the RNA target and full rotational freedom of the ligand. While not performed in this work, selectivity tests could be incorporated into the HTS by addition of non-specific, unlabeled RNA (like tRNA or total RNA), or specific unlabeled RNA counter-targets to which the desired ligand should not bind. Furthermore, the CB ASsay can easily be downscaled, decreasing material usage, and enabling a higher throughput. Last but not least, the assay is quick, exemplified by the fact that the entire Enamine RNA focused library of 15,520 compounds was screened within half a day. All materials for the HTS can be obtained from established commercial sources at a relatively low cost, ensuring readily available high-quality materials and reducing variability between screenings.

Developing the CB ASsay for a new RNA target requires certain prior knowledge about RNA secondary structure. The AS should ideally target a secondary structural element containing both one strand of an RNA duplex and part of an interesting small-molecule binding pocket. Additionally, the target sequence should play a significant role in the global folding or the biological function of the RNA structure, increasing the probability of discovering biologically active ligands. The optimal AS length or nucleotide composition can first be determined with native PAGE, provided that the target and target-AS complex are not equally compact. Certainly, the availability of a positive control ligand greatly facilitates the process of fine-tuning the stringency of the assay. If no such control exists, it is recommended to test different AS lengths or modified nucleobase content, until one exhibits ideally an 80% bound – 20% unbound pattern with an equimolar ratio of target to AS. Based on our experience, this pattern yields a reasonable signal-to-background ratio in the CB ASsay, while allowing for the detection of ligands with moderate affinity.

It is important to note that the CB ASsay is an artificial biochemical assay, meaning that the assay conditions do not perfectly replicate a biological system. Differences in natural context of the target RNA, including flanking RNA sequences and cellular environment, may lead to alternative RNA (folding) conformations or ligand-binding behaviors. Secondary assays for hit validation to confirm ligand-target interactions in a biological environment, along with validation by biological assays are highly recommended. However, in the scenario where hits do not display biological activity but exhibit promising binding affinity and/or selectivity, they may still prove valuable by functionalizing them with RNA degrader molecules such as RIBOTACs^52,53^ or PINADs^54^.

In summary, the CB ASsay stands to be a valuable tool for future drug discovery platforms aimed at targeting complex RNA structures. Future efforts directed towards exploring other complex RNA structures, including those identified in SARS-CoV-2, will not only broaden our understanding but also play a pivotal role in advancing the development of innovative treatments for a wide range of diseases.

## Associated content

The custom KNIME workflows used for HTS data analysis are available free of charge on Zenodo at DOI: 10.5281/zenodo.10912410

## Author information

## Author Contributions

S.E.L.W. developed the competitive binding assay under supervision of M.A. and R.C.L.O. The HTS was performed by S.E.L.W. under guidance of P.P.G. and B.R.v.D. Secondary validation of hits was carried out by S.E.L.W., J.S.H, M.D.T. and I.B. Docking simulations were performed by A.P.A.J. S.E.L.W., M.A. and R.C.L.O conceived the idea, supervised the project and wrote the manuscript.

## Notes

The authors declare no competing financial interest.

## Acknowledgments

We thank Leiden University (KIEM grant to M.A., S.E.L.W. and R.C.L.O, ref: SAILOR) for financial support. Prof. dr. G.J.P. van Westen is kindly acknowledged for the access to the molecular docking tools needed for this work. Special acknowledgment is extended to Prof. dr. J.M.F.G. Aerts, head of the Medical Biochemistry Department at the LIC, for his valuable insights and constructive comments throughout the project.

## Supplementary Information

### Methods & Materials

#### Materials

All labeled oligonucleotides (HPLC purified) Sigma-Aldrich or IDT. All unlabeled oligonucleotides (desalted) were purchased from Sigma-Aldrich and used without further purification. All used materials were certified as DNAse and RNAse free.

##### Sequences

**Table S1:**
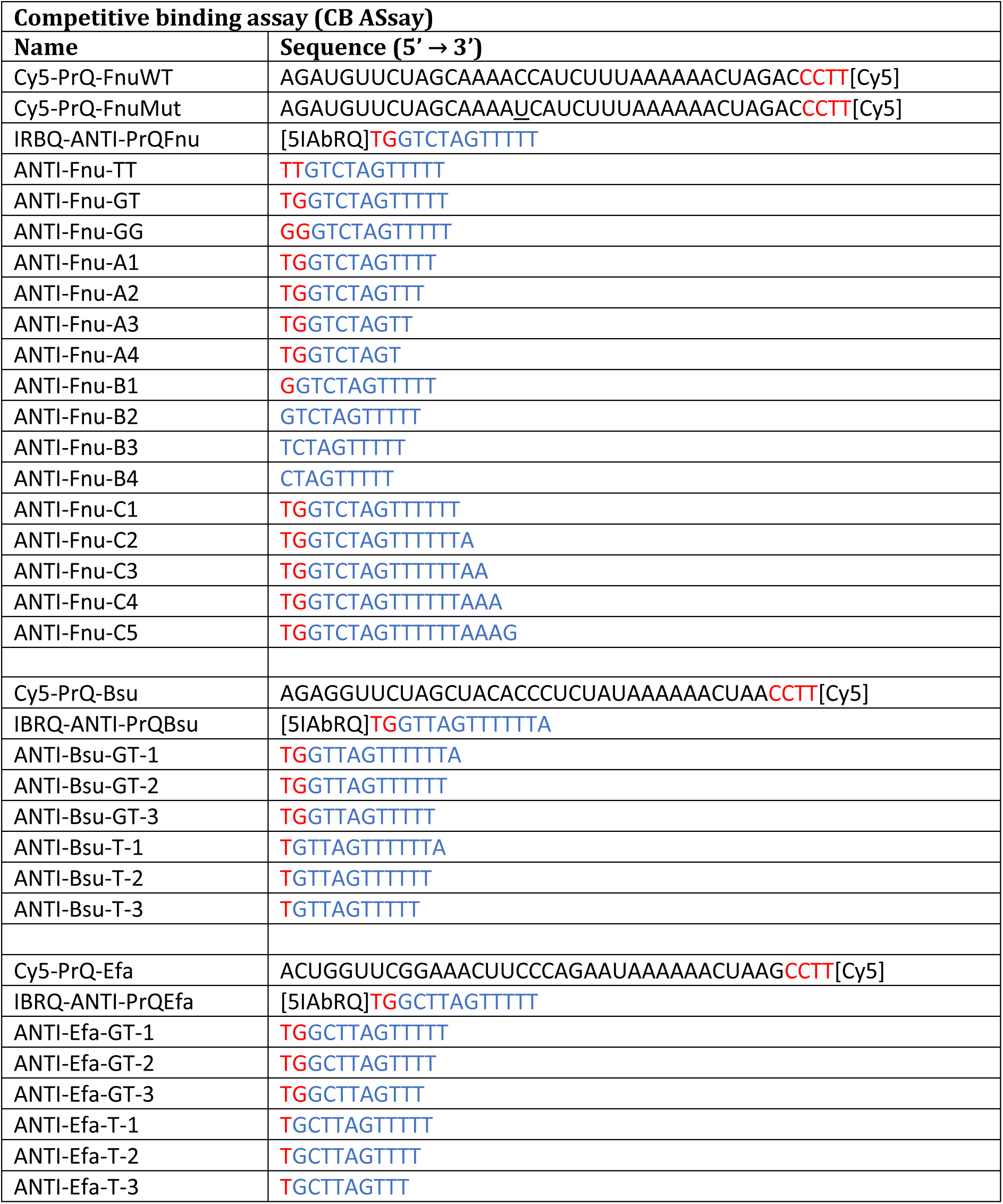
Sequences of the Competitive Binding Assay (CB ASsay). RNA indicated in black. Linker (DNA) indicated in red. DNA indicated in blue. Mutated bases are <u>underlined</u>.

**Table S2:**
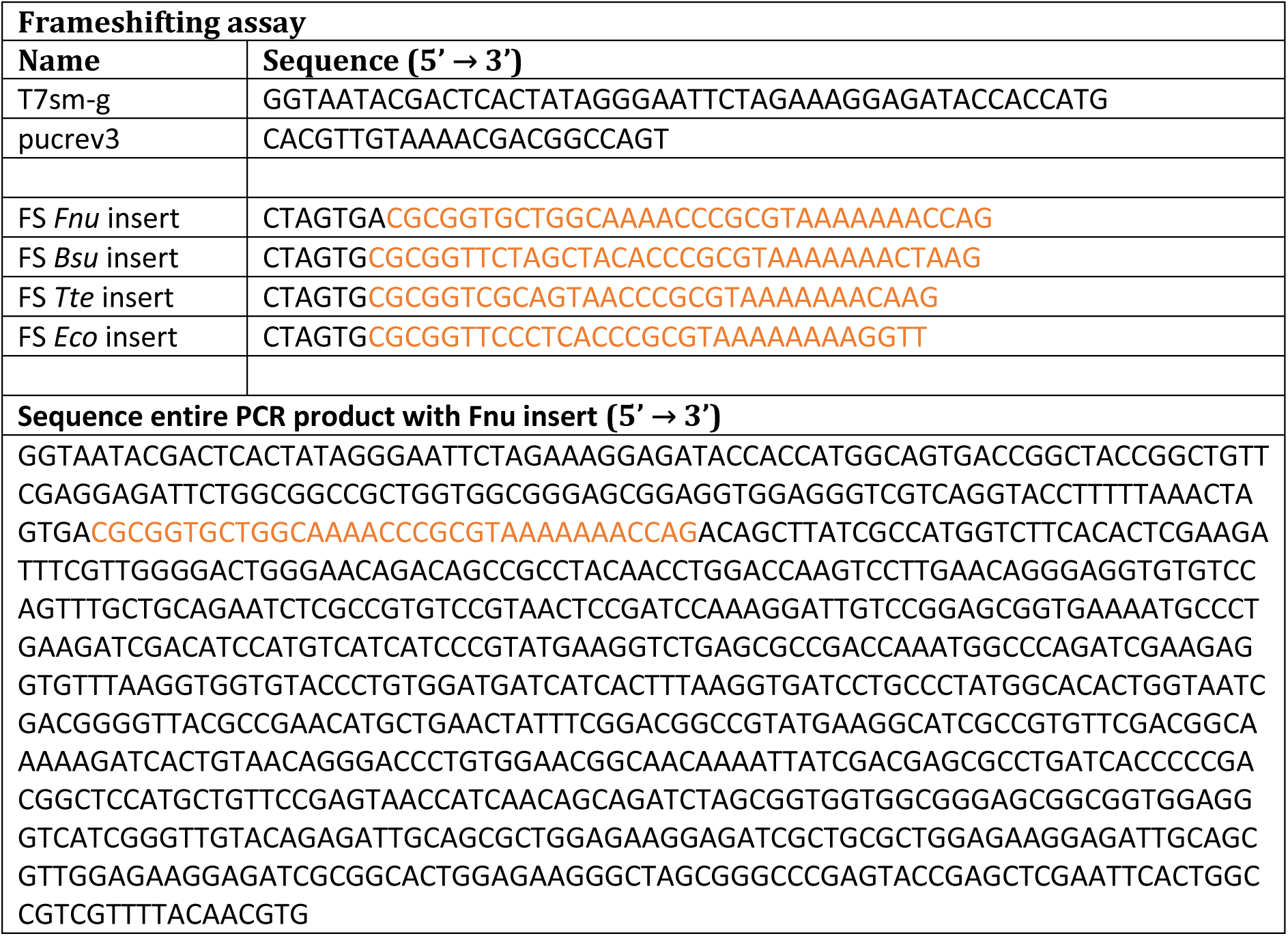
Sequences of the Frameshifting assay. PreQ_1_ riboswitch sequences indicated in orange.

**Table S3:**
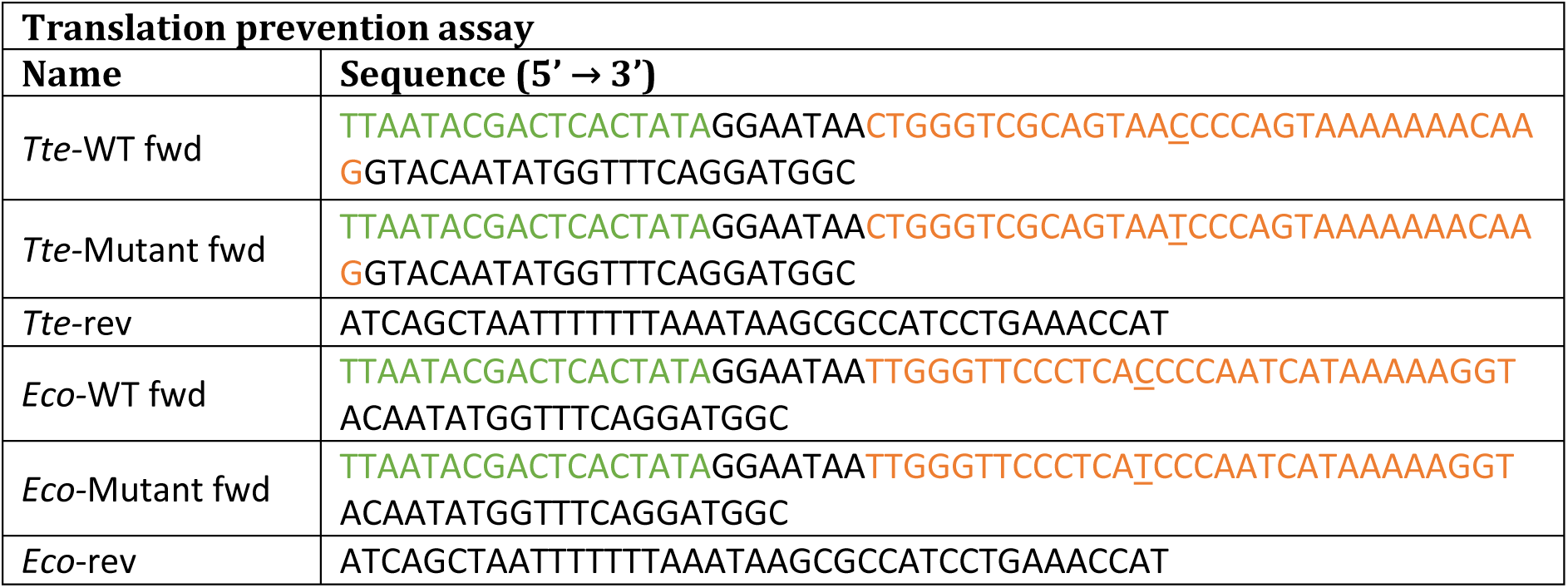
Sequences of the Translation prevention assay. The T7 polymerase promotor is indicated in green. PreQ_1_ riboswitch sequences indicated in orange. The essential cytosine (WT) or thymine (mutant) are underlined.

### Methods

#### Competitive Binding (CB) antisense assay (ASsay)

For all CB ASsays, Cy5-labeled PreQ_1_ riboswitch (Cy5-PK) and IowaBlack® RQ-labeled antisense (IBRQ-AS) were dissolved and diluted to 1 µM in milli-Q. All ligands were dissolved in DMSO. Positive control: a 10-fold excess of unlabeled AS. Negative control: DMSO. All CB experiments were carried out in a CB buffer (100 mM Tris (pH=7.6), 100 mM KCl, 10 mM NaCl, 1 mM MgCl_2_, 0.1% (v/v) DMSO and 0.01% (v/v) Tween20).

In a typical CB ASsay, 0.5 µL ligand was pipetted into a black 384-wells plate (Greiner Bio-One, Small Volume, 784076). To this, a mixture of 0.5 µL 1 µM Cy5-PK solution (in milliQ) and 6.5 µL of CB buffer was added, and the solution was incubated at RT for 1 hour. After this, a mixture of 0.5 µL 1 µM IBRQ-AS and 2.5 µL CB buffer was added, after which the mixture was mixed by pipetting up and down, and subsequently incubated at RT for additional 2 hours. Final ligand concentrations ranged from 0.0015 to 500 µM. The fluorescence was measured with a CLARIOstar (BMG LABTECH) using λ_ex_ = 610 ± 30 nm, λ_em_ = 675 ± 50 nm, gain = 2200 and focal point = 10.2. Obtained fluorescent signals were plotted against ligand concentration. EC_50_ values were determined in GraphPad Prism 8 using the ‘[Inhibitor] *versus* response – Variable slope (four parameters)’ analysis with Bottom constraint = average of DMSO control and Top constraint = average of 10-fold excess unlabeled AS.

#### High throughput competitive binding (HTS CB) ASsay

The commercially available RNA-focused library, containing 15,520 compounds (10 mM in DMSO) divided over 49 384-wells Low Dead Volume (LDV) ECHO plates (Labcyte), was purchased from Enamine and this library was screened in duplo.

In a typical HTS CB ASsay, 25 nL of ligand (10 mM) was added to a black 384-wells plate (Greiner Bio-One, Small Volume, 784076) using an ECHO550 Acoustic Liquid Handler (Labcyte). To this, a mixture of 0.5 µL of Cy5-PK (1 µM) and 6.5 µL CB buffer was added with a BioTek Multiflo FX dispenser (BioTek). Then, the plate was shaken for 10 seconds at medium speed and incubated at RT for 1 hour. After this, a mixture of 0.5 µL of IBRQ-AS (1 µM) and 2.5 µL CB buffer was added with a Multiflo dispenser, after which the plate was shaken for 10 seconds at medium speed, centrifuged for 1 minute at 1000 rpm and subsequently incubated at RT for 2.5 hours. The fluorecence was measured with a CLARIOstar (BMG LABTECH) using λ_ex_ = 610 ± 30 nm, λ_em_ = 675 ± 50 nm, gain = 2200 and focal point = 10.2.

Raw fluorescence data were analyzed in KNIME Analytics Platform with a custom workflow (see Supporting KNIME files). For all plates, Z’-scores were calculated to confirm high quality screening data. Data were normalized with standard KNIME nodes to yield Z-scores and B-scores. Hits were determined based on calculated B-scores, differentiating between two hit categories: ‘High activity hit’ = B-score >10 in a single screening, and ‘Moderate activity hit’ = B-score 5 - 10 in both screenings. Additionally, compounds with a B-score 3 - 10 in both screenings were manually inspected to prevent premature exclusion of potential hits. Initial hits were then manually grouped based on molecular structure, and close analogues with a SUM B-score of >6 (from both screenings) were included in subsequent hit confirmation.

#### Competitive binding assay on native gel electrophoresis

Cy5-labeled PreQ_1_ riboswitch (Cy5-PK) and IowaBlack® RQ -labeled antisense (IBRQ-AS) were dissolved and diluted to 1 µM in milli-Q, whereas all ligands were dissolved in DMSO. In all experiments, a 10x CB buffer (1 M Tris (pH=7.6), 1 M KCl, 100 mM NaCl, 10 mM MgCl_2_, 1% DMSO and 0.1% Tween20) was used. To an Eppendorf tube, 0.5 µL of Cy5-PK (10 µM), 3 µL of milli-Q water, 0.5 µL of 10x CB buffer and 0.5 µL of ligand (concentrations ranging from 5 nM to 100 mM) were added, and the mixture was incubated at RT for 1 hour. To this, 0.5 µL of IBRQ-AS (10 µM) was added and the mixture was subsequently incubated at RT for 10 minutes. Different RNA conformations were separated on a 20% native polyacrylamide gel at 4 °C, bands were visualized on a GelDoc (Bio-Rad) and processed using Image Lab software (Bio-Rad).

#### Microscale thermophoresis (MST)

Cy5-labeled *Fnu* WT PreQ_1_ riboswitch (Cy5-PK) was diluted to 80 nM in CB buffer (100 mM Tris (pH=7.6), 100 mM KCl, 10 mM NaCl, 1 mM MgCl_2_, 0.1% DMSO and 0.01% Tween20). A two-fold dilution series of ligand in buffer + DMSO was prepared, maintaining a final DMSO concentration of 5%. To the ligand, an equal volume of diluted Cy5-PK was added, resulting in a final Cy5-PK concentration of 40 nM and ligand concentrations ranging from 15.3 nM to 500 µM. Samples were incubated at RT for 15 min, after which the samples were transferred to standard coated capilaries (NanoTemper Technologies) and subsequently subjected to MST analysis. MST experiments were conducted in triplicate on a Monolith NT.115 system (NanoTemper Technologies). The results were analyzed with MO.Affinity Analysis v2.1.5 software, and the obtained ‘Fraction bound’ values were plotted against ligand concentrations. The apparent dissociation constants were determined in GraphPad Prism 8 using the [Inhibitor] vs response – variable slope (four parameters) analysis with Bottom constraint = 0 and Top constraint = 1.

#### Computational modelling of hits

Molecular docking was performed using the ICM Molsoft Pro suite. Briefly, the structure of the *Bsu* PreQ_1_ riboswitch in complex with PreQ_1_ (7-deaza-7-aminomethyl-guanine) (PDB ID: 3FU2) or *Tte* riboswitch (PDB: 6VUI) was retrieved from the Protein Data Bank using ICM Molsoft’s inbuilt feature. Of this structure chain C (3FU2) or chain A (6VUI) was converted to an ICM object, with ‘optimize hydrogens’ set to true. ‘Tight’ water molecules were retained, however this led to no conserved water molecules being present. The bound ligand was isolated to a separate object. The 3D pocket finding tool was utilized to identify the binding site. Of the potential binding sites identified, the pocket containing the co-crystallized ligand was selected. A docking project was set up based on this selection using default settings. The identified hits and their relevant tauto-meric forms were loaded from an SD-file and docked with the ‘thoroughness’ parameter set to 10, and the 10 best poses were retained. The resulting poses were manually inspected, selected, and visualized using the Open Source PyMOL application.

#### Frameshift assays

PreQ_1_ riboswitches were cloned into SpeI-HindIII digested plasmid pMOFS as described in Diweg, Oskam *et al.* (2023)^1^. This plasmid produces functional nanoluciferase when -1 frameshifing takes place. To enhance frameshifting, stem S1 of the PreQ_1_ pseudoknots was replaced by five GC base pairs as reported by Yu *et al.* (2013)^2^. Transcripts were synthesized by T7 RNA polymerase (NEB T7 high yield RNA synthesis kit) using PCR templates obtained by amplification with primers T7sm-g and pucrev3. Synthesized RNA was diluted to an approximate concentration of 10 ng/µL and 0.5 µL of this RNA solution was incubated with 0.5 µL ligand (dissolved in 8% DMSO in milli-Q) for 15 minutes at RT. Subsequently, 1 µL rabbit reticulocyte lysate (Nuclease-treated, Promega) was added and incubated at 28 °C for 45 minutes (final concentration DMSO was 2%). The reaction was quenched with 10 mM Tris-HCl (pH=7.5) and transferred to a white 96-wells plate, to which 2.5 µL of 100x diluted Nano-Glo® HiBiT Lytic Substrate (Promega) was added. Luminescence was measured on a GloMax Multi+ Detection System (Promega) and data were normalized to an in-frame control.

#### Translation prevention riboswitch assay

DNA constructs containing a T7 promotor, a PreQ_1_ riboswitch and the sequence of HiBiT Nano Luciferase were obtained from full-length primers (Sigma-Aldrich) that were annealed and amplified by PCR. Transcripts were synthesized by T7 RNA polymerase (NEB T7 high yield RNA synthesis kit) and were not purified before use. 1 µL of ligand (dissolved in 5% DMSO) was added to a 0.5 µL solution of 40x diluted RNA. Next, 0.5 µL of PURExpress® Solution A (New England Biolabs), 0.375 µL PURExpress® Solution B and 2.625 µL dilution buffer (10 mM Tris (pH=7.5), 202 mM KOAc and 15 mM Mg(OAc)_2_) were added, and the mixture was incubated for 30 minutes at 37 °C (final concentration DMSO was 2%). The reaction was quenched with 10 mM Tris-HCl (pH=7.5) buffer supplemented with LgBiT Protein (Promega). The solution was then transferred into a white 96-wells plate and 2.5 µL of Nano-Glo® substrate (Promega) was added. Luminescence was measured on a GloMax Multi+ Detection System (Promega) and normalized to the DMSO control.

### Supporting Figures and tables

**Figure S1:**
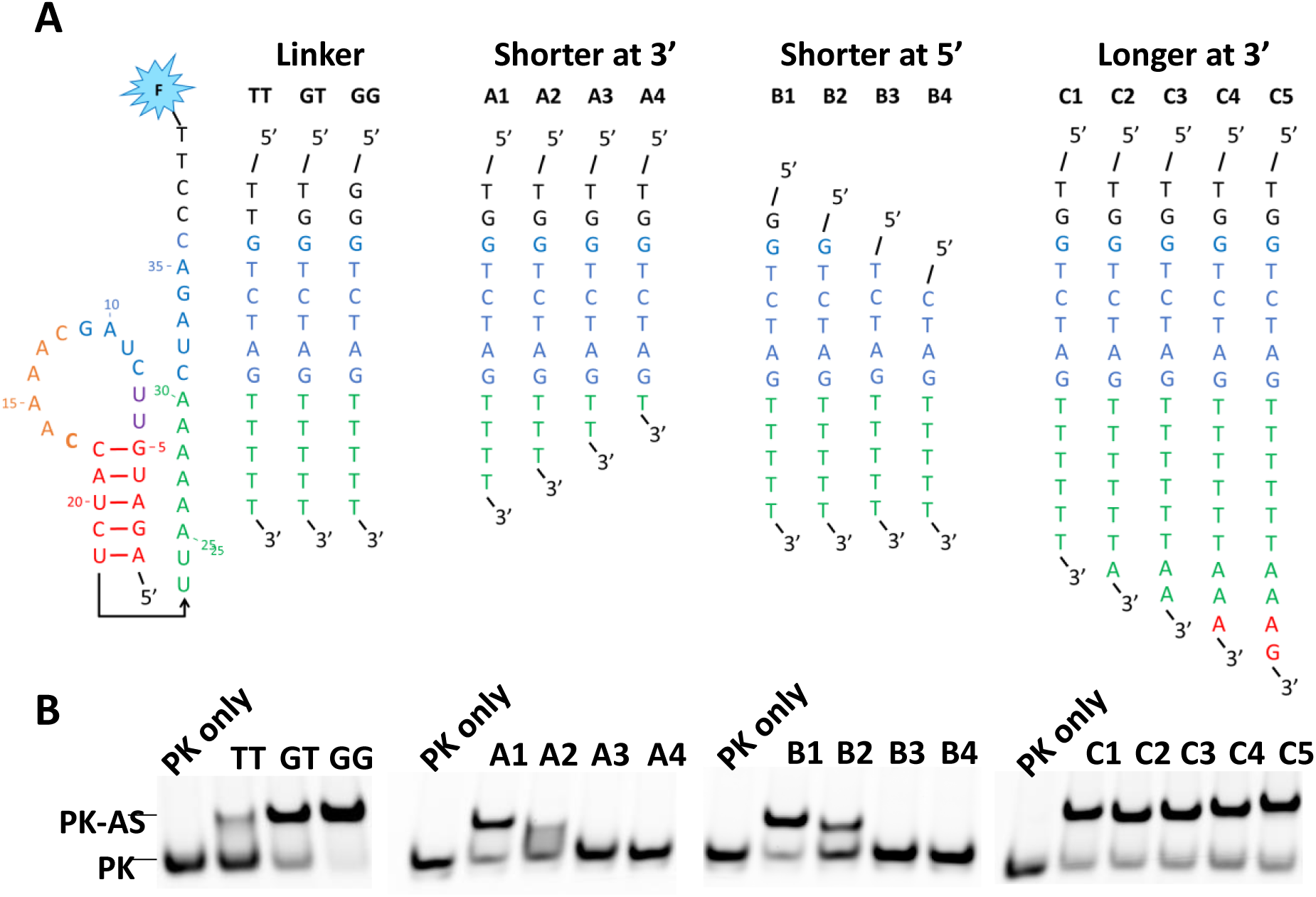
**A)** Antisense sequences screened for the optimization of the competitive binding ASsay for the *Fusobacterium nucleatum* (*Fnu*) riboswitch. **B)** Competitive binding ASsay of the *Fnu* riboswitch with PreQ_1_ and different antisenses, visualized on a native PAGE. Cy5-PK (1 eq.) was incubated with unlabeled AS (1 eq.) and the two possible conformations (PK and PK-AS) were separated on a 20% polyacrylamide gel.

**Figure S2:**
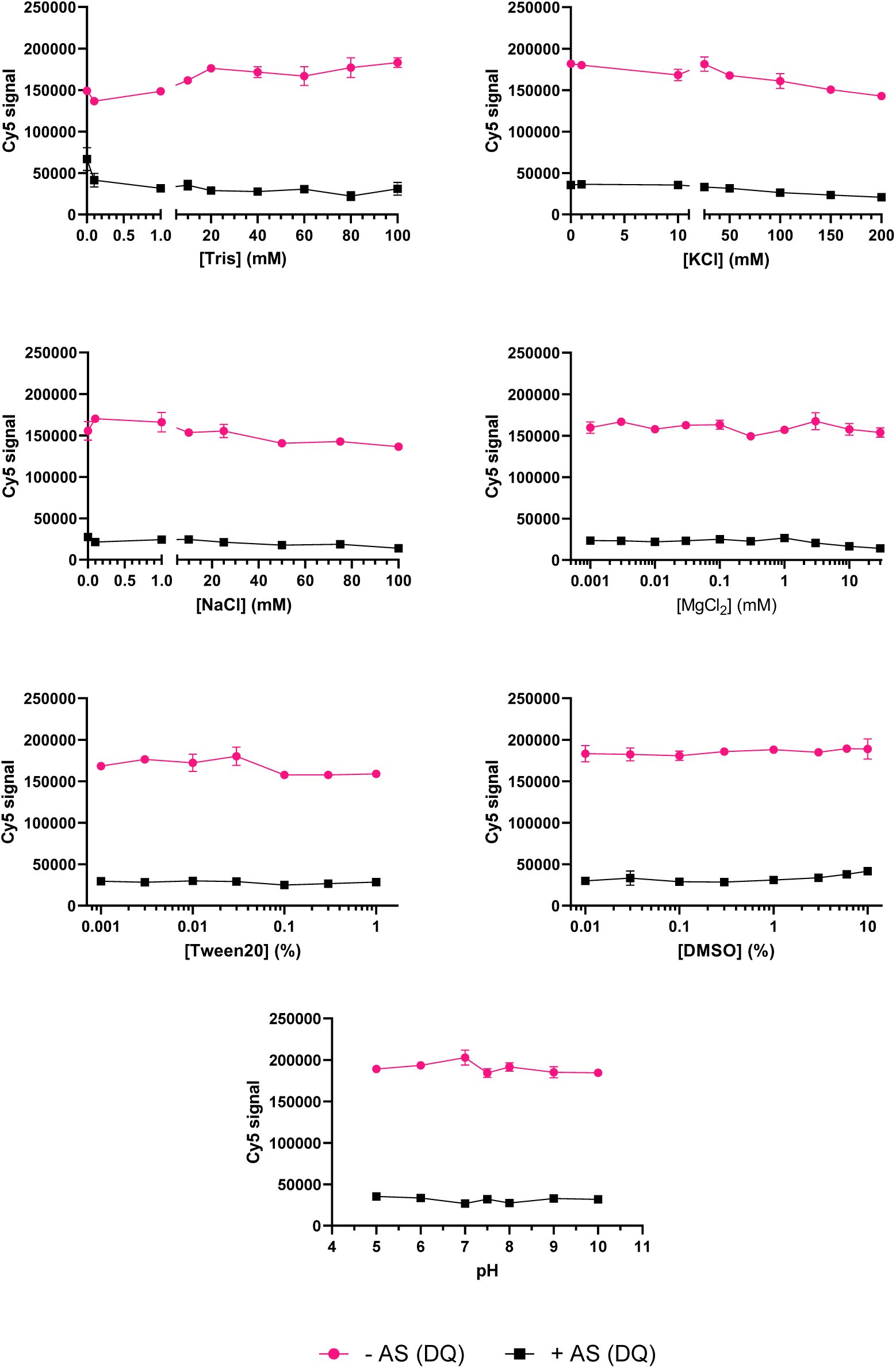
Buffer optimization for the competitive binding ASsay for the *Fnu* WT PreQ_1_ riboswitch. Standard buffer conditions were 100 mM Tris (pH = 7.5), 100 mM KCl, 10 mM NaCl, 1 mM MgCl_2_, 0.1% DMSO and 0.01% Tween20, unless varied for optimization.

**Figure S3:**
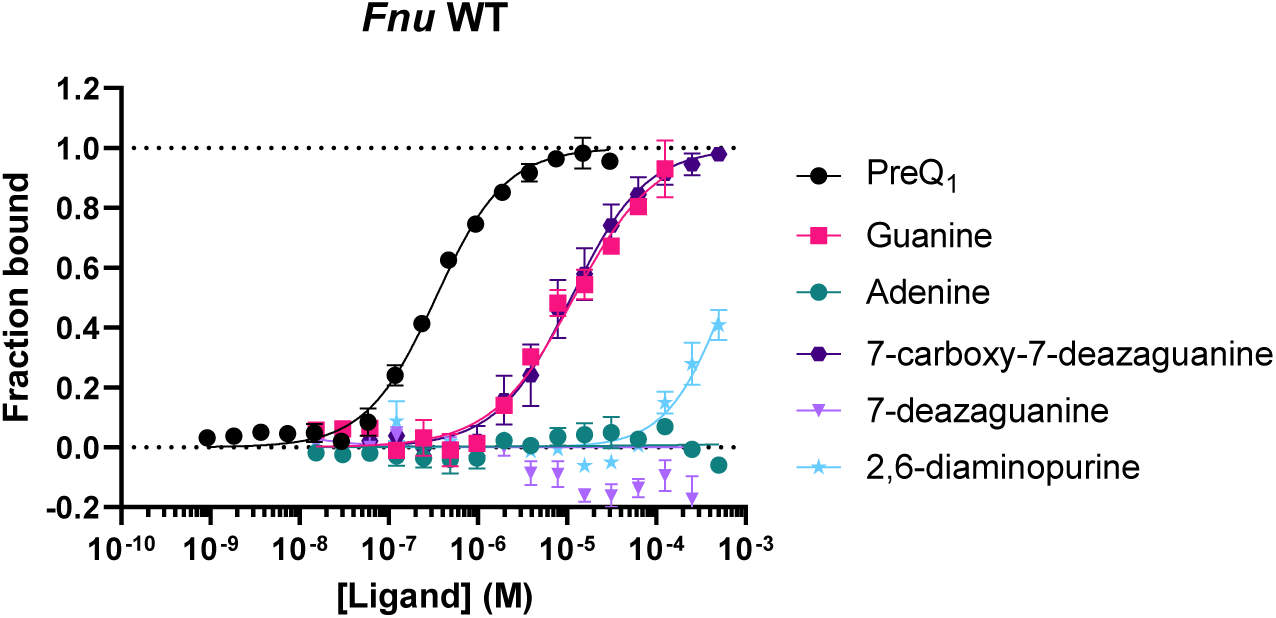
Microscale thermophoresis (MST) measurements of PreQ_1_, guanine, adenine, 7-carboxy-7-deazaguanine, 7-deazaguanine and 2,6-diaminopurine to the WT *Fnu* riboswitch. Displayed datapoints represent the average of three separate measurements.

**Figure S4:**
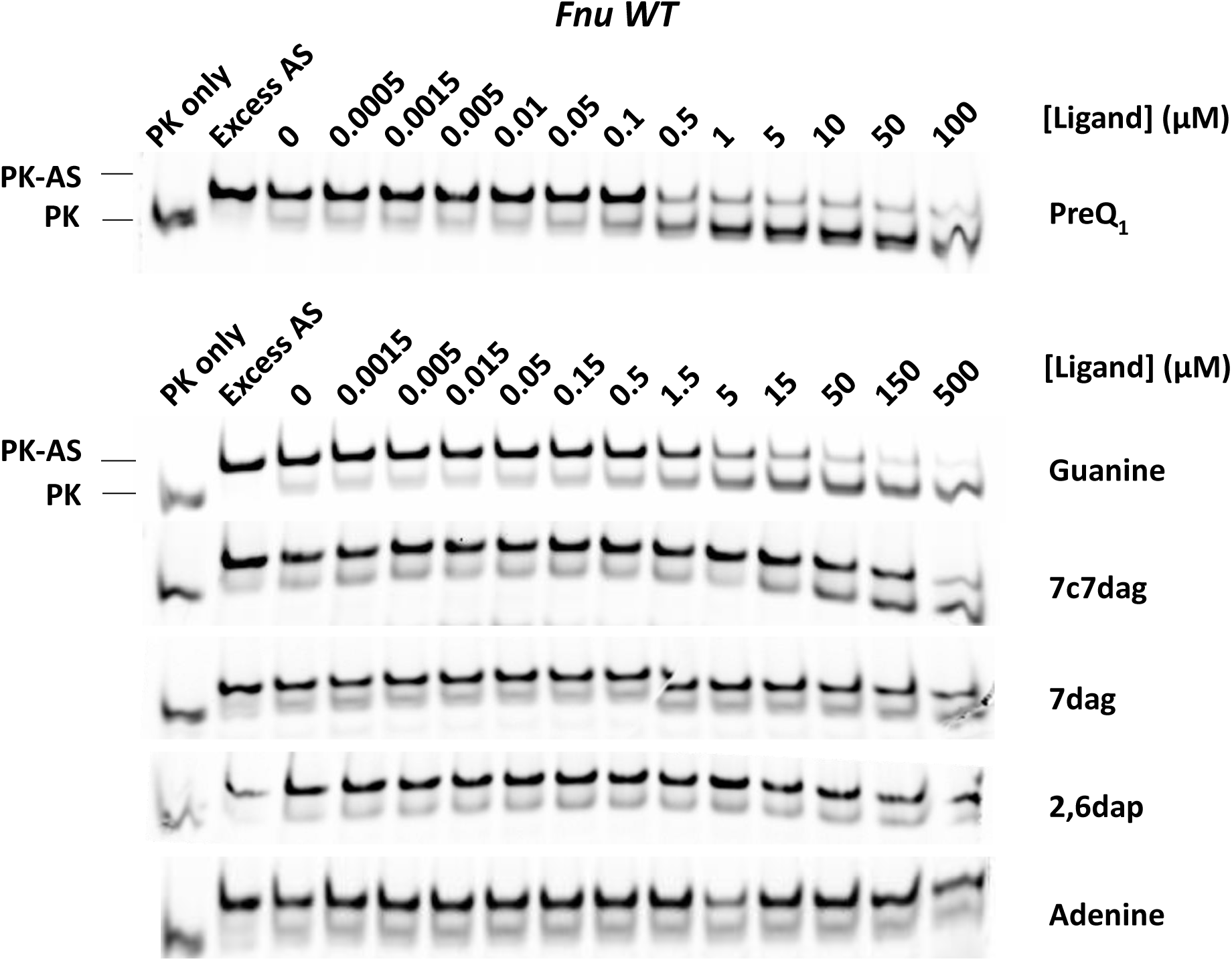
Competitive binding assay with the *Fnu* riboswitch visualized on native PAGE. Cy5-PK (1 eq.) was pre-incubated with a ligand, after which unlabeled AS (1 eq.) was added and the two possible conformations (PK and PK-AS) were separated on a 20% polyacrylamide gel.

**Figure S5:**
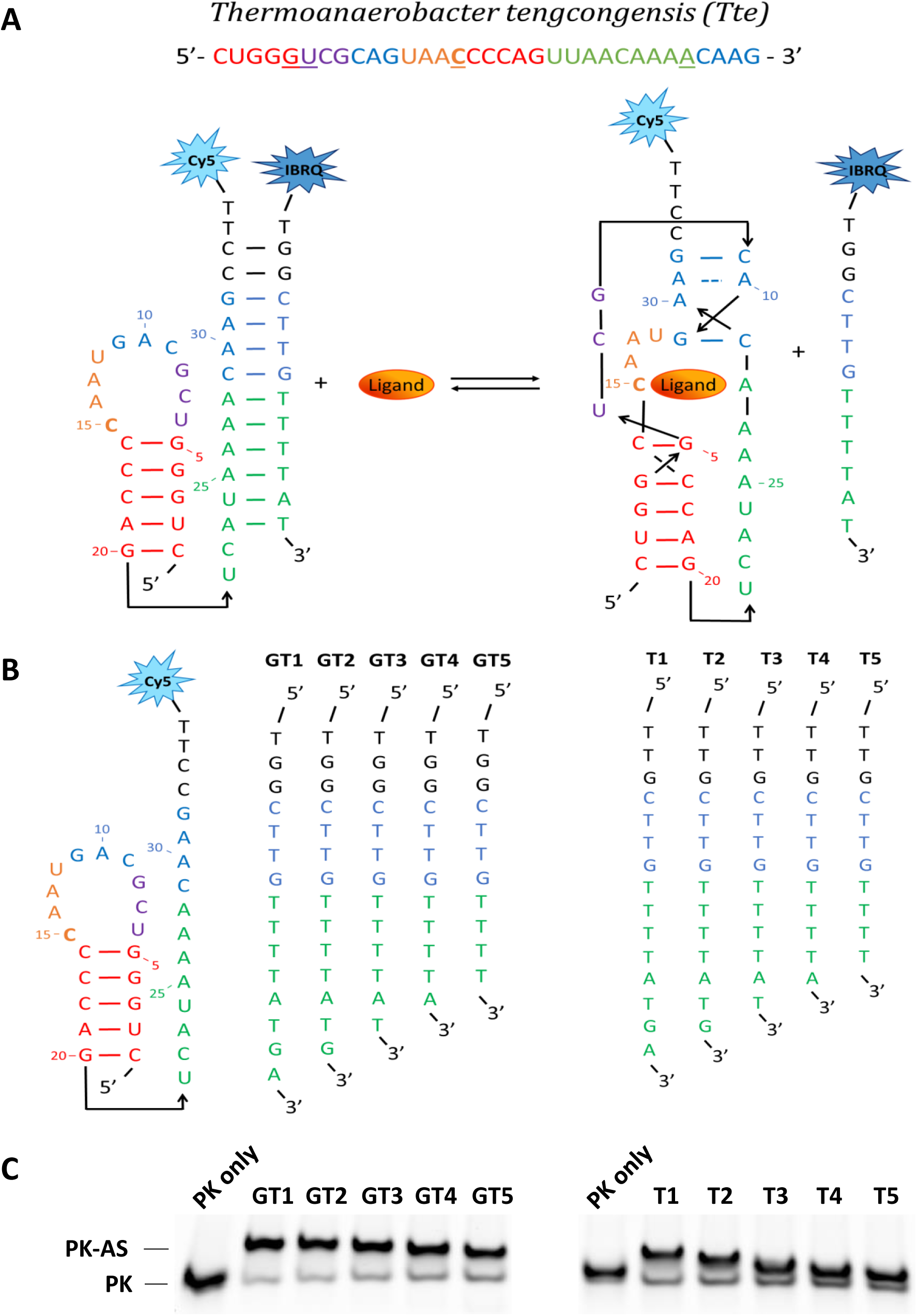
**A**) Schematic representation of the competitive binding ASsay for *Thermoanaerobacter tengcongensis* (*Tte*) PreQ_1_ riboswitch. **B)** Antisense sequences for optimizing the competitive binding assay for the *Tte* riboswitch. **C)** Competitive binding ASsay of the *Tte* riboswitch with PreQ_1_ and different antisenses, visualized on a native PAGE. Cy5-PK (1 eq.) was incubated with unlabeled AS (1 eq.) and the two possible conformations (PK and PK-AS) were separated on a 20% polyacrylamide gel.

**Figure S6:**
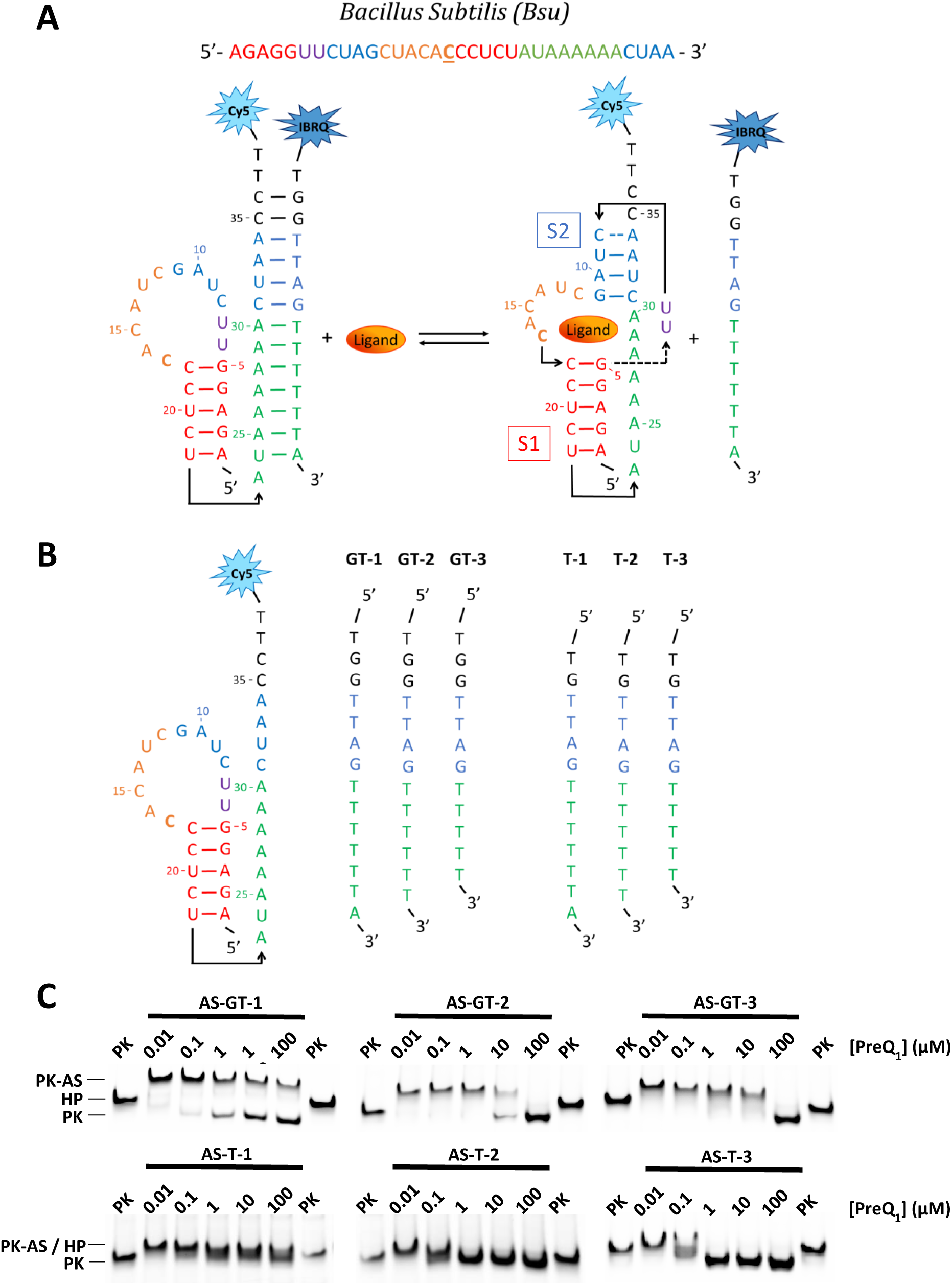
**A**) Schematic representation of the competitive binding ASsay for the *Bacillus subtilis* (*Bsu*) PreQ_1_ riboswitch. **B)** Antisense sequences for optimizing the competitive binding assay for the *Bsu* riboswitch. **C)** Competitive binding assay of the *Bsu* riboswitch with PreQ_1_ and different antisenses, visualized on native PAGE. Cy5-PK (1 eq.) was preincubated with PreQ_1_, after which unlabeled AS (1 eq.) was added and the two possible conformations (PK and PK-AS) were separated on a 20% polyacrylamide gel.

**Figure S7:**
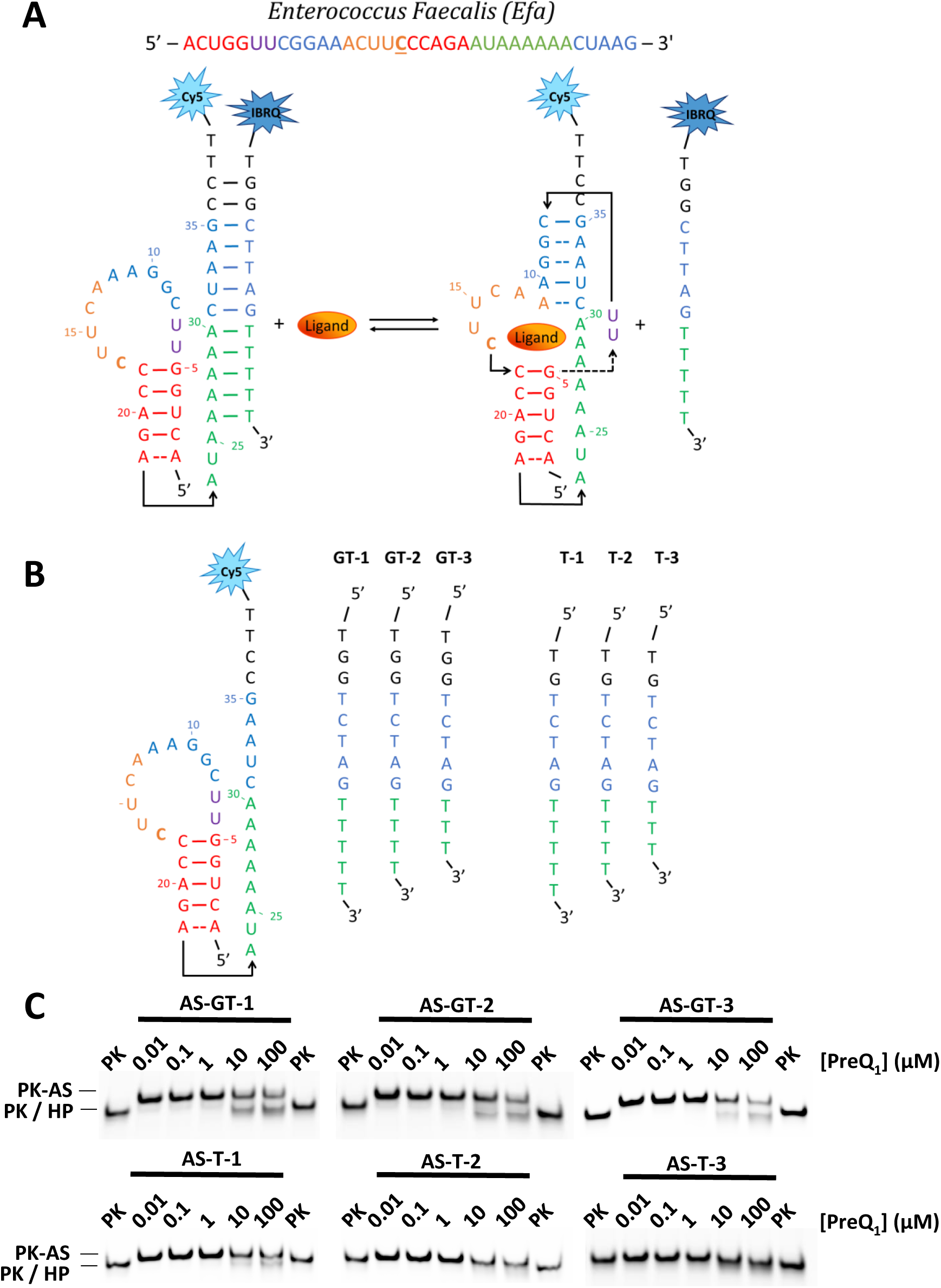
**A**) Schematic representation of the competitive binding ASsay for the *Enterococcus faecalis* (*Efa*) PreQ_1_ riboswitch (ligand-bound conformation not experimentally proven). **B)** Antisense sequences for optimizing the competitive binding assay for the *Efa* riboswitch. **C)** Competitive binding assay of the *Efa* riboswitch with PreQ_1_ and different antisenses, visualized on native PAGE. Cy5-PK (1 eq.) was pre-incubated with PreQ_1_, after which unlabeled AS (1 eq.) was added and the two possible conformations (PK and PK-AS) were separated on a 20% polyacrylamide gel.

**Figure S8:**
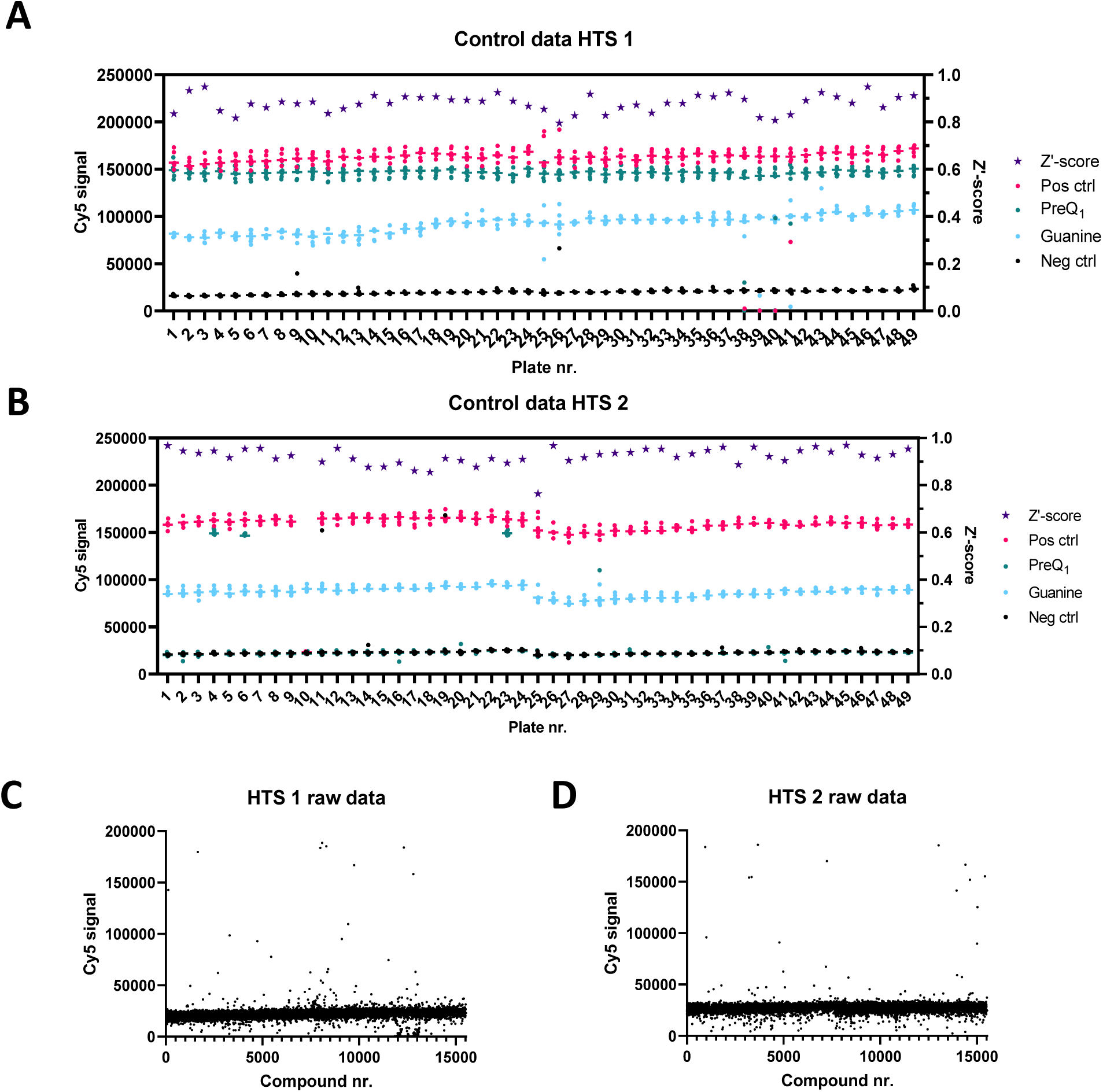
High throughput screening controls and raw data. **A-B**) Controls for HTS 1 (**A**) and HTS 2 (**B**), including on the left Y-axis the Cy5-signal of the positive control (10 eq. unlabeled antisense), negative control (DMSO), PreQ_1_ (25 µM) and guanine (25 µM), and on the right Y-axis the calculated Z’-scores. **C-D**) Raw fluorescence data of HTS 1 (**C**) and HTS 2 (**D**).

**Figure S9:**
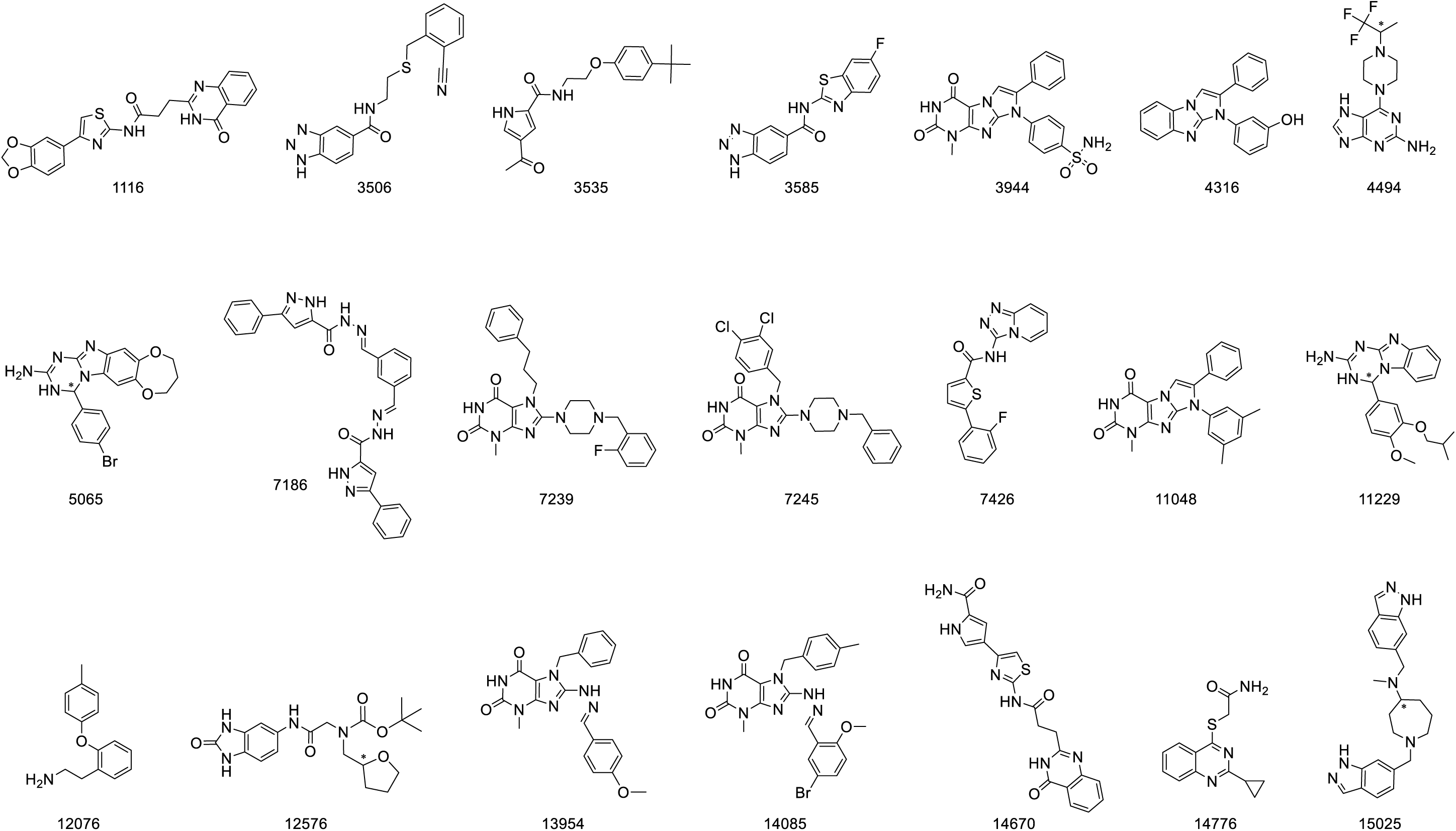
Molecular structures of most promising hits. * denotes a chiral center.

**Figure S10:**
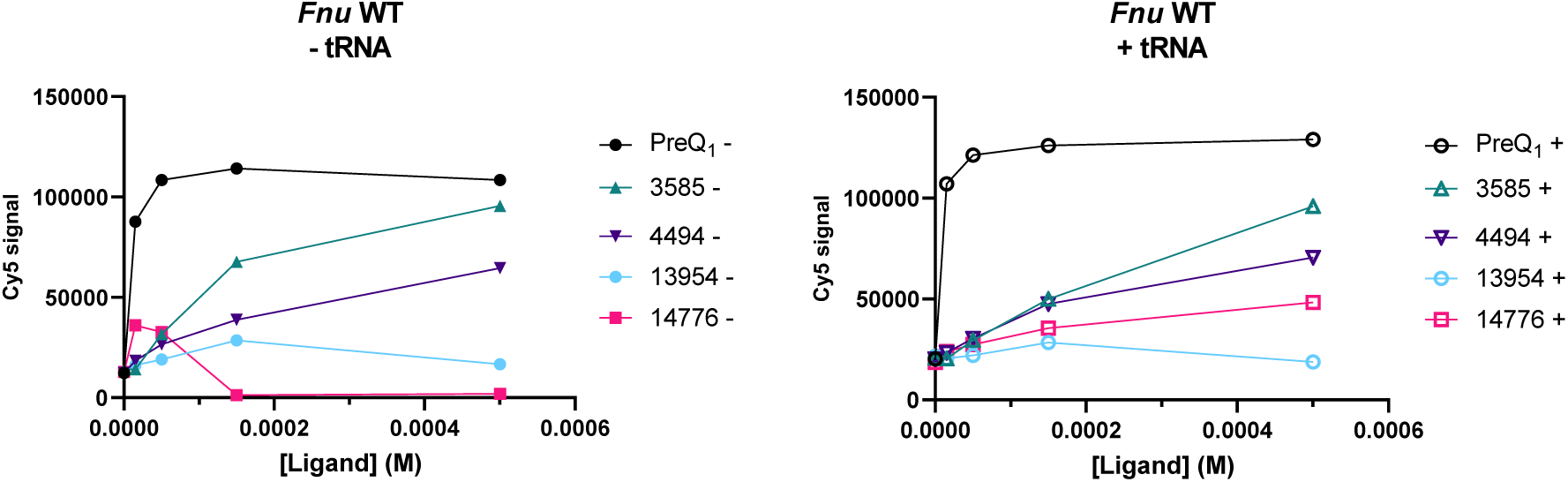
Competitive binding ASsay of PreQ_1_ and identified hits for the *Fnu* WT riboswitch, with (right) and without (left) 200 molar equivalents of tRNA.

**Figure S11:**
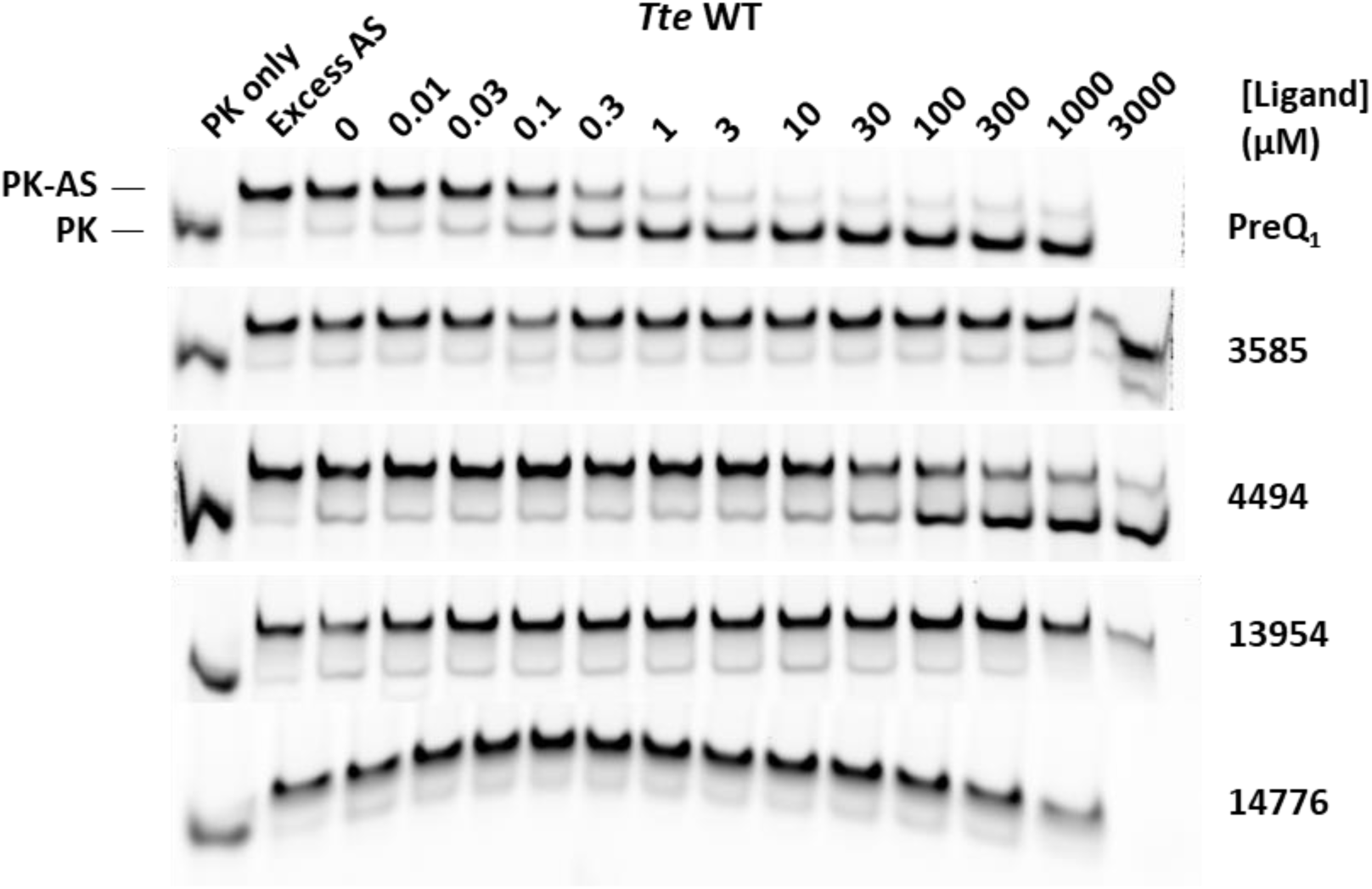
Competitive binding assay with the *Tte* riboswitch visualized on native PAGE. Cy5-PK (1 eq.) was preincubated with a ligand, after which unlabeled AS (1 eq.) was added and the two possible conformations (PK and PK-AS) were separated on a 20% polyacrylamide gel.

**Figure S12:**
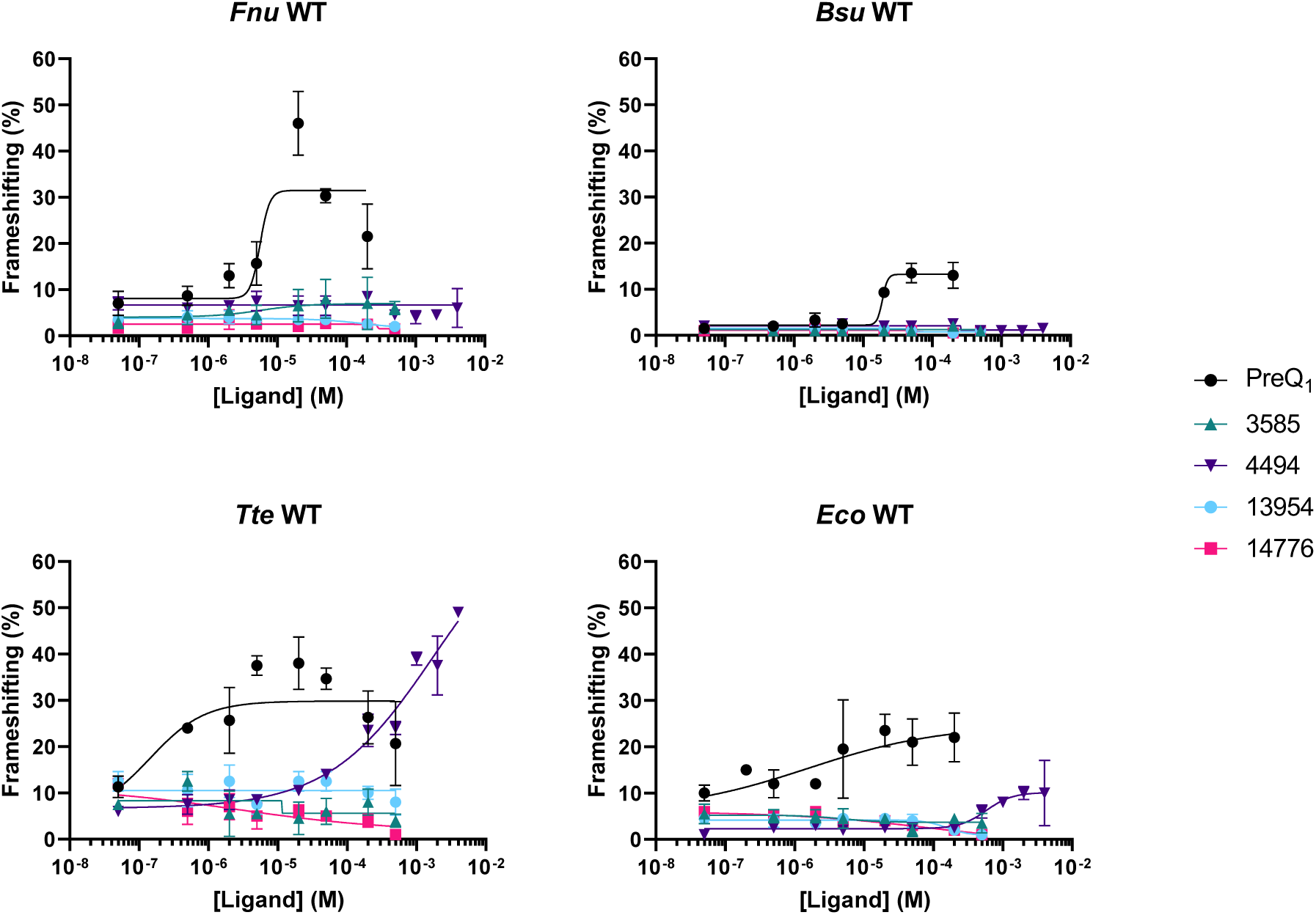
Frameshift assays of PreQ_1_ and hit compounds with the *Fnu*, *Bsu*, *Tte* and *Eco* WT PreQ_1_ riboswitches. Frameshift percentages were calculated by normalizing the luminescent signal to the in-frame control.

**Figure S13:**
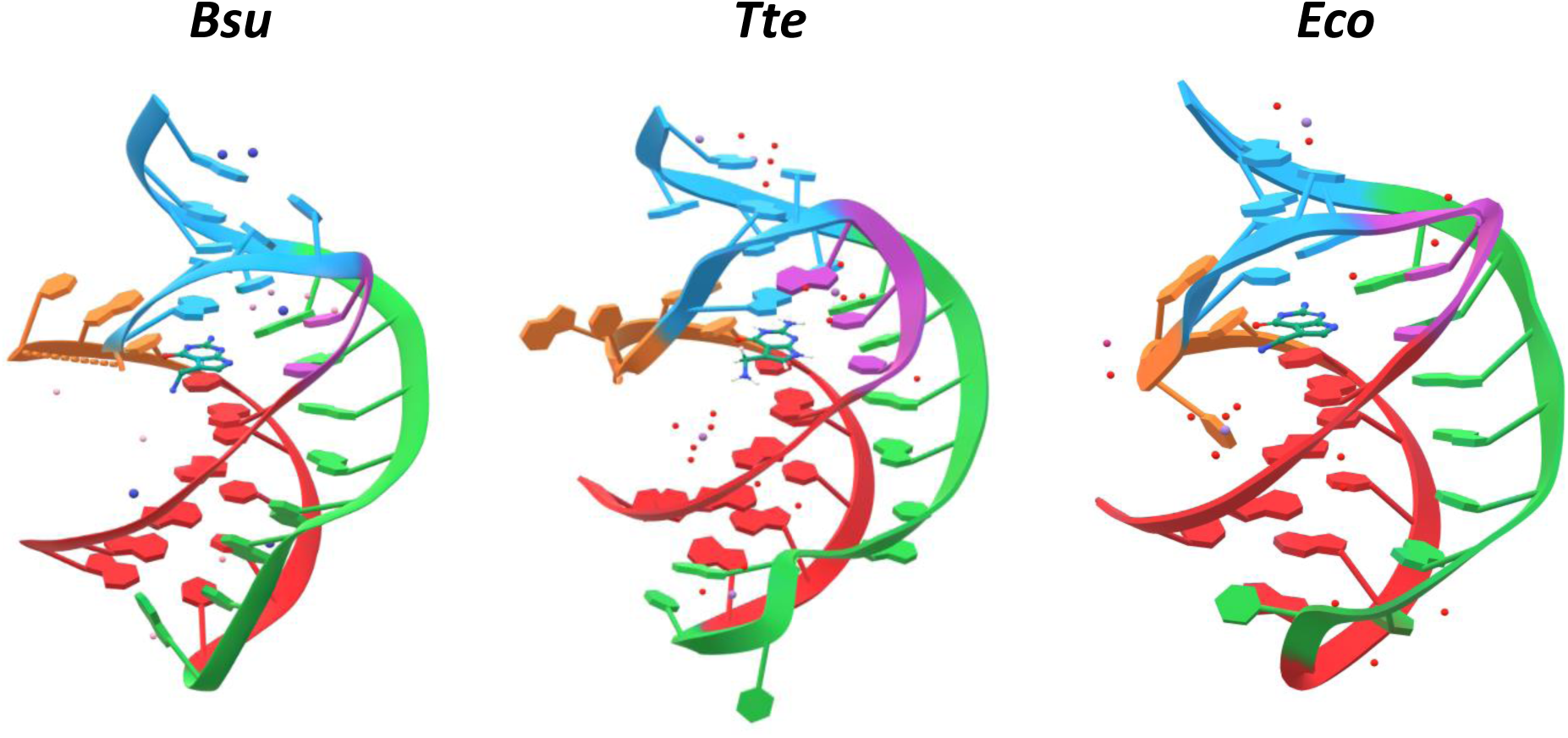
Published crystal structures of the *Bacillus subtilis* (*Bsu*) (PBD: 3FU2), *Thermoanaerobacter tengcongensis* (*Tte*) (PDB: 6VUI) and *Escherichia coli* (*Eco*) (PDB: 8FZA) PreQ_1_ riboswitches, co-crystallized with PreQ_1_.

**Figure S14:**
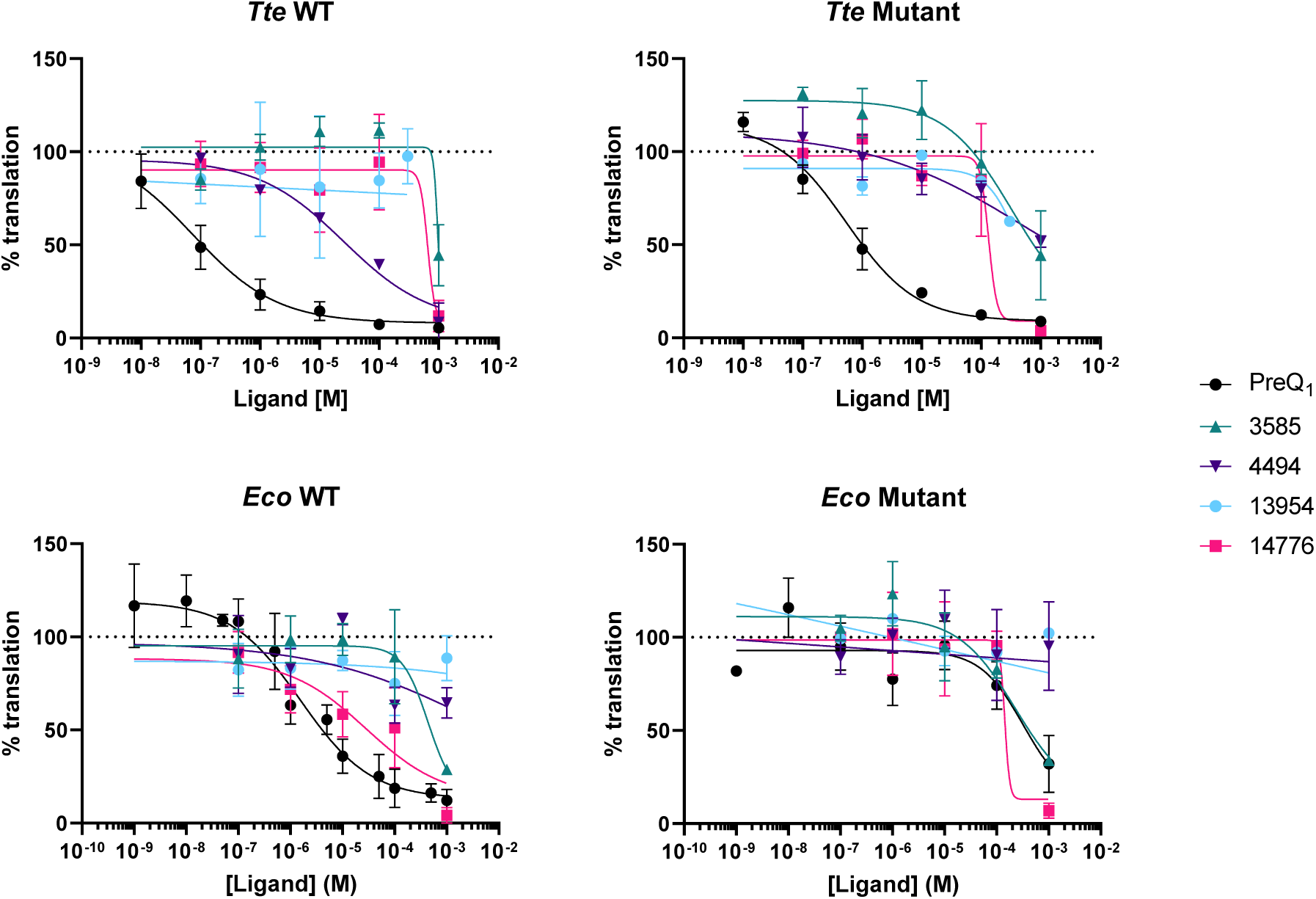
Translation prevention assays of PreQ_1_ and hit compounds with the *Tte* and *Eco* WT and mutant PreQ_1_ riboswitches. Translation percentages were calculated by normalizing the luminescent signal to the DMSO control.

**Figure S15:**
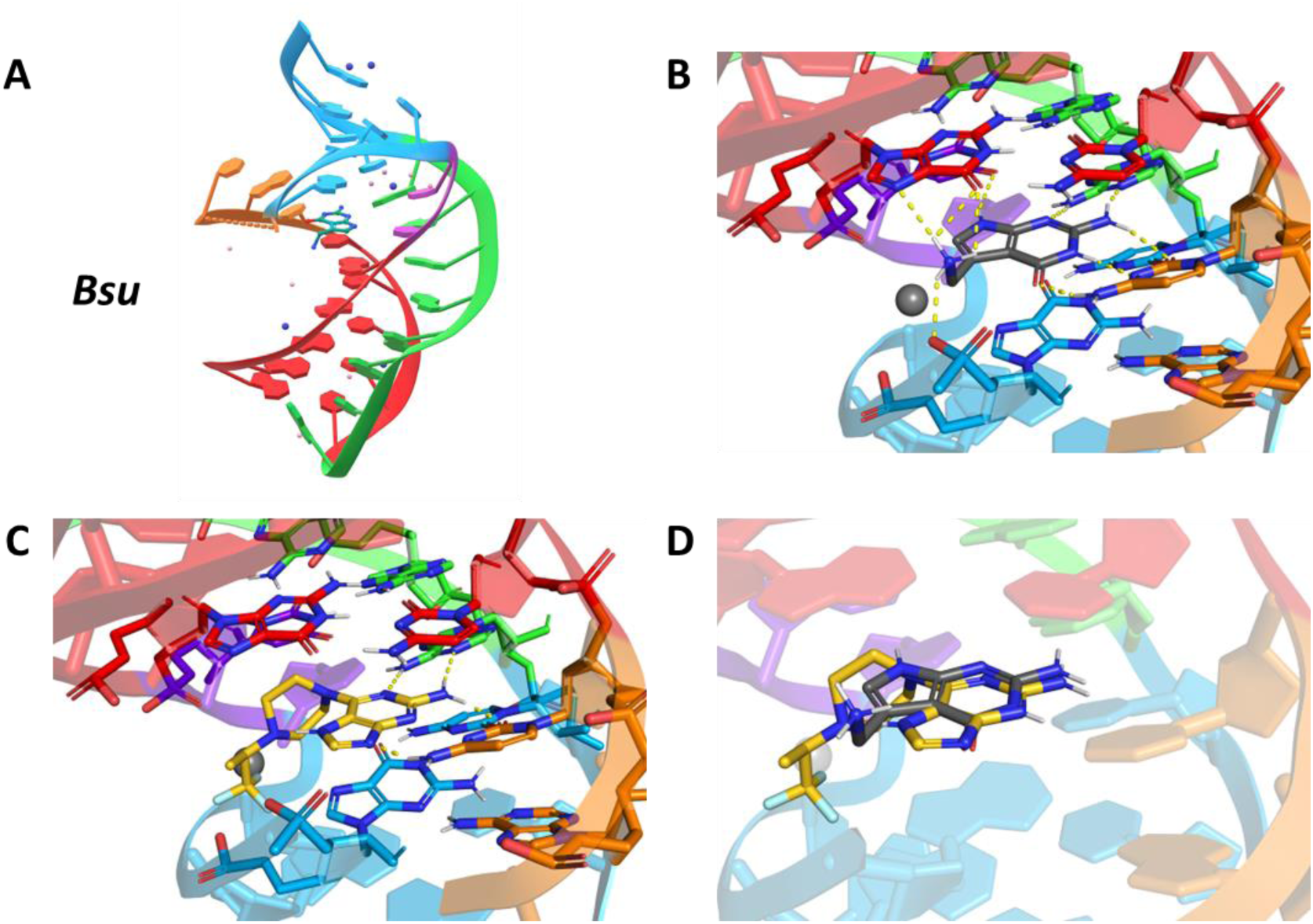
Docking of PreQ_1_ and **4494** to the *Bacillus subtilis* (*Bsu*) PreQ_1_ riboswitch (PDB ID: 3FU2). Most probable binding pose of **4494** was manually selected. Hydrogen bonds are indicated with yellow dashed lines. **A**) Crystal structure of the *Bsu* PreQ_1_ riboswitch co-crystallized with PreQ_1_. **B-C**) Close up docked structure of PreQ_1_ (**B**) or **4494** (**C**) bound to the *Bsu* PreQ_1_ riboswitch. **D**) Superimposed binding poses of PreQ_1_ and **4494** to the riboswitch.

**Figure S16:**
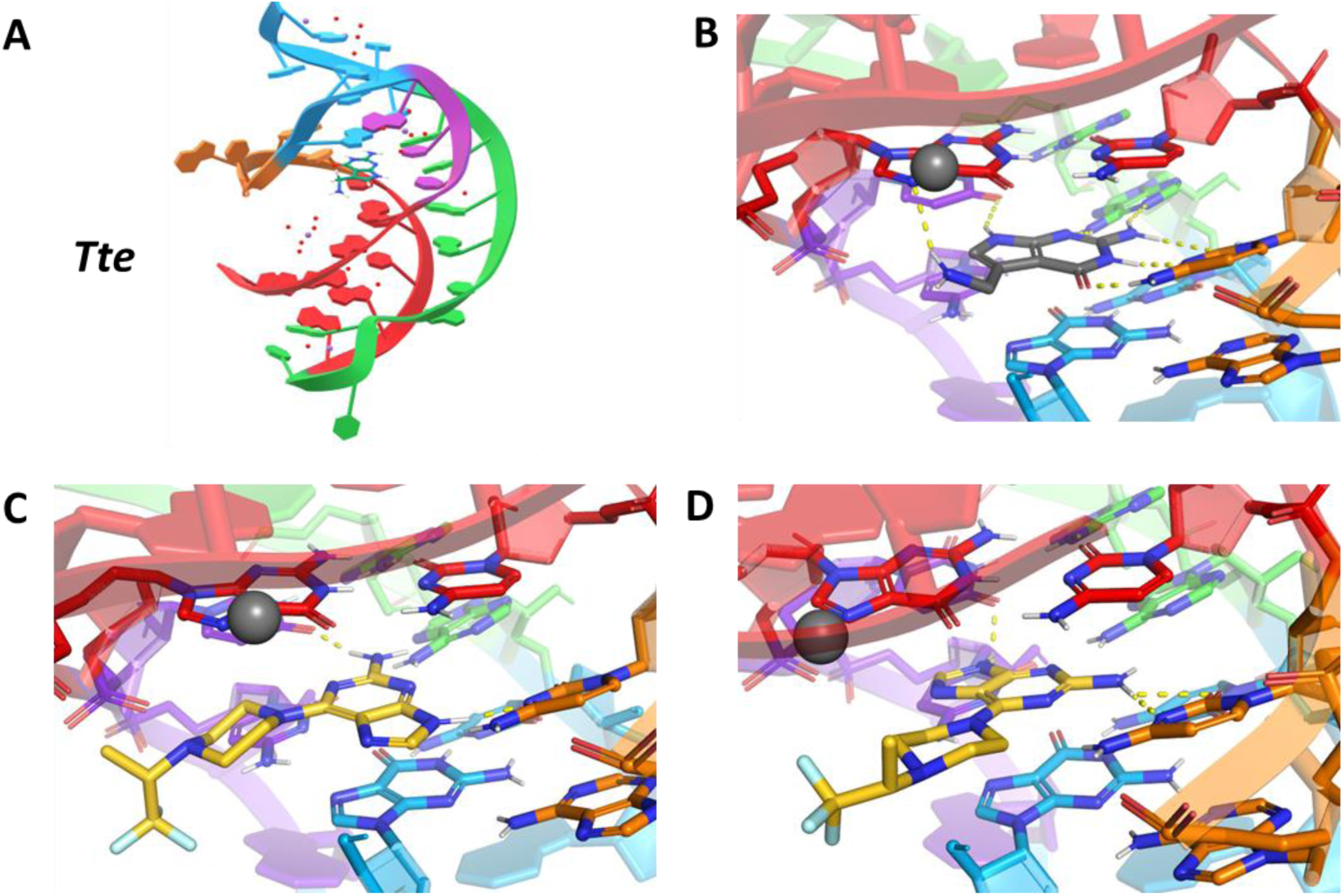
Docking of PreQ_1_ and **4494** to the *Thermoanaerobacter tengcongensis* (*Tte*) PreQ_1_ riboswitch (PDB ID: 6VUI). Most probable binding pose of **4494** was manually selected. Hydrogen bonds are indicated with yellow dashed lines. A) Crystal structure of the *Tte* PreQ_1_ riboswitch co-crystallized with PreQ_1_. B-D) Close up docked structure of PreQ_1_ (B) or **4494** (C and D) bound to the *Tte* PreQ_1_ riboswitch.

**Figure S17:**
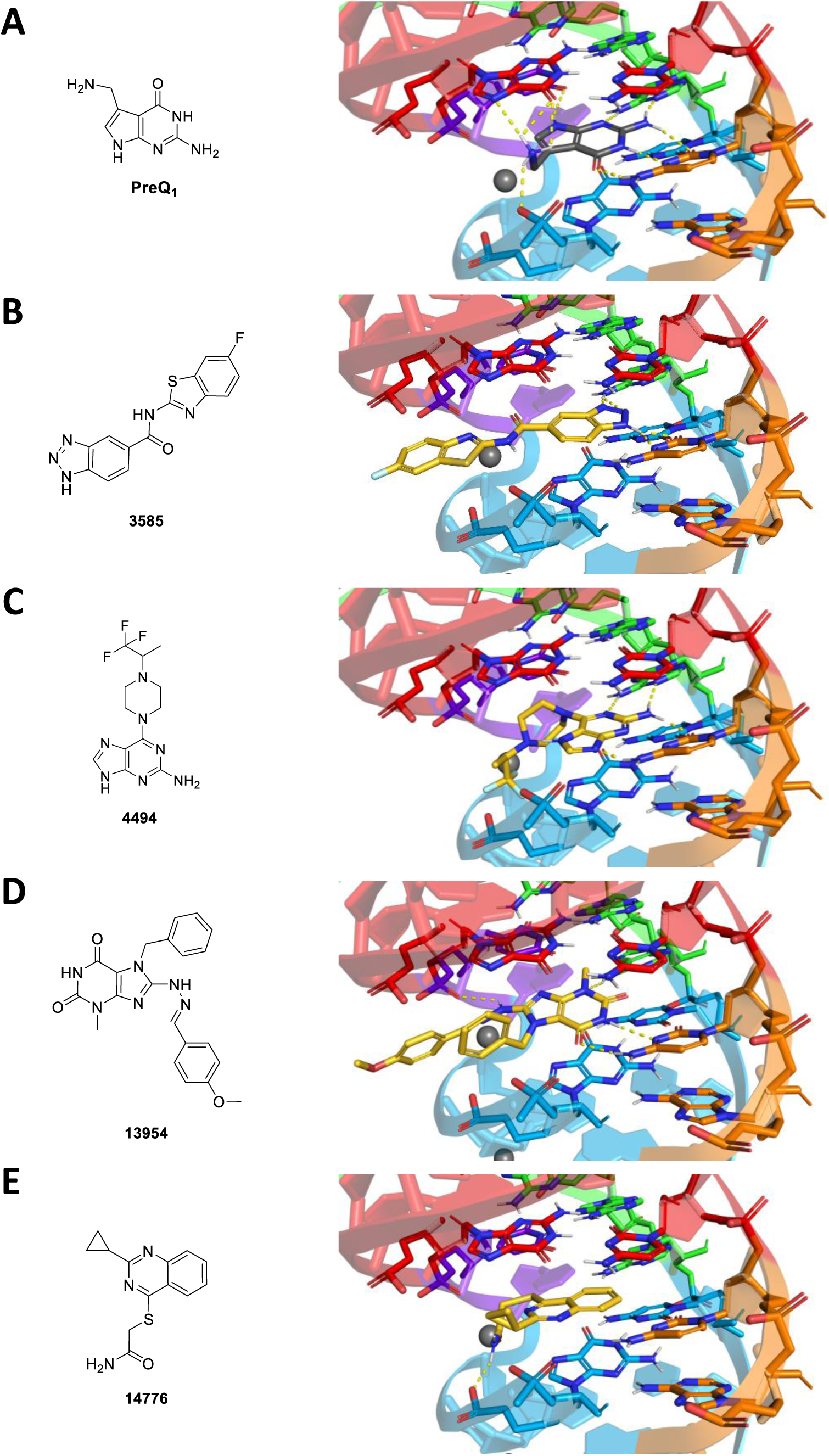
Molecular docking of hits to the *Bacillus subtilis* (*Bsu*) PreQ_1_ riboswitch (PBD: 3FU2). Most probable binding poses were manually selected. Hydrogen bonds are indicated with yellow dashed lines. **A**) PreQ_1_. **B**) **3585**. **C**) **4494**. **D**) **13954**. **E**) **14776**.

**Figure S18:**
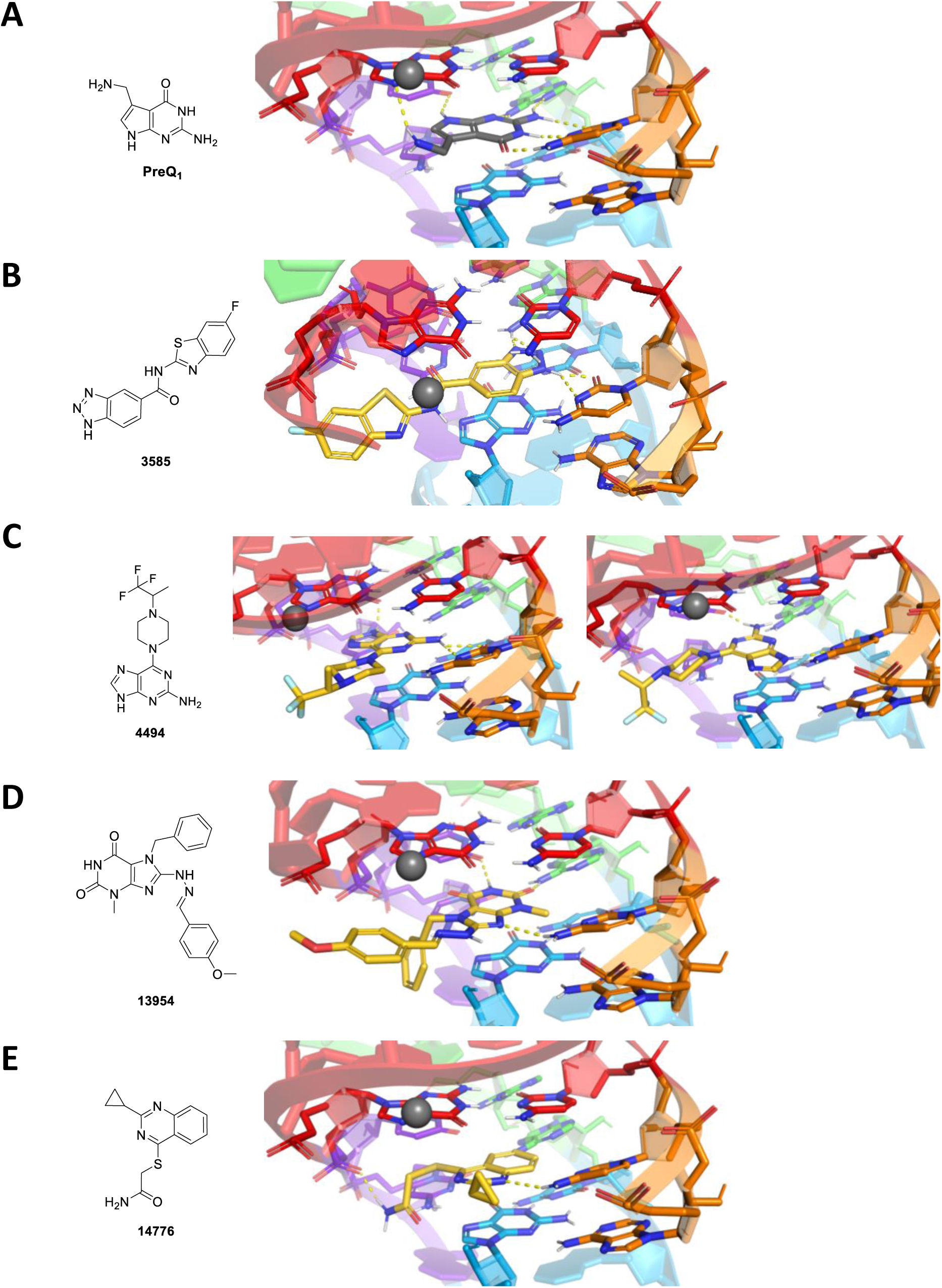
Molecular docking of hits to the *Thermoanaerobacter tengcongensis* (*Tte*) PreQ_1_ riboswitch (PBD: 6VUI). Most probable binding poses were manually selected. Hydrogen bonds are indicated with yellow dashed lines. **A**) PreQ_1_. **B**) **3585**. **C**) **4494**. **D**) **13954**. **E**) **14776**.

**Table S4:** Summary of HTS and hit validation data of promising hits. All data was normalized to the DMSO control for ease of comparison, and a color gradient indicates low (green) to high (red) values. HTS data from both screenings, hit confirmation 1 (100 µM) and hit confirmation 2 (5, 25, 50 and 100 µM) are shown. Blue compounds were selected for re-purchase (‘most promising hits’). Below the table, the workflow of hit selection and confirmation is shown.

| Hit | Data normalized to DMSO ctrl (% Cy5 signal) |  |  |  |  |  |  |  |  |  |  |  |  |  |
| --- | --- | --- | --- | --- | --- | --- | --- | --- | --- | --- | --- | --- | --- | --- |
| | HTS (25 μM) | | Hit confirmation 1 (100 μM) | | | | Hit confirmation 2 (5, 25, 50 and 100 μM) | | | | | | | |
|  | HTS 1 | HTS 2 | 1 | 2 | 3 | 4 | 0 | 5 | 5 | 25 | 25 | 50 | 50 | 100 |
| 1116 | 168 | 175 | 117 | 119 | 126 | 126 | 102 | 97 | 98 | 99 | 95 | 123 | 128 | 115 |
| 3506 | 97 | 130 | 98 | 144 |  |  | 102 | 104 | 102 | 557 | 99 | 101 | 100 | 98 |
| 3535 | 151 | 131 | 166 | 165 |  |  | 100 | 92 | 97 | 93 | 93 | 147 | 146 | 187 |
| 3585 | 170 | 110 | 164 | 163 | 181 | 165 | 100 | 102 | 99 | 100 | 102 | 108 | 106 | 180 |
| 3944 | 130 | 125 | 113 | 108 |  |  | 101 | 103 | 101 | 103 | 101 | 113 | 143 | 126 |
| 4316 | 141 | 122 | 126 | 104 |  |  | 97 | 101 | 99 | 84 | 99 | 99 | 104 | 116 |
| 4494 | 136 | 149 | 214 | 218 | 237 | 240 | 97 | 110 | 107 | 495 | 139 | 191 | 176 | 236 |
| 5065 | 191 | 182 | 139 | 149 | 159 | 111 | 101 | 97 | 99 | 94 | 89 | 93 | 93 | 290 |
| 7186 | 199 | 245 | 385 | 369 |  |  | 101 | 108 | 113 | 140 | 494 | 206 | 223 | 543 |
| 7239 | 82 | 621 | 133 | 182 | 189 | 147 | 100 | 100 | 97 | 96 | 99 | 94 | 89 | 205 |
| 7245 | 126 | 121 | 164 | 176 | 172 | 168 | 100 | 97 | 103 | 97 | 98 | 136 | 112 | 169 |
| 7426 | 202 | 137 | 110 | 186 | 201 | 175 | 100 | 96 | 97 | 96 | 97 | 101 | 97 | 244 |
| 11048 | 108 | 163 | 158 | 161 | 105 | 107 | 100 | 100 | 97 | 94 | 79 | 161 | 147 | 179 |
| 11229 | 104 | 135 | 257 | 268 |  |  | 99 | 96 | 96 | 93 | 96 | 106 | 121 | 254 |
| 12076 | 168 | 151 | 281 | 108 |  |  | 103 | 97 | 100 | 94 | 98 | 95 | 99 | 236 |
| 12576 | 132 | 131 | 125 | 128 |  |  | 104 | 109 | 109 | 111 | 114 | 114 | 119 | 124 |
| 13954 | 112 | 124 | 135 | 141 |  |  | 101 | 101 | 101 | 102 | 99 | 107 | 106 | 144 |
| 14085 | 155 | 134 | 104 | 107 | 121 | 114 | 98 | 101 | 99 | 97 | 98 | 103 | 102 | 117 |
| 14670 | 134 | 153 | 166 | 169 |  |  | 108 | 110 | 109 | 117 | 122 | 134 | 130 | 154 |
| 14776 | 153 | 156 | 16 | 26 |  |  | 101 | 116 | 116 | 182 | 186 | 213 | 202 | 73 |
| 15025 | 124 | 470 | 134 | 132 |  |  | 101 | 101 | 102 | 109 | 109 | 120 | 113 | 129 |

## NMR characterization of 4494

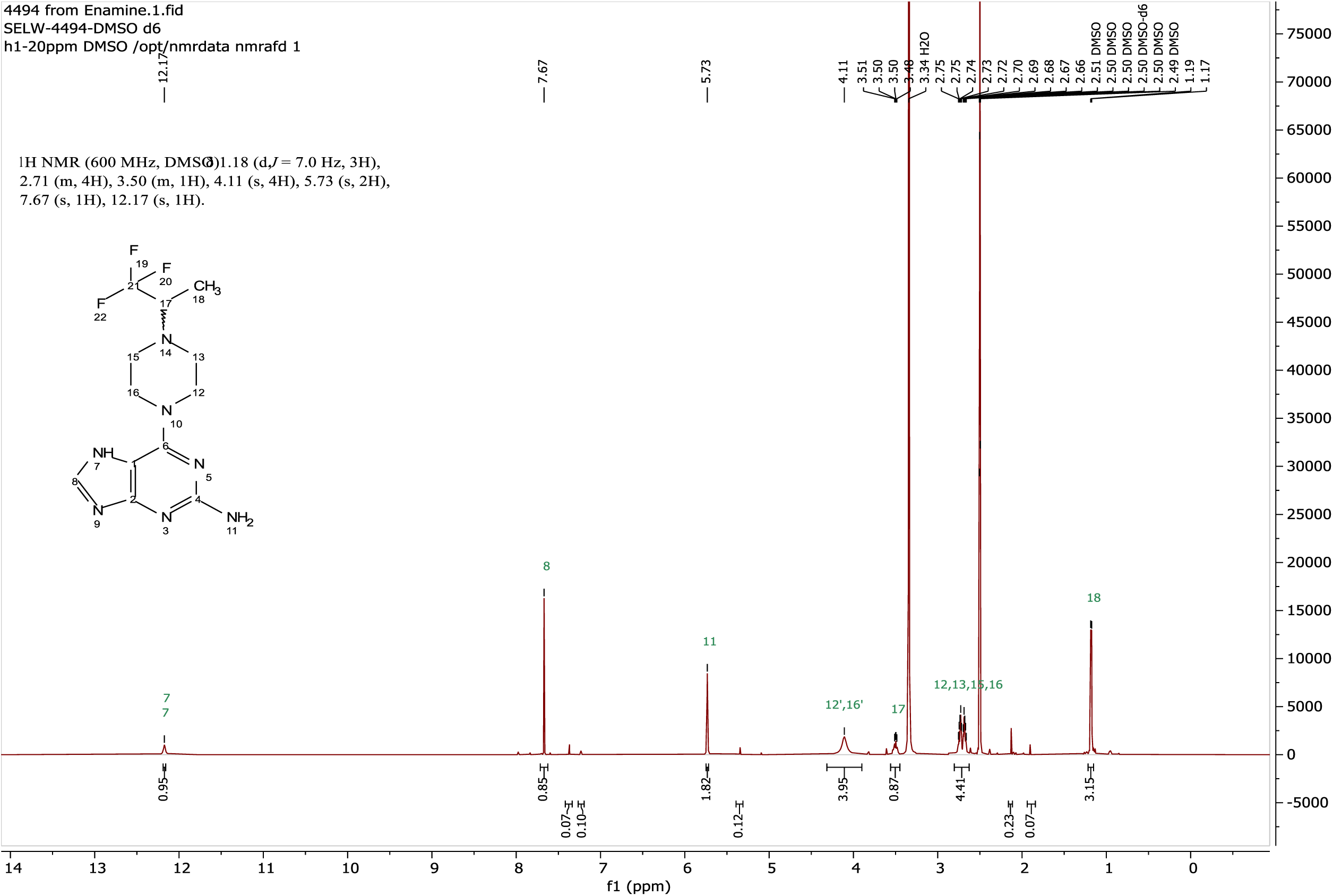

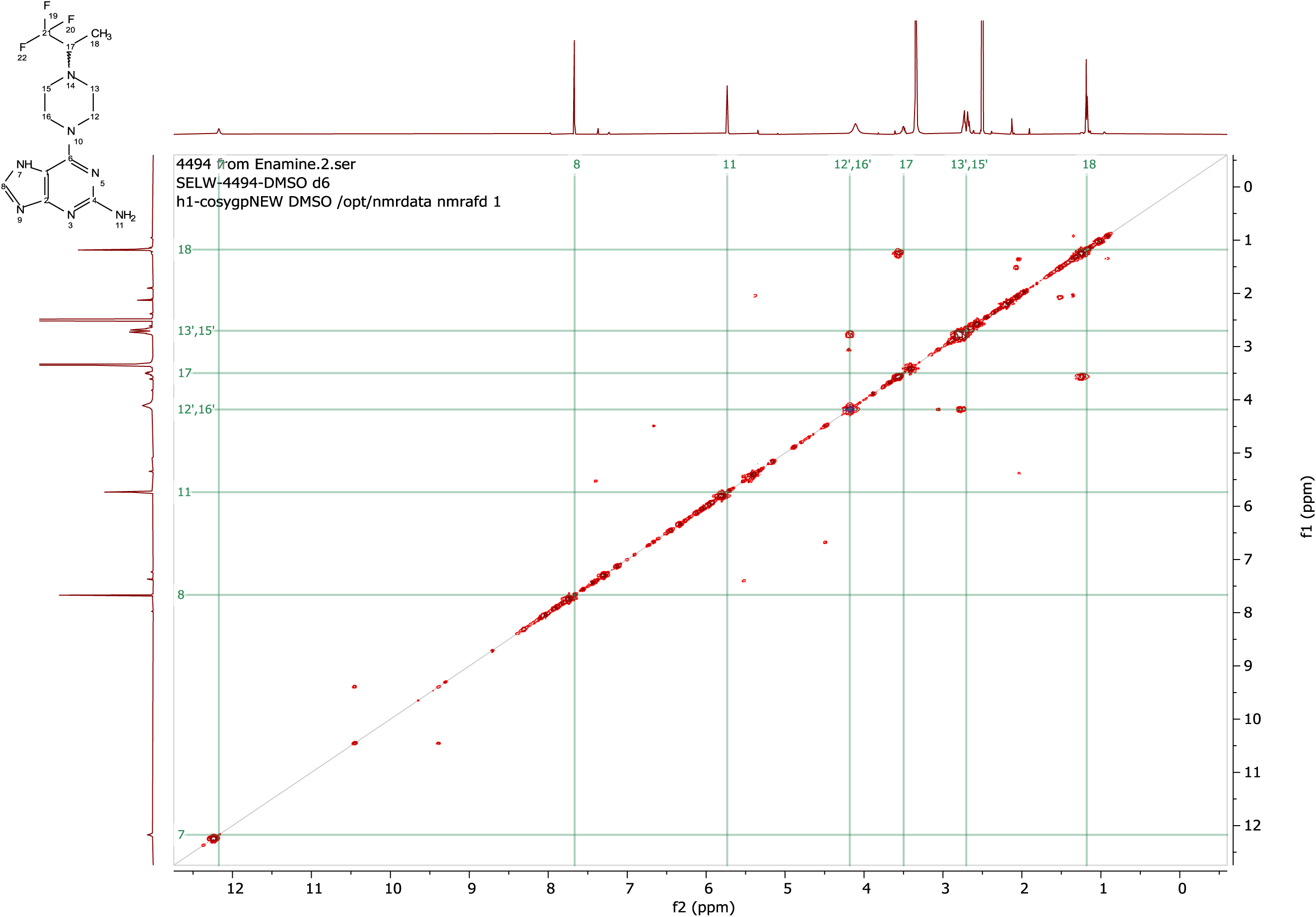

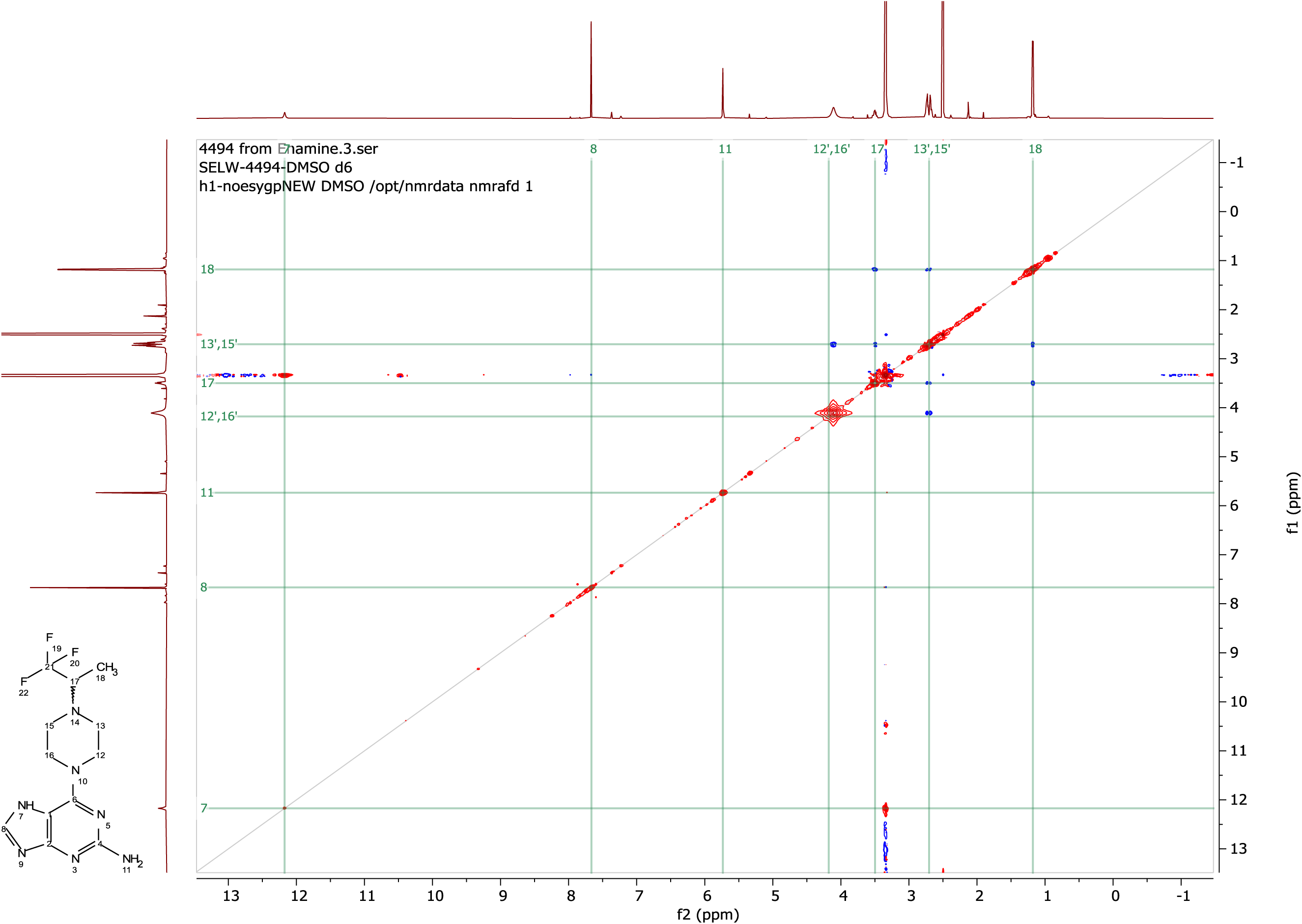

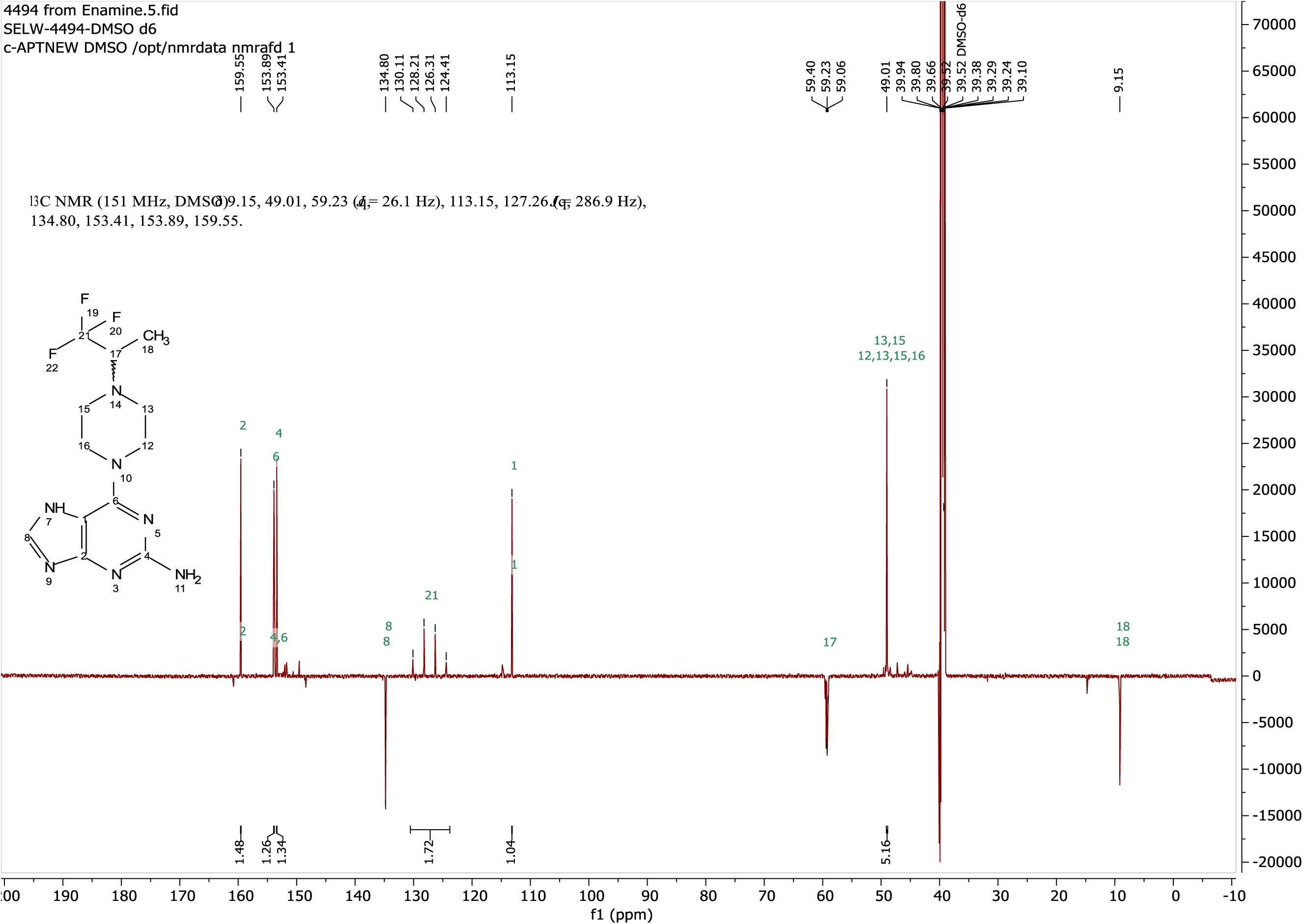

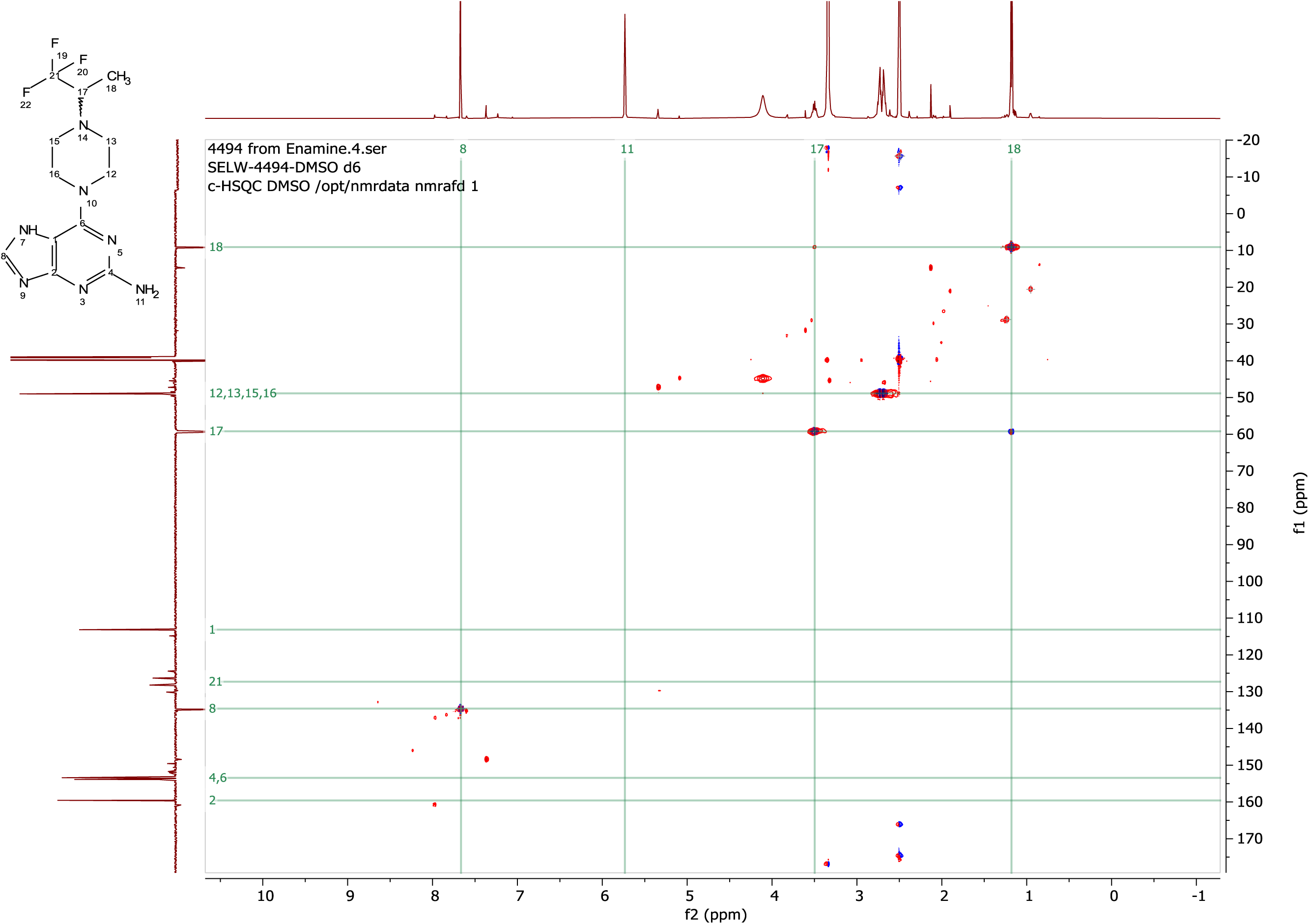

## Raw gels

### Gels Figure 6C

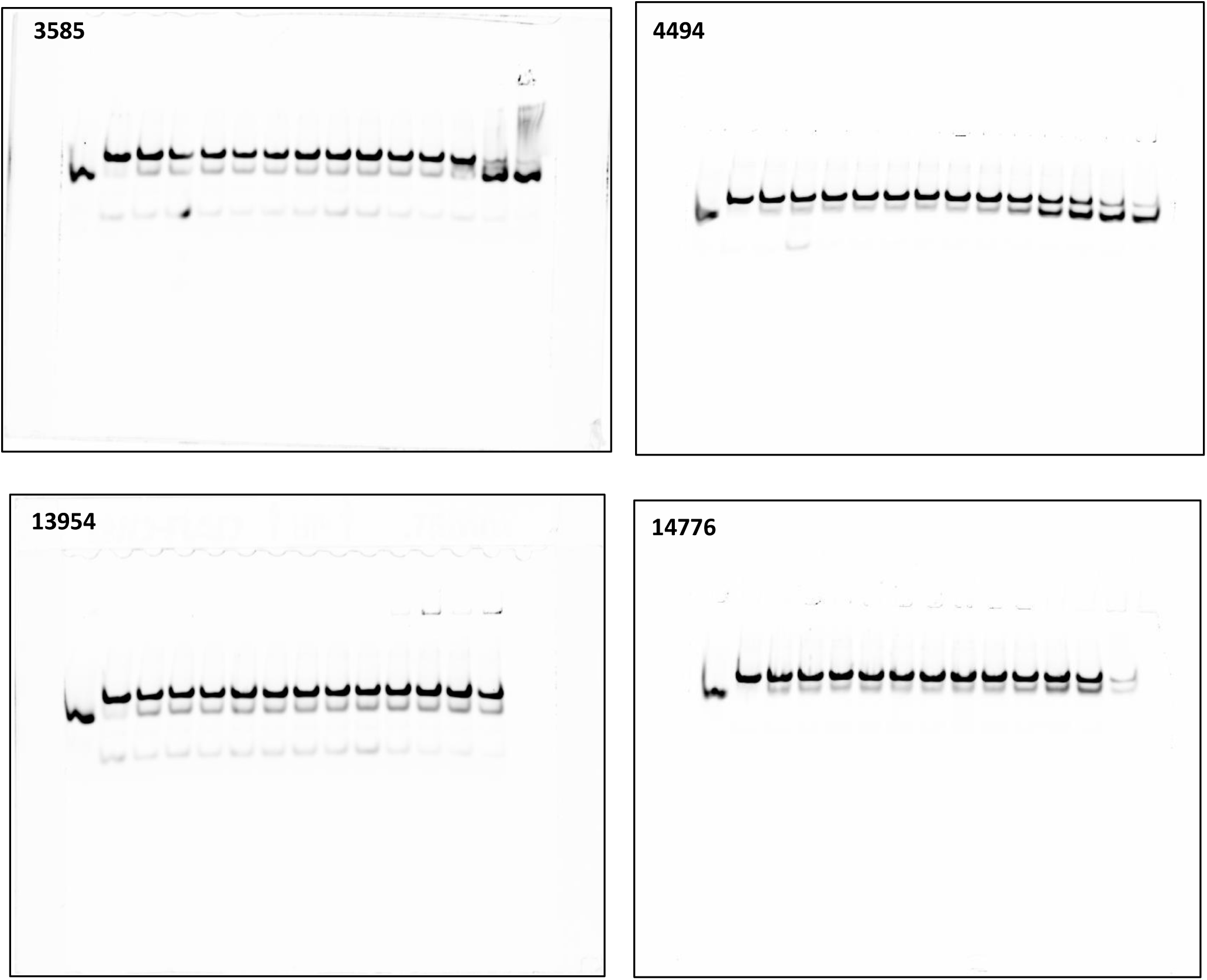

### Gels Figure S4

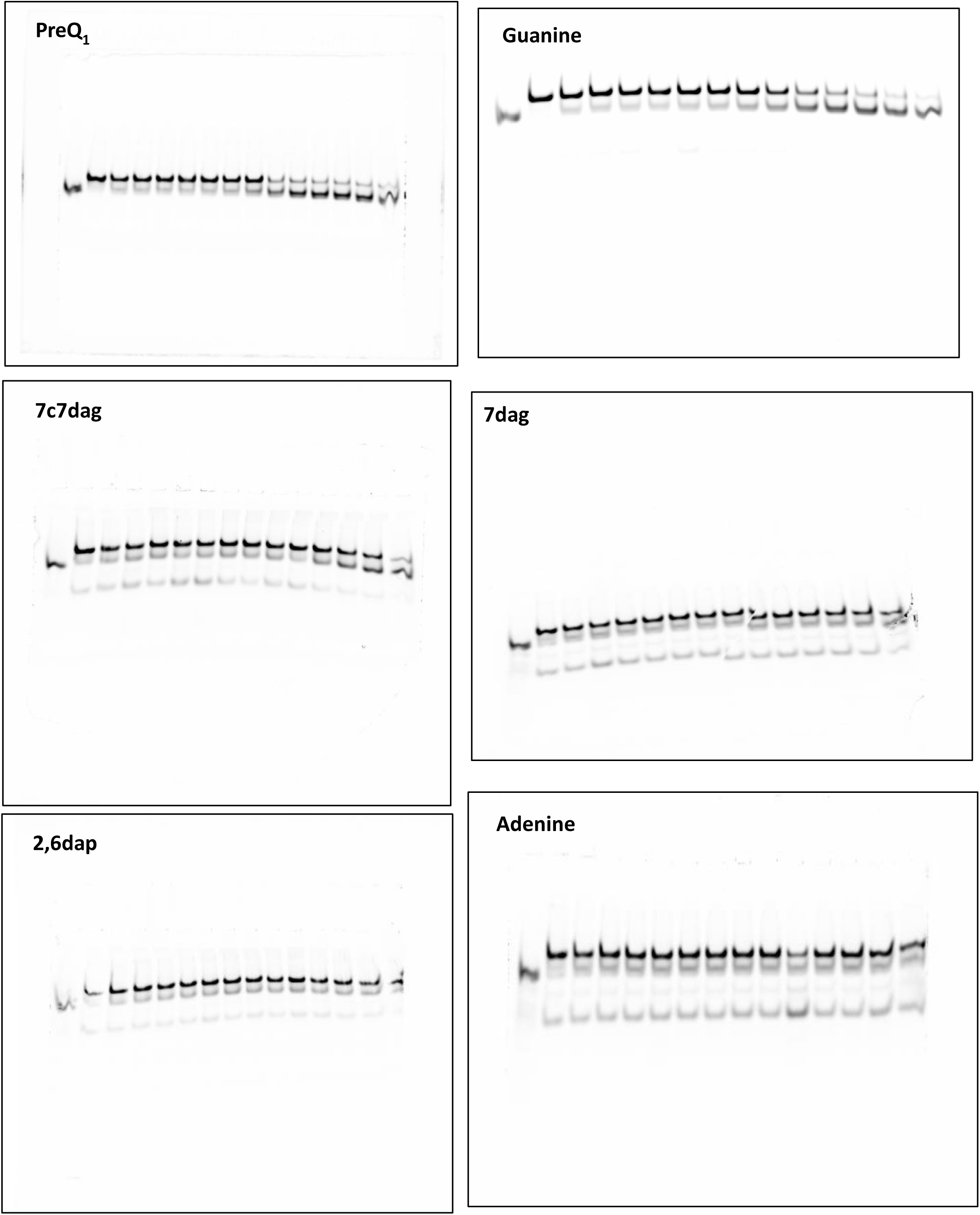

### Gels Figure S5C

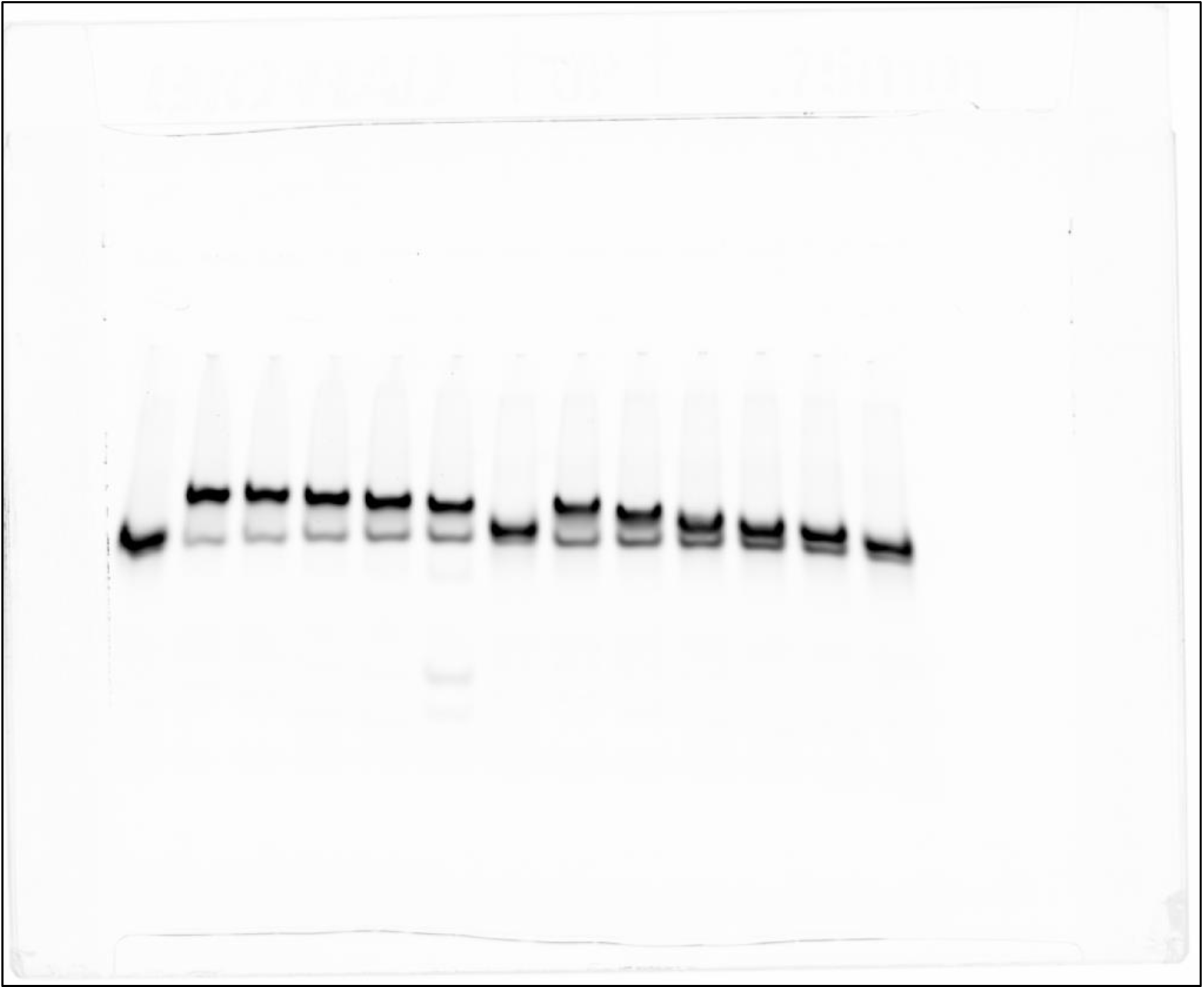

### Gels Figure S6C

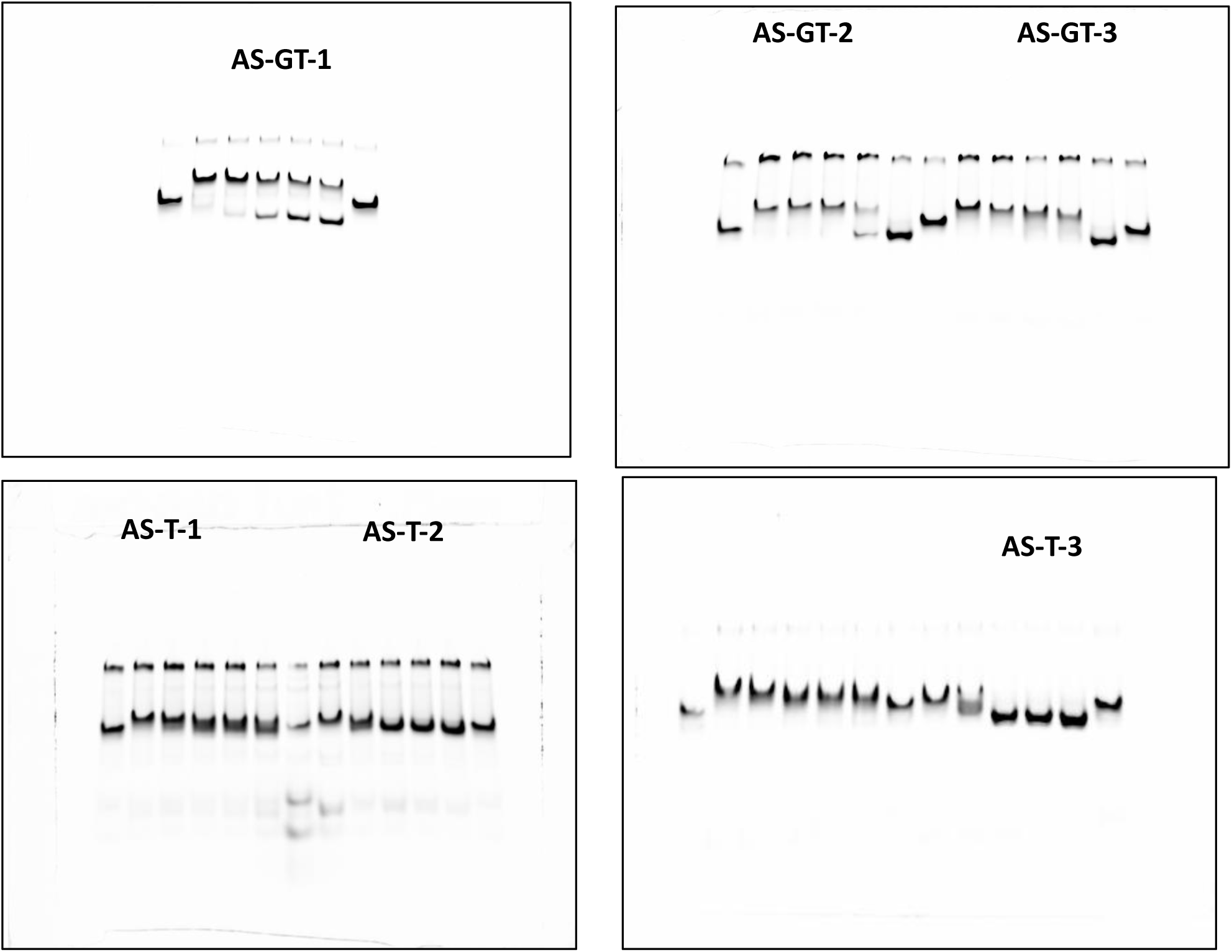

### Gels Figure S7C

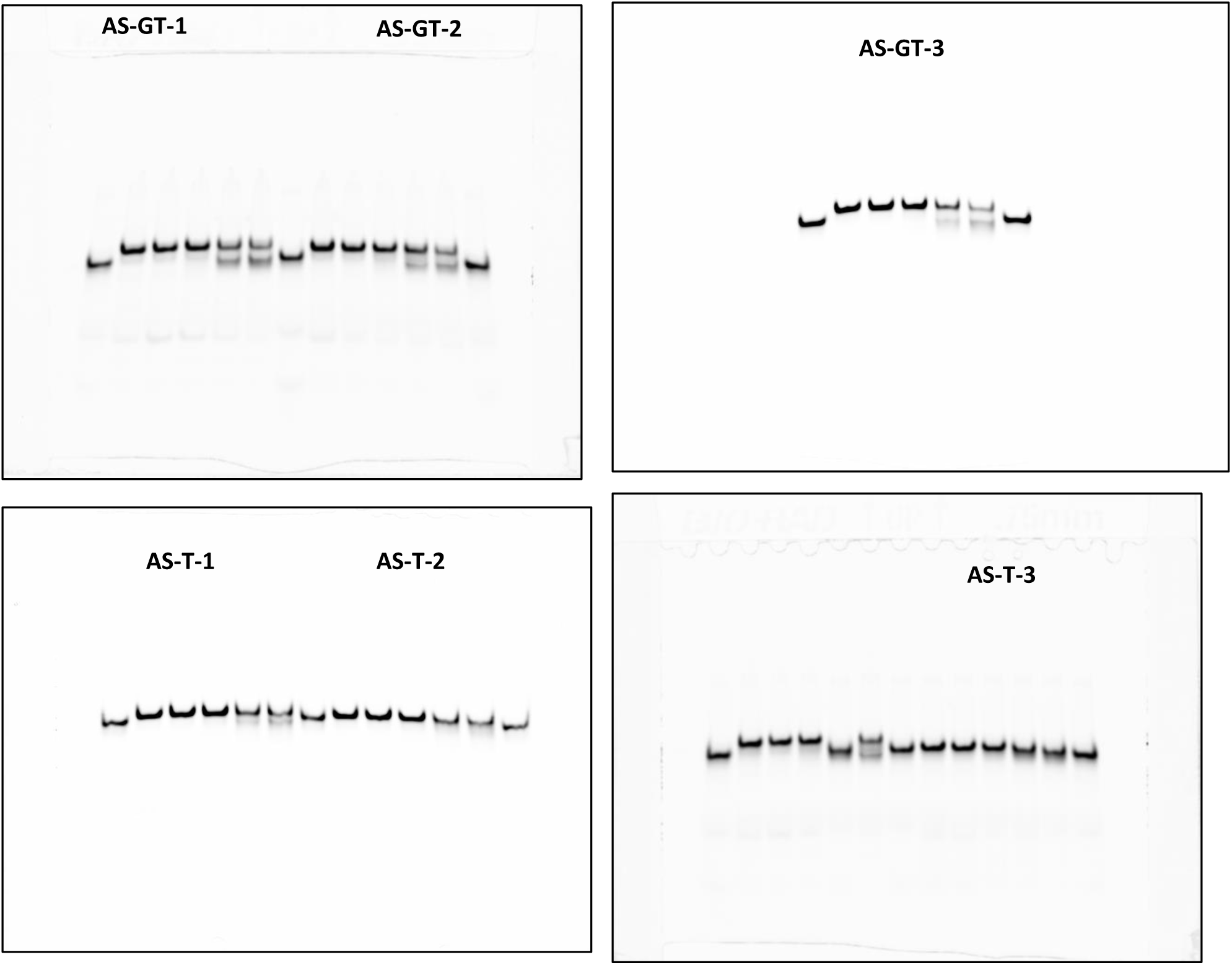

### Gels Figure S11

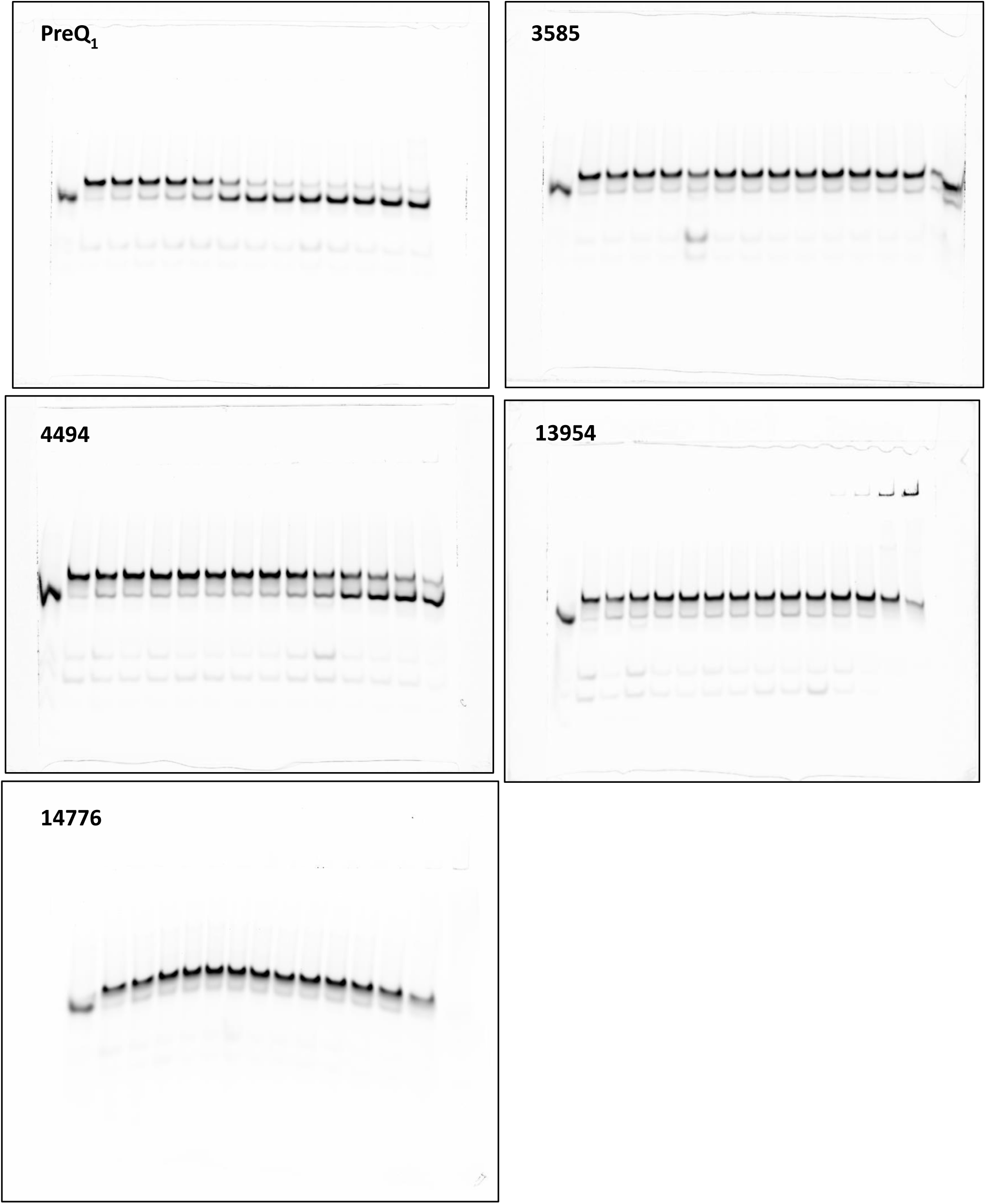

## References

(1) Lünse, C. E.; Schüller, A.; Mayer, G. The Promise of Riboswitches as Potential Antibacterial Drug Targets. Int. J. Med. Microbiol. 2014, 304 (1), 79–92. 10.1016/j.ijmm.2013.09.002.

(2) Mandal, M.; Breaker, R. R. Gene Regulation by Riboswitches. Nat. Rev. Mol. Cell Biol. 2004, 5 (6), 451–463. 10.1038/nrm1403.

(3) Tucker, B. J.; Breaker, R. R. Riboswitches as Versatile Gene Control Elements. Curr. Opin. Struct. Biol. 2005, 15 (3), 342–348. 10.1016/j.sbi.2005.05.003.

(4) Serganov, A.; Nudler, E. A Decade of Riboswitches. Cell 2013, 152 (1), 17–24. 10.1016/j.cell.2012.12.024.

(5) Garst, A. D.; Edwards, A. L.; Batey, R. T. Riboswitches: Structures and Mechanisms. Cold Spring Harb. Perspect. Biol. 2011, 3 (6), a003533. 10.1101/cshperspect.a003533.

(6) Nudler, E.; Mironov, A. S. The Riboswitch Control of Bacterial Metabolism. Trends Biochem. Sci. 2004, 29 (1), 11–17. 10.1016/j.tibs.2003.11.004.

(7) Blount, K. F.; Breaker, R. R. Riboswitches as Antibacterial Drug Targets. Nat. Biotechnol. 2006, 24 (12), 1558–1564. 10.1038/nbt1268.

(8) Roth, A.; Winkler, W. C.; Regulski, E. E.; Lee, B. W. K.; Lim, J.; Jona, I.; Barrick, J. E.; Ritwik, A.; Kim, J. N.; Welz, R.; Iwata-Reuyl, D.; Breaker, R. R. A Riboswitch Selective for the Queuosine Precursor preQ1 Contains an Unusually Small Aptamer Domain. Nat. Struct. Mol. Biol. 2007, 14 (4), 308–317. 10.1038/nsmb1224.

(9) McCown, P. J.; Liang, J. J.; Weinberg, Z.; Breaker, R. R. Structural, Functional, and Taxonomic Diversity of Three PreQ1 Riboswitch Classes. Chem. Biol. 2014, 21 (7), 880–889. 10.1016/j.chembiol.2014.05.015.

(10) Pavlova, N.; Penchovsky, R. Bioinformatics and Genomic Analyses of the Suitability of Eight Riboswitches for Antibacterial Drug Targets. Antibiotics 2022, 11 (9), 1177. 10.3390/an-tibiotics11091177.

(11) Pavlova, N.; Kaloudas, D.; Penchovsky, R. Riboswitch Distribution, Structure, and Function in Bacteria. Gene 2019, 708, 38–48. 10.1016/j.gene.2019.05.036.

(12) Harada, F.; Nishimura, S. Possible Anticodon Sequences of tRNAHis, tRNAAsn, and tRNAAsp from Escherichia Coli. Universal Presence of Nucleoside O in the First Position of the Anticodons of These Transfer Ribonucleic Acid. Biochemistry 1972, 11 (2), 301–308. 10.1021/bi00752a024.

(13) Hillmeier, M.; Wagner, M.; Ensfelder, T.; Korytiakova, E.; Thumbs, P.; Müller, M.; Carell, T. Synthesis and Structure Elucidation of the Human tRNA Nucleoside Mannosyl-Queuosine. Nat. Commun. 2021, 12 (1), 7123. 10.1038/s41467-021-27371-9.

(14) Iwata-Reuyl, D. Biosynthesis of the 7-Deazaguanosine Hypermodified Nucleosides of Transfer RNA. Bioorganic Chem. 2003, 31 (1), 24–43. 10.1016/S0045-2068(02)00513-8.

(15) Meier, F.; Suter, B.; Grosjean, H.; Keith, G.; Kubli, E. Queuosine Modification of the Wobble Base in tRNAHis Influences ‘in Vivo’ Decoding Properties. EMBO J. 1985, 4 (3), 823–827. 10.1002/j.1460-2075.1985.tb03704.x.

(16) Durand, J. M. B.; Dagberg, B.; Uhlin, B. E.; Björk, G. R. Transfer RNA Modification, Temperature and DNA Superhelicity Have a Common Target in the Regulatory Network of the Virulence of Shigella Flexneri: The Expression of the virF Gene. Mol. Microbiol. 2000, 35 (4), 924–935. 10.1046/j.1365-2958.2000.01767.x.

(17) Noguchi, S.; Nishimura, Y.; Hirota, Y.; Nishimura, S. Isolation and Characterization of an Escherichia Coli Mutant Lacking tRNA-Guanine Transglycosylase. Function and Biosynthesis of Queuosine in tRNA. J. Biol. Chem. 1982, 257 (11), 6544–6550. 10.1016/S0021-9258(20)65176-6.

(18) Eichhorn, C. D.; Kang, M.; Feigon, J. Structure and Function of preQ1 Riboswitches. Biochim. Biophys. Acta BBA - Gene Regul. Mech. 2014, 1839 (10), 939–950. 10.1016/j.bbagrm.2014.04.019.

(19) McCarty, R. M.; Bandarian, V. Biosynthesis of Pyrrolopyrimidines. Bioorganic Chem. 2012, 43, 15–25. 10.1016/j.bioorg.2012.01.001.

(20) Meyer, M. M.; Roth, A.; Chervin, S. M.; Garcia, G. A.; Breaker, R. R. Confirmation of a Second Natural preQ1 Aptamer Class in Streptococcaceae Bacteria. RNA 2008, 14 (4), 685–695. 10.1261/rna.937308.

(21) Kim, J. N.; Breaker, R. R. Purine Sensing by Riboswitches. Biol. Cell 2008, 100 (1), 1–11. 10.1042/BC20070088.

(22) Pleij, C. W. A.; Rietveld, K.; Bosch, L. A New Principle of RNA Folding Based on Pseudoknotting. Nucleic Acids Res. 1985, 13 (5), 1717–1731. 10.1093/nar/13.5.1717.

(23) Klein, D. J.; Edwards, T. E.; Ferré-D’Amaré, A. R. Cocrystal Structure of a Class I preQ1 Riboswitch Rveals a Pseudoknot Recognizing an Essential Hypermodified Nucleobase. Nat. Struct. Mol. Biol. 2009, 16 (3), 343–344. 10.1038/nsmb.1563.

(24) Rieder, U.; Lang, K.; Kreutz, C.; Polacek, N.; Micura, R. Evidence for Pseudoknot Formation of Class I preQ1 Riboswitch Aptamers. ChemBioChem 2009, 10 (7), 1141–1144. 10.1002/cbic.200900155.

(25) Jenkins, J. L.; Krucinska, J.; McCarty, R. M.; Bandarian, V.; Wedekind, J. E. Comparison of a PreQ1 Riboswitch Aptamer in Metabolite-Bound and Free States with Implications for Gene Regulation *. J. Biol. Chem. 2011, 286 (28), 24626–24637. 10.1074/jbc.M111.230375.

(26) Suddala, K. C.; Rinaldi, A. J.; Feng, J.; Mustoe, A. M.; Eichhorn, C. D.; Liberman, J. A.; Wedekind, J. E.; Al-Hashimi, H. M.; Brooks, C. L.; Walter, N. G. Single Transcriptional and Translational preQ1 Riboswitches Adopt Similar Pre-Folded Ensembles That Follow Distinct Folding Pathways into the Same Ligand-Bound Structure. Nucleic Acids Res. 2013, 41 (22), 10462–10475. 10.1093/nar/gkt798.

(27) Connelly, C. M.; Numata, T.; Boer, R. E.; Moon, M. H.; Sinniah, R. S.; Barchi, J. J.; Ferré-D’Amaré, A. R.; Schneekloth, J. S. Synthetic Ligands for PreQ1 Riboswitches Provide Structural and Mechanistic Insights into Targeting RNA Tertiary Structure. Nat. Commun. 2019, 10 (1), 1501. 10.1038/s41467-019-09493-3.

(28) Neuner, E.; Frener, M.; Lusser, A.; Micura, R. Superior Cellular Activities of Azido-over Amino-Functionalized Ligands for Engineered preQ1 Riboswitches in E.Coli. RNA Biol. 2018, 15 (10), 1376–1383. 10.1080/15476286.2018.1534526.

(29) Kallert, E.; Fischer, T. R.; Schneider, S.; Grimm, M.; Helm, M.; Kersten, C. Protein-Based Virtual Screening Tools Applied for RNA–Ligand Docking Identify New Binders of the preQ1-Riboswitch. J. Chem. Inf. Model. 2022, 62 (17), 4134–4148. 10.1021/acs.jcim.2c00751.

(30) Kallert, E.; Fischer, T. R.; Schneider, S.; Grimm, M.; Helm, M.; Kersten, C. *Protein-Based Virtual Screening Tools Applied for RNA-Ligand Docking Identify New Binders of the preQ _1_* -Riboswitch; preprint; Bioinformatics, 2022. 10.1101/2022.06.10.494309.

(31) Wicks, S. L.; Hargrove, A. E. Fluorescent Indicator Displacement Assays to Identify and Characterize Small Molecule Interactions with RNA. Methods 2019, 167, 3–14. 10.1016/j.ymeth.2019.04.018.

(32) Mei, H.-Y.; Mack, D. P.; Galan, A. A.; Halim, N. S.; Heldsinger, A.; Loo, J. A.; Moreland, D. W.; Sannes-Lowery, K. A.; Sharmeen, L.; Truong, H. N.; Czarnik, A. W. Discovery of Selective, Small-Molecule Inhibitors of RNA Complexes—1. The Tat Protein/TAR RNA Complexes Required for HIV-1 Transcription. Bioorg. Med. Chem. 1997, 5 (6), 1173–1184. 10.1016/S0968-0896(97)00064-3.

(33) Gelus, N.; Bailly, C.; Hamy, F.; Klimkait, T.; Wilson, W. D.; Boykin, D. W. Inhibition of HIV-1 Tat-TAR Interaction by Diphenylfuran Derivatives: Effects of the Terminal Basic Side Chains. Bioorg. Med. Chem. 1999, 7 (6), 1089–1096. 10.1016/S0968-0896(99)00041-3.

(34) Patwardhan, N. N.; Cai, Z.; Newson, C. N.; Hargrove, A. E. Fluorescent Peptide Displacement as a General Assay for Screening Small Molecule Libraries against RNA. Org. Biomol. Chem. 2019, 17 (7), 1778–1786. 10.1039/C8OB02467G.

(35) Ganser, L. R.; Lee, J.; Rangadurai, A.; Merriman, D. K.; Kelly, M. L.; Kansal, A. D.; Sathyamoorthy, B.; Al-Hashimi, H. M. High-Performance Virtual Screening by Targeting a High-Resolution RNA Dynamic Ensemble. Nat. Struct. Mol. Biol. 2018, 25 (5), 425–434. 10.1038/s41594-018-0062-4.

(36) Menichelli, E.; Lam, B. J.; Wang, Y.; Wang, V. S.; Shaffer, J.; Tjhung, K. F.; Bursulaya, B.; Nguyen, T. N.; Vo, T.; Alper, P. B.; McAllister, C. S.; Jones, D. H.; Spraggon, G.; Michellys, P.-Y.; Joslin, J.; Joyce, G. F.; Rogers, J. Discovery of Small Molecules That Target a Tertiary-Structured RNA. Proc. Natl. Acad. Sci. 2022, 119 (48), e2213117119. 10.1073/pnas.2213117119.

(37) Rieder, U.; Kreutz, C.; Micura, R. Folding of a Transcriptionally Acting PreQ1 Riboswitch. Proc. Natl. Acad. Sci. 2010, 107 (24), 10804–10809. 10.1073/pnas.0914925107.

(38) Yu, C.-H.; Luo, J.; Iwata-Reuyl, D.; Olsthoorn, R. C. L. Exploiting preQ1 Riboswitches To Regulate Ribosomal Frameshifting. ACS Chem. Biol. 2013, 8 (4), 733–740. 10.1021/cb300629b.

(39) Chen, Y.; Huang, Z.; Tang, Z.; Huang, Y.; Huang, M.; Liu, H.; Ziebolz, D.; Schmalz, G.; Jia, B.; Zhao, J. More Than Just a Periodontal Pathogen –the Research Progress on Fusobacterium Nucleatum. Front. Cell. Infect. Microbiol. 2022, 12, 815318. 10.3389/fcimb.2022.815318.

(40) Leamy, K. A.; Assmann, S. M.; Mathews, D. H.; Bevilacqua, P. C. Bridging the Gap between in Vitro and in Vivo RNA Folding. Q. Rev. Biophys. 2016, 49. 10.1017/S003358351600007X.

(41) Feig, A. L.; Uhlenbeck, O. C. The Role of Metal Ions in RNA Biochemistry. Cold Spring Harb. Monogr. Ser. 1999, 37, 287–320.

(42) Braasch, D. A.; Corey, D. R. Locked Nucleic Acid (LNA): Fine-Tuning the Recognition of DNA and RNA. Chem. Biol. 2001, 8 (1), 1–7. 10.1016/S1074-5521(00)00058-2.

(43) Malo, N.; Hanley, J. A.; Cerquozzi, S.; Pelletier, J.; Nadon, R. Statistical Practice in High-Throughput Screening Data Analysis. Nat. Biotechnol. 2006, 24 (2), 167–175. 10.1038/nbt1186.

(44) Turek-Etienne, T. C.; Small, E. C.; Soh, S. C.; Xin, T. A.; Gaitonde, P. V.; Barrabee, E. B.; Hart, R. F.; Bryant, R. W. Evaluation of Fluorescent Compound Interference in 4 Fluorescence Polarization Assays: 2 Kinases, 1 Protease, and 1 Phosphatase. SLAS Discov. 2003, 8 (2), 176–184. 10.1177/1087057103252304.

(45) Lipinski, C. A.; Lombardo, F.; Dominy, B. W.; Feeney, P. J. Experimental and Computational Approaches to Estimate Solubility and Permeability in Drug Discovery and Development Settings. Adv. Drug Deliv. Rev. 2001, 46 (1–3), 3–26. 10.1016/s0169-409x(00)00129-0.

(46) Martin, W. J.; Grandi, P.; Marcia, M. Screening Strategies for Identifying RNA- and Ribonucleoprotein-Targeted Compounds. Trends Pharmacol. Sci. 2021, 42 (9), 758–771. 10.1016/j.tips.2021.06.001.

(47) Balaratnam, S.; Rhodes, C.; Bume, D. D.; Connelly, C.; Lai, C. C.; Kelley, J. A.; Yazdani, K.; Homan, P. J.; Incarnato, D.; Numata, T.; Schneekloth Jr, J. S. A Chemical Probe Based on the PreQ1 Metabolite Enables Transcriptome-Wide Mapping of Binding Sites. Nat. Commun. 2021, 12 (1), 5856. 10.1038/s41467-021-25973-x.

(48) Bessler, L.; Kaur, N.; Vogt, L.-M.; Flemmich, L.; Siebenaller, C.; Winz, M.-L.; Tuorto, F.; Micura, R.; Ehrenhofer-Murray, A. E.; Helm, M. Functional Integration of a Semi-Synthetic Azido-Queuosine Derivative into Translation and a tRNA Modification Circuit. Nucleic Acids Res. 2022, 50 (18), 10785–10800. 10.1093/nar/gkac822.

(49) Chen, C. Z.; Sobczak, K.; Hoskins, J.; Southall, N.; Marugan, J. J.; Zheng, W.; Thornton, C. A.; Austin, C. P. Two High Throughput Screening Assays for Aberrant RNA-Protein Interactions in Myotonic Dystrophy Type-1. Anal. Bioanal. Chem. 2012, 402 (5), 1889–1898. 10.1007/s00216-011-5604-0.

(50) Disney, M. D.; Labuda, L. P.; Paul, D. J.; Poplawski, S. G.; Pushechnikov, A.; Tran, T.; Velagapudi, S. P.; Wu, M.; Childs-Disney, J. L. Two-Dimensional Combinatorial Screening Identifies Specific Aminoglycoside−RNA Internal Loop Partners. J. Am. Chem. Soc. 2008, 130 (33), 11185–11194. 10.1021/ja803234t.

(51) Disney, M. D.; Winkelsas, A. M.; Velagapudi, S. P.; Southern, M.; Fallahi, M.; Childs-Disney, J. L. Inforna 2.0: A Platform for the Sequence-Based Design of Small Molecules Targeting Structured RNAs. ACS Chem. Biol. 2016, 11 (6), 1720–1728. 10.1021/acschembio.6b00001.

(52) Costales, M. G.; Matsumoto, Y.; Velagapudi, S. P.; Disney, M. D. Small Molecule Targeted Recruitment of a Nuclease to RNA. J. Am. Chem. Soc. 2018, 140 (22), 6741–6744. 10.1021/jacs.8b01233.

(53) Haniff, H. S.; Tong, Y.; Liu, X.; Chen, J. L.; Suresh, B. M.; Andrews, R. J.; Peterson, J. M.; O’Leary, C. A.; Benhamou, R. I.; Moss, W. N.; Disney, M. D. Targeting the SARS-CoV-2 RNA Genome with Small Molecule Binders and Ribonuclease Targeting Chimera (RIBOTAC) Degraders. ACS Cent. Sci. 2020, 6 (10), 1713–1721. 10.1021/acscentsci.0c00984.

(54) Mikutis, S.; Rebelo, M.; Yankova, E.; Gu, M.; Tang, C.; Coelho, A. R.; Yang, M.; Hazemi, M. E.; Pires de Miranda, M.; Eleftheriou, M.; Robertson, M.; Vassiliou, G. S.; Adams, D. J.; Simas, J. P.; Corzana, F.; Schneekloth, J. S. Jr.; Tzelepis, K.; Bernardes, G. J. L. Proximity-Induced Nucleic Acid Degrader (PINAD) Approach to Targeted RNA Degradation Using Small Molecules. ACS Cent. Sci. 2023, 9 (5), 892–904. 10.1021/acscentsci.3c00015.

## References

(1) Dilweg, I. W.; Oskam, M. G.; Overbeek, S.; Olsthoorn, R. C. L. Investigating the Correlation between Xrn1-Resistant RNAs and Frameshifter Pseudoknots. RNA Biol. 2023, 20 (1), 409–418. 10.1080/15476286.2023.2205224.

(2) Yu, C.-H.; Luo, J.; Iwata-Reuyl, D.; Olsthoorn, R. C. L. Exploiting preQ1 Riboswitches To Regulate Ribosomal Frameshifting. ACS Chem. Biol. 2013, 8 (4), 733–740. 10.1021/cb300629b.

